# RNA-Focused DNA-Encoded Library Construction, Screening, and Integration of Docking Identify Bioactive Ligands of Pathogenic r(G_4_C_2_)^exp^ RNA

**DOI:** 10.1101/2025.11.06.686979

**Authors:** Xueyi Yang, Amirhossein Taghavi, Yoshihiro Akahori, Martina Pedrini, Takahiro Ishii, Matthew D. Disney

## Abstract

Disease-associated RNAs are increasingly recognized as promising therapeutic targets for small-molecule intervention. While DNA-encoded libraries (DELs) have long been established for protein ligand discovery, recent studies have demonstrated their feasibility for identifying RNA-binding small molecules. To further advance RNA-targeted ligand discovery, a diverse, solid-phase DEL enriched in privileged RNA-binding scaffolds was constructed and applied to identify ligands of r(G_4_C_2_)^exp^, a toxic RNA repeat expansion implicated in amyotrophic lateral sclerosis (ALS) and frontotemporal dementia (FTD). DEL selection outcomes were analyzed through large-scale molecular docking integrated with physicochemical and structure-activity relationship (SAR) analyses. Strong correlations were observed between docking predictions and experimental enrichment trends, supporting lead identification. The lead compound was subsequently optimized based on its docked pose to an NMR structure, resulting in analogs with enhanced binding affinity and bioactivity. These findings demonstrate that RNA ligand identification can be effectively achieved by combining DNA-encoded library technology with computational approaches for rational design and analysis, and highlight a broadly adaptable platform for RNA-targeted small molecule discovery.

**TOC graphic:** 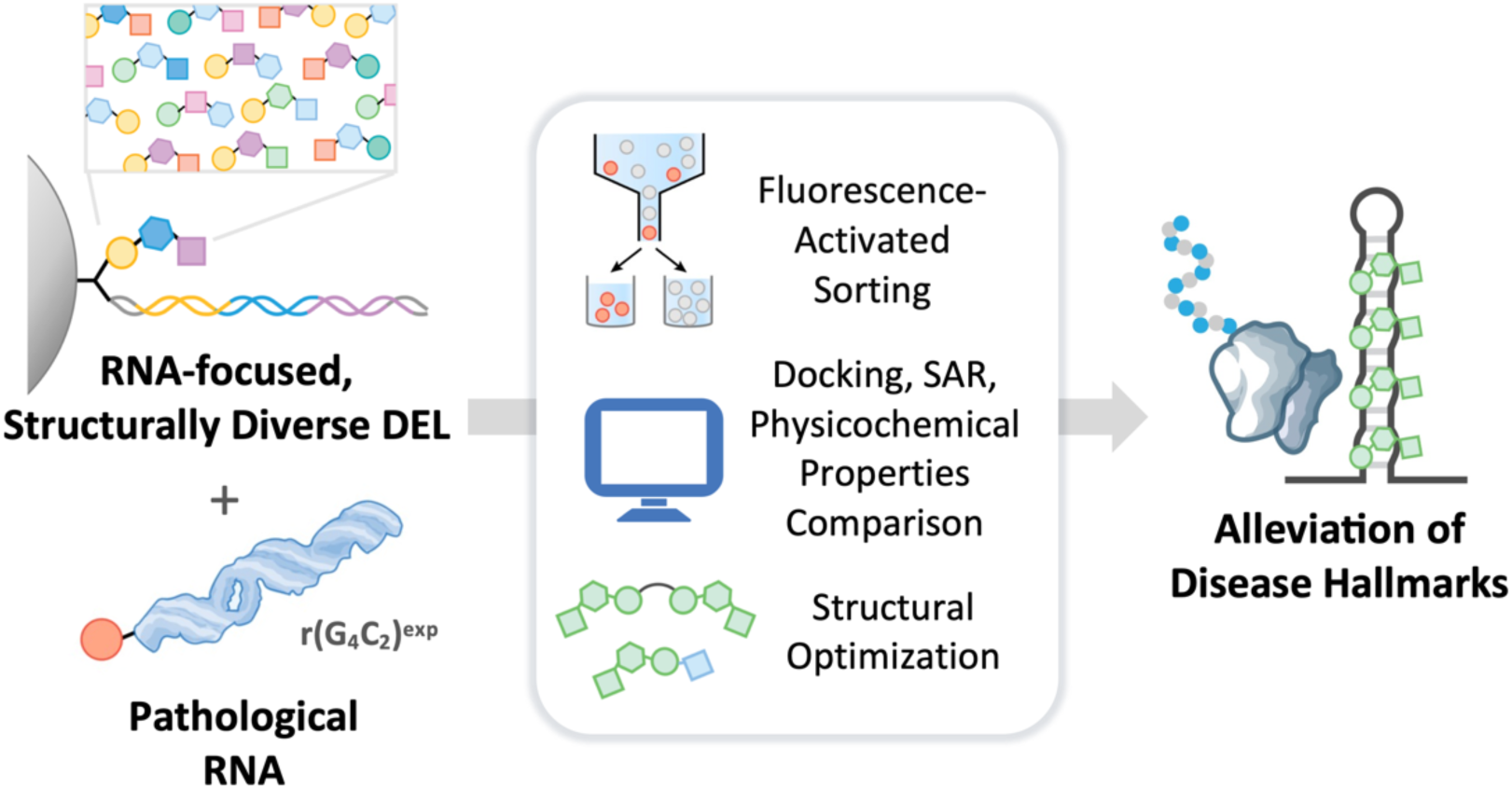

## Introduction

RNA is an attractive therapeutic target given its essential role in regulating cellular processes and disease progression. Historically, antisense oligonucleotides have been utilized to bind and/or degrade unstructured RNA sequence via an RNase H mediated pathway.^1^ Complementarily, small molecules can engage structured RNA folds, thereby modulating RNA function in various biological processes.^2–5^ Developing high-throughput screening methods to identify RNA ligands is an important goal in the field, yet broadly applicable approaches are still limited. DNA encoded library (DEL) technology, a simple, cost-effective and information-rich method, has become a favorable approach for ligand identification.^6–9^ A DNA encoded library contains a library of small molecules with identity encoded in conjugated DNA sequences. Post affinity enrichment, the DNA barcodes can be rapidly decoded by next-generation sequencing (NGS), enabling hit identification, followed by compound resynthesis and validation.^10^ The DEL technology has been utilized extensively in identifying protein ligands, with the targets including purified proteins,^7, 11–15^ protein/RNA degradation modulators,^16–18^ and protein in or on live cells.^19–22^

With increasing interest in small molecule RNA chemical probes and lead therapeutics, several studies have applied DEL technology to RNA ligand discovery, underscoring its feasibility and potential.^23–28^ However, DEL selection on RNA is challenged by unintended DNA-RNA interactions, which can disrupt small molecule binding and bias selection results. To mitigate biases associated with standard solution-phase DEL selection, methods such as co-incubation with RNA fragment competitors, elimination of highly complementary DNA barcodes, and conversion of single-stranded to double-stranded DNA regions have been developed.^26, 28^ Alternatively, employing a One-Bead-One-Compound (OBOC) solid-phase DEL format may inherently minimize such bias, as the DNA barcode is present at a much lower stoichiometry relative to the compound (1:2500).^29, 30^ This approach has proven effective in both target-agnostic library-versus-library screens and target-centric selections.^24, 25^

Because commercial DELs are typically solution-phase and designed to maximize chemical diversity at the billion-to trillion-compound scale, scaffolds that preferentially bind RNA are often underrepresented (e.g., compounds containing multiple hydrogen-bond donors, aromatic rings, and heteroaromatic nitrogens). Developing an RNA-focused solid-phase DEL through the careful selection of building blocks with RNA-engaging features could accelerate ligand discovery and expand the DEL toolkit. Our previous success in the construction of a small benzimidazole-focused library (∼13,000 diversity)^25^ in house inspired the design of a significantly larger library (∼580,000), termed Ribo-DEL. Ribo-DEL incorporated a broader spectrum of known RNA-binding scaffolds and increased overall diversity, providing the potential to identify previously unrecognized RNA-binding chemotypes. Moreover, a suite of computational tools was employed to streamline hit analysis and validation, integrating structure-activity relationship analysis, physicochemical property profiling, and molecular docking. These components establish a streamlined and generalizable platform for RNA-targeted ligand discovery.

To demonstrate the utility of Ribo-DEL in RNA-targeted ligand discovery, the r(G_4_C_2_)^exp^ system was chosen as a model. The r(G_4_C_2_)^exp^ is abnormally expanded in the *C9orf72* locus, and is the most common genetic cause of amyotrophic lateral sclerosis and frontotemporal dementia (ALS/FTD), two neurodegenerative diseases characterized by motor neuron loss and behavioral deficiencies.^31–33^ The r(G_4_C_2_)^exp^ transcripts contribute to ALS/FTD pathology by sequestering RNA binding proteins that disrupt their normal function,^34^ or by undergoing repeat associated non-AUG (RAN) translation that leads to the production of toxic dipeptides.^35–38^ Designing small molecule drug that binds r(G_4_C_2_)^exp^ and disrupt its disease-causing functionality could provide a novel therapeutic avenue for treating ALS/FTD. Previous work has identified molecules that modulate r(G_4_C_2_)^exp^-associated disease progression through mechanisms including inhibition of RAN translation, RNase L-mediated transcript degradation, and enhanced intron splicing.^39–41^ Nonetheless, these efforts relied on a rigid tetra-cyclic aromatic core, which presents synthetic challenges and exhibits suboptimal solubility and cellular toxicity. This highlights an opportunity to leverage DEL technology to discover new chemical matter that target the r(G_4_C_2_)^exp^ and mitigate disease hallmarks.

Herein, an RNA-focused DNA-encoded library platform integrating experimental selection with computational analysis was developed. When applied to the r(G_4_C_2_)^exp^, this platform enabled the identification of a lead compound that engages the RNA both *in vitro* and in cells. The compound was subsequently optimized through structure-based design to enhance binding affinity and bioactivity.

## Results

### Design of an RNA-focused DEL

The general design of the Ribo-DEL was based on the previously developed OBOC-derived solid-phase DEL,^29^ in which each bead is loaded with copies of the same compound, and less than 0.4% of DNA code. This design, in contrast to solution-phase DEL where each compound is directly conjugated with its own DNA tag (compound to DNA ratio is 1:1), is advantageous in screening RNA ligands, because it minimizes the potential non-specific interaction between DNA and structured RNA, which could result in false positives in the selection process.

To increase the amount of privileged RNA interacting scaffolds in the new design, commercially available building blocks were surveyed and intersected with chemical matter that have been reported to interact with RNAs. This process was guided by prior works that performed comprehensive analysis on known RNA binders and identified scaffolds or chemical properties that aid in the binding of small molecules to RNA.^24, 42–46^ Indeed, motifs such as quinoline, naphthalene, pyridinium, benzimidazole, as well as other aromatic and heteroaromatic rings were selected (Fig. S1 and Table S1). Physicochemical properties, such as high nitrogen content, increased numbers of hydrogen bond donors and acceptors, lower LogP and LogD values, were also considered as criteria for choosing building blocks.^47, 48^ To avoid biasing the library toward known RNA binders and to enhance overall chemical diversity, various non-aromatic building blocks bearing isobutyl, tetrahydropyran, cyclopropane, or thiopyran moieties were included in the building-block pool. Moreover, DNA-compatible peptoid synthesis and reductive amination reactions were incorporated to diversify the backbone architecture, introducing both “linear” and “branched” compounds (Fig. 1A and Fig. S2C).^49^ This RNA-focused DEL library, namely Ribo-DEL, comprises approximately 580,000 diversified compounds.

**Figure 1.**
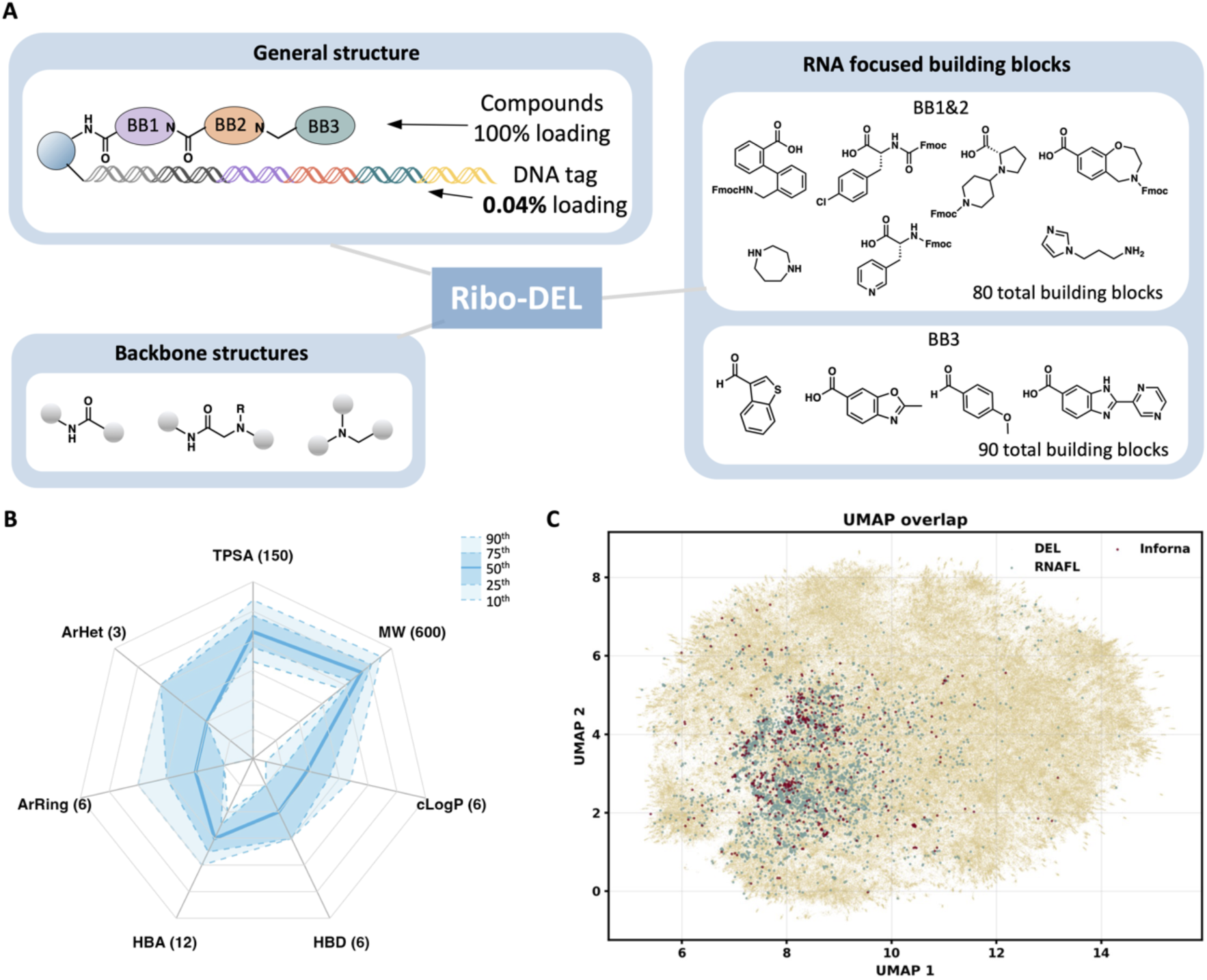
The design and characterization of Ribo-DEL. (A) The design of Ribo-DEL. Each bead is conjugated to small molecules and DNA codes (compound to DNA ratio is 1 : 0.04%). Building blocks are connected via DNA compatible reactions, including peptide coupling, peptoid formation and reductive amination. Representative building block structures in the DEL library are listed. (B) Physicochemical properties of Ribo-DEL. Calculated properties include molecular weight (MW; 0-600), topological polar surface area (TPSA; 0-150), number of aromatic heterocycles (ArHet; 0-3), number of aromatic rings (ArRings; 0-6), hydrogen bond acceptor (HBA; 0-12), hydrogen bond donor (HBD; 0-6), and calculated partition coefficient (cLogP; (−1)−6). Dashed and solid lines, as indicated in the figure, represent the 90^th^, 75^th^, 50^th^, 25^th^, and 10^th^ percentiles. (C) Chemical space analysis of Ribo-DEL. DEL virtual library (light yellow) was analyzed using UMAP analysis and compared with Inforna (red) and RNA-focused library (RNAFL; blue).

The designed DEL library was virtually enumerated and constructed using RDKit, enabling systematic characterization of its chemical properties.^50^ The virtual library was constructed by sequentially enumerating building blocks (BB1-BB3) based on their compatible functional group reactivity, yielding the final Ribo-DEL design. Analysis of the enumerated compounds revealed that the majority exhibited drug-like physicochemical properties, with molecular weights ranging from 350 to 550 g/mol, 0-3 hydrogen bond donors, 4-8 hydrogen bond acceptors, calculated logP values below 3, total polar surface areas (TPSA) under 140 Å², and QED score ranging from 0.3 to 0.8 (Figure 1B and Supplemental files). Moreover, the Ribo-DEL compounds predominantly contained 1-4 aromatic rings and 0-2 aromatic heterocycles, reflecting balanced hydrophobicity and structural complexity well suited for RNA-targeted small-molecule design (Figure 1B). To evaluate the diversity and distribution of the designed library, a chemical space analysis was performed using Uniform Manifold Approximation and Projection (UMAP) with Morgan fingerprints (radius = 2). This approach enables visualization of scaffold similarity, functional group composition, and neighborhood relationships across large molecular collections. Three libraries were analyzed and visualized: (1) Ribo-DEL, (2) Inforna,^42^ a curated database of all published RNA-binding small molecules, and (3) a previously reported RNA-focused library.^44^ The analysis revealed that Ribo-DEL overlaps with known RNA-binding chemical space, confirming its relevance to RNA recognition. Importantly, Ribo-DEL also extends into new regions of chemical space, indicating greater structural and physicochemical diversity relative to existing RNA-targeted collections (Fig. 1C).

### Construction of Ribo-DEL

Ribo-DEL was synthesized using a combinatorial “split-and-pool” method.^29, 51, 52^ TentaGel beads (10 µm, amine-functionalized) were modified to introduce both an alkyne and an Fmoc-protected amine (Fig. S2A). The alkyne was coupled to azide-functionalized headpiece DNA (HDNA) under substoichiometric conditions to control DNA loading, while the Fmoc-protected amine served as the initiation site for chemical synthesis. During “split-and-pool” synthesis, various small-molecule fragments were attached to TentaGel beads in separate reaction wells through DNA-compatible reactions, followed by enzymatic ligation of DNA barcodes that contain the identity of each building block to HDNA (Fig. S2B and C, Table S2). This process was repeated three times yielding a final library with three variable positions that contributed to around 580,000 diversified compounds.

To monitor the quality of the synthesis, 10 μm quality control (QC) beads that are acid-cleavable were synthesized in parallel of the library synthesis. Compounds conjugated to QC beads were cleaved after the synthesis, and their identity were confirmed by matrix-assisted laser desorption/Ionization (MALDI) with time-of-glight (TOF) mass spectrometry (Fig. S3A). Among the three designed backbone structures, QC beads from two backbones exhibited compound masses consistent with expected values, while those from the third did not. As a result, the third backbone was excluded prior to computational analysis and was not incorporated into the final Ribo-DEL library. The integrity of the DNA code was examined by PCR amplification of the library followed by sanger sequencing and next-generation sequencing (NGS) (Fig. S3B and C). Regions of the DNA code (coding sequence, unique molecular index, NGS adaptor) were successfully amplified by both methods, and the frequency of each DNA code was approximately 0.5%-2% (Fig. S3B). Collectively, these results confirmed the good quality and integrity of the constructed library.

### Fluorescence-activated flow cytometry (FACS) for r(G_4_C_2_)^exp^ ligand identification

A pre-established two-color fluorescence activated cell sorting (FACS) strategy was exploited for selecting ligands that bind r(G_4_C_2_)^exp^.^25^ The r(G_4_C_2_)^exp^ folds into two distinct structures in cells, hairpin and G-quadruplex, and this equilibrium can be modulated by the absence or presence of potassium ions in the folding buffer.^41, 53, 54^ Efforts were directed toward screening hairpin binders, as this structure has been shown to drive RNA translation and toxic dipeptide generation, processes directly linked to disease pathology (Fig. 2A), although both structures are therapeutically relevant.^55–57^ During selection, fluorescently-labeled RNA (Dy647-(G_4_C_2_)_8_) and base-paired control RNA (Alexa750-(GC)_8_) were incubated with DEL library (Fig. 2B), and sorted by FACS. Beads that showed high fluorescence in Dy647 and low fluorescence in Alexa750 channel, indicating that they selectively bind r(G_4_C_2_)^exp^ (Dy647-(G_4_C_2_)_8_) over the base-paired control RNA (Alexa750-(GC)_8_), were gated and collected (Fig. 2C). A final hit rate of 0.14% was obtained, yielding approximately 4000 beads for each of the three technical replicates. The bound beads were isolated, PCR amplified and decoded from NGS result to obtain the chemical structure and redundancy of hit compounds. Among three technical replicates, 125 hits overlapped across all three groups with redundancy score larger than 2 (Fig. S4A and B), indicating that these hits were repetitively selected during sorting and represent promising candidates for downstream computational analysis and experimental validation.^58, 59^

**Figure 2.**
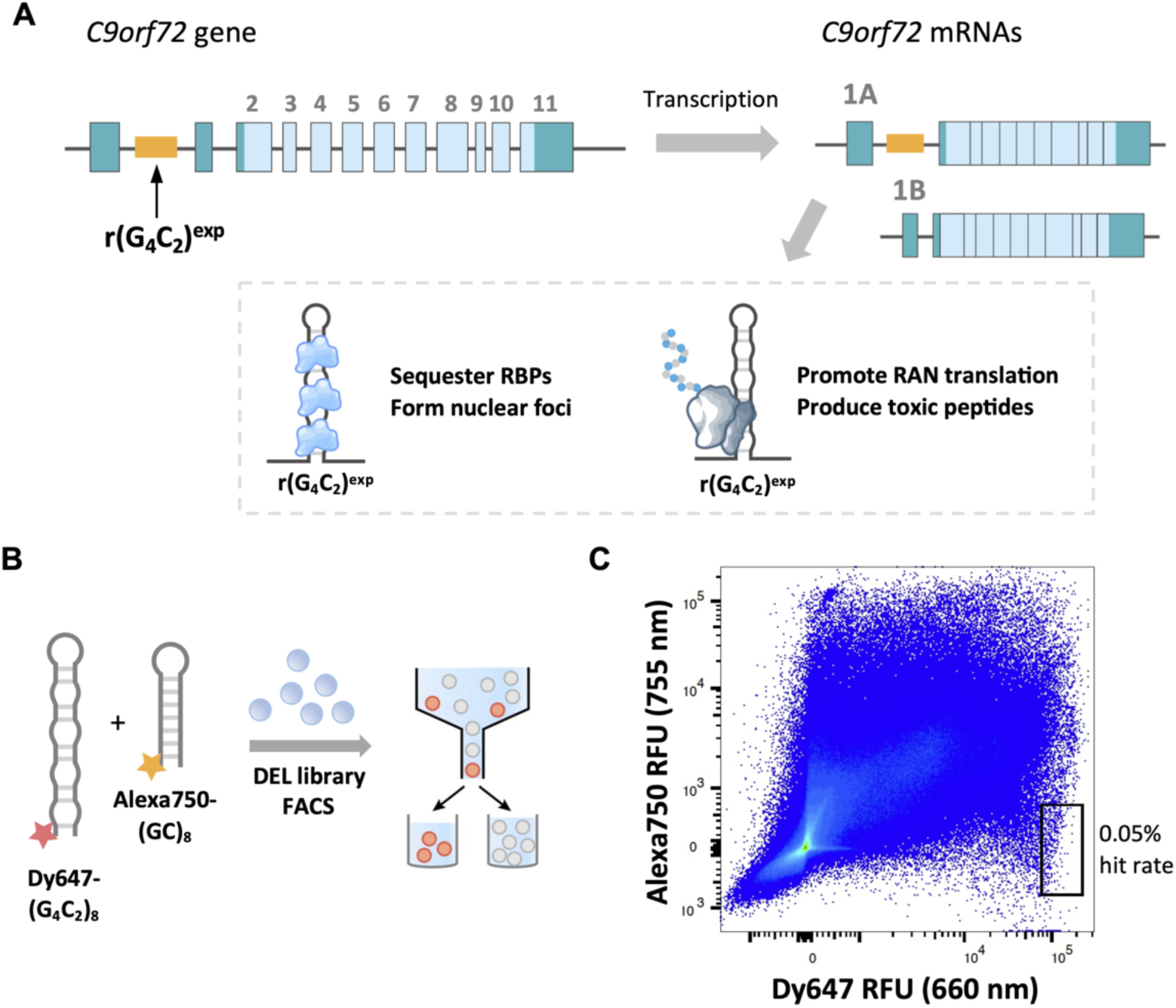
DEL screening for discovery of r(G_4_C_2_)^exp^ ligands. (A) Scheme of the pathology of r(G_4_C_2_)^exp^. Toxic r(G_4_C_2_)^exp^ resides in the intron 1 region of *c9orf72* gene. Once transcribed, the repeat-containing RNA isoform can sequester nuclear proteins or undergo repeat-associated non-AUG (RAN) translation in the cytoplasm. (B) Scheme of the DEL selection process. Dy647 labeled (G_4_C_2_)_8_ and Alexa750 labeled (GC)_8_ RNA were incubated with DEL library and subjected to FACS. (C) Two-color flow cytometry dot plot showing the gating and sorting results. Beads exhibiting high fluorescence in the Dy647 channel and low fluorescence in the Alexa750 channel were collected for further analysis.

### Computational evaluation of hit compounds

DEL is a powerful technology that provides high information content from a single experiment. When integrated with computational analysis, DEL screening results can reveal global patterns of small-molecule binding to target RNA structures and pinpoint the specific chemical moieties that drive these interactions. (Fig. 3A). To identify chemical features enriched during the selection of r(G_4_C_2_)^exp^ hairpin binders, SAR analysis and physicochemical property evaluation were performed on the hit compounds. The enrichment scores of individual building blocks within the hit pool were first assessed to guide SAR interpretation and identify preferentially selected chemical motifs. To visualize combinatorial trends, the hit compounds were plotted in a three-dimensional space, in which each axis represents the building block identity at positions 1, 2, and 3, respectively (Fig. 3B). Notably, hit compounds clustered together and formed two “planes” in the 3D space, indicating that these compounds are variable at position 1 and 2 but share common structures at position 3. Within these planes, additional “lines” were observed for position 2 and 3, demonstrating that chemical scaffold at positions further away from the solid support are more influential in determining RNA-compound interactions. Specifically, imidazole (BB3-91) and quinoline (BB3-71), in combination with homopiperazine (BB2-85) were identified as scaffolds that preferentially interact with r(G_4_C_2_)^exp^ hairpin.

**Figure 3.**
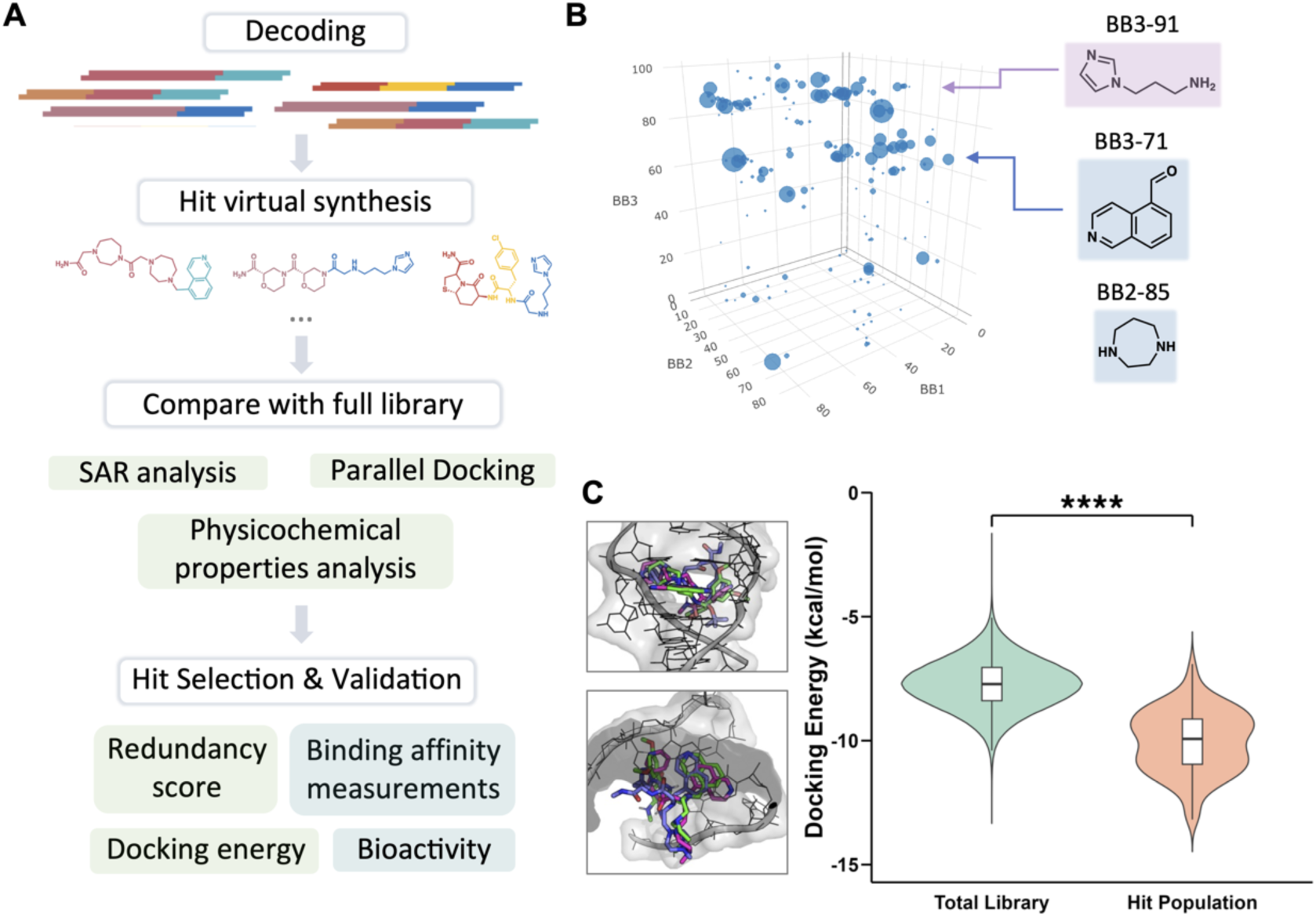
Computational analysis of DEL hits. (A) Scheme of the analysis workflow. NGS results were decoded, and overlapping hits were enumerated. A series of computational analyses was then performed to identify enriched features and guide hit prioritization, including SAR analysis, molecular docking, and physicochemical property evaluation. Hits were prioritized based on redundancy score and docking energy, and subsequently validated through binding affinity measurements and bioactivity assessment. (B) 3D plot illustrating features of building blocks enriched by DEL selection. The three axes represent building block identities at positions 1, 2, and 3. Dot size corresponds to the redundancy score for each hit. Two chemical motifs (labeled in purple and blue) were enriched following DEL selection. (C) Massive parallel docking of the full DEL library. (Left) Docking of the top 10 compounds with the lowest binding energy to a 1 × 1 nucleotide G/G internal loop NMR structure. (Right) Binding energy comparison of Ribo-DEL (Total library) and hit population.

To further extract enriched chemical features from the selection process, hit compounds were virtually synthesized and compared to a representative 5,000-member Ribo-DEL subset derived by k-means clustering.^60^ This representative subset mitigates bias arising from the substantial disparity in sample sizes (125 hits vs. 0.5 million total members). A total of 1,800 Mordred descriptors covering 2D topology, surface charge, and 3D shape were calculated,^61^ and only statistically significant descriptors meeting pre-defined effect-size thresholds were retained to ensure meaningful molecular differentiation. Among all descriptors, several properties known to correlate with RNA-ligand interactions were enriched, including number of basic functional groups, the bond polarizability index (associated with π-π interactions), and aromatic and heteroaromatic ring counts (Fig. S5 and Table S4). Specifically, 10-membered fused rings systems were enriched, likely reflecting the prevalence of quinoline motif in the hit population. A subset of higher-order global descriptors associated with van der Waals surface area (VSA) and spectral topological indices (SpMAD_Dz and SpDiam_Dz) were enriched in the hit population, indicating molecular surfaces with high polarizability, elongated topology, heavy-atom contributions, and electronic heterogeneity (Fig. S5 and Table S4). These observations support a binding mode in which ligands engage r(G_4_C_2_)^exp^ through electrostatic complementarity and π– π stacking interactions. In contrast, several local topological descriptors (ATSC1i, ATSC2c, MATS2c, MATS2are, etc.) were de-enriched in the hit population, indicating that uniform distribution of charge, electronegativity, and branching is not favored for binding (Fig. S5 and Table S4). Together, analyses of both higher-order global descriptors and atom-level features support a binding mode wherein r(G_4_C_2_)^exp^ ligands engage through electrostatic interactions mediated by basic or charged groups and π–π stacking facilitated by aromatic scaffolds.

To investigate the molecular basis of binding for each compound, massive parallel docking leveraging the speed and accuracy of AutoDock-GPU was performed on both the hit population and the full library using a model RNA containing a 1 × 1 nucleotide G/G internal loop.^39, 62^ Excitingly, the hit population exhibited significantly lower average binding energy to the RNA target compared with the total library (−10.1 vs −7.7 kcal/mol) (Fig. 3C), whereas molecular weight distributions were comparable (Fig. S5N), ruling out potential bias from higher molecular weight compounds during docking. This energetic difference suggests that the selection effectively filtered out weak or nonspecific interactions, preferentially retaining compounds with favorable molecular complementarity to the RNA binding pocket. An in-depth examination of the binding poses of the top hits revealed that building blocks at position 3 insert into the 1 × 1 nucleotide G/G internal loop, forming π-π interactions with adjacent guanine and cytosine bases, whereas building blocks at positions 1 and 2 extend outward from the pocket toward the major groove, some forming hydrogen bonds with the Hoogsteen edge of the base pairs (Fig. 3C).

In conclusion, the combined SAR analysis, Mordred descriptor evaluation, and docking study consistently point to the same group of chemical features and interaction patterns underlying RNA-ligand recognition. This agreement highlights the predictive value of integrated computational analysis for guiding the rational development of RNA-targeted therapeutics.

### Experimental validation of hit compounds

For experimental validation, hit compounds were ranked based on their redundancy score obtained from the selection as well as their docking energies to the (G_4_C_2_)_8_ NMR construct, assigning twice as much weight to experimental evidence as to computational predictions. Manual examination of the 10 highest-ranked hits confirmed that they incorporated the prevalent building blocks at positions 2 and 3 and exhibited physicochemical features enriched in the hit population compared with the full library. These hits were then synthesized in-house, and seven were confirmed to be cleanly synthesized based on liquid chromatography-mass spectrometry (LC/MS) analysis. These seven compounds were then evaluated experimentally (Fig. S6A).

To assess RNA binding *in vitro*, two orthogonal assays were performed in parallel to ensure the robustness of the validation results. Biolayer interferometry (BLI) was used to test each compound at a single dose (100 μM) against biotin-(G_4_C_2_)_8_ and biotin-(G_2_C_2_)_8_ (Fig. S6B). Three of the seven compounds displayed greater binding response to biotin-(G_4_C_2_)_8_ than to biotin-(G_2_C_2_)_8_, indicating preferential binding to the structured region of (G_4_C_2_)_8_ rather than nonspecific association with the RNA backbone.

In addition, Chemical Cross-Linking and Isolation by Pull-down (Chem-CLIP), a covalent-based target validation strategy, was employed to evaluate compound binding to target RNA.^63, 64^ In a Chem-CLIP experiment, a small molecule probe functionalized with a diazirine and an alkyne handle is incubated with radio-labeled RNA. Upon UV irradiation, the diazirine group is activated and covalently crosslinked to proximal RNA. Cross-linked RNA is then pulled down via a copper-catalyzed “click” reaction and quantified by measuring radio activity. Competitive-Chem-CLIP (C-Chem-CLIP) was initially performed using a published Chem-CLIP probe **10** that was known to bind (G_4_C_2_)_8_ (Fig. S6C).^40^ Four of the seven compounds competed with **10** by at least 50% at the highest concentration tested (100 μM), suggesting that they occupied the same binding site as **10** (Fig. S6C).

The result from the BLI and C-Chem-CLIP assays together identified compound **1a** as the lead binder of r(G_4_C_2_)^exp^, as it showed the highest selectivity in BLI and strongest competition with **10** in the C-Chem-CLIP assay (Fig. S6). Follow-up BLI analysis of the N-propylamide version of **1a** (**1**) revealed that it binds to (G_4_C_2_)_8_ with a Kd of 38 μM, whereas its affinity for (G_2_C_2_)_8_ was weak (> 100 μM; Fig. 4A and B). To further assess the target engagement *in vitro* and in cell, compound **2**, the Chem-CLIP probe version of **1**, as well as a control probe **3** that does not contain RNA binding module were synthesized (Fig. 4A). Compound **2**, but not **3**, dose-dependently pulldown ^32^P-(G_4_C_2_)_8_ *in vitro*, capturing 16% of total RNA at 50 μM (Fig. 4C). The modest pulldown efficiency of **2** may be attributed to the potential quenching of diazirine by water. Notably, neither **2** nor **3** pulled down the base-paired RNA (GC)_8_ at concentrations up to 50 μM, supporting that **2** binds to the G/G internal loop of r(G_4_C_2_)^exp^ rather than the RNA backbone.

**Figure 4.**
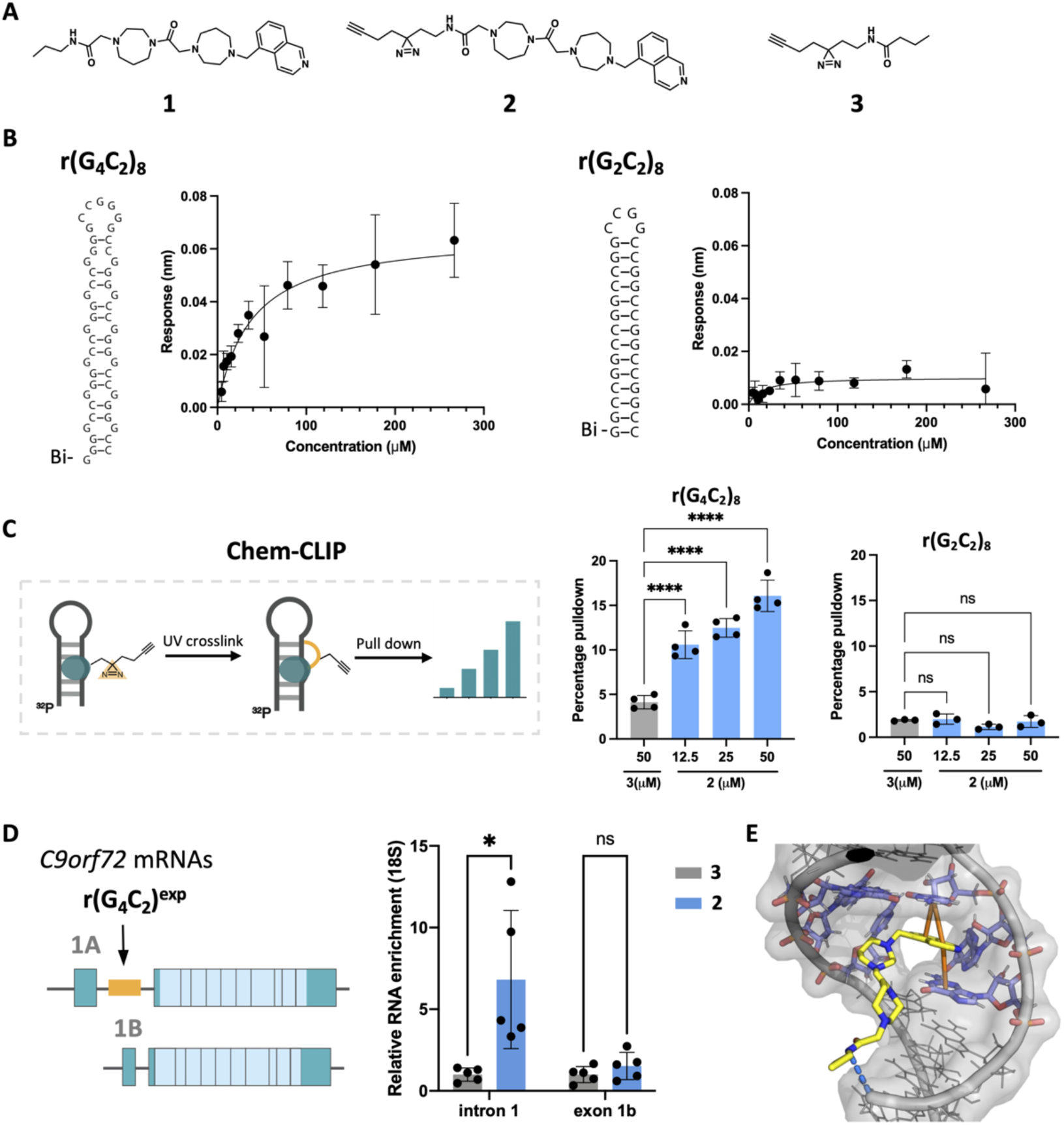
Compound **1** engaged r(G_4_C_2_)^exp^ *in vitro* and in cells. (A) Structures of compound **1** (lead compound) 2 (Chem-CLIP-probe) and 3 (control probe). (B) Biolayer Interferometry experiment of **1** binding to Bi-(G_4_C_2_)_8_ construct (left) and Bi-(G_2_C_2_)_8_ (right). (C) Chem-CLIP pulldown of ^32^P-(G_4_C_2_)_8_ and ^32^P-(G_2_C_2_)_8_ by **2**. (D) In cell pulldown of compound **2** (Chem-CLIP- probe) in C9/ALS patient-derived iPSCs. Compound **2** or **3** were treated to cell culture overnight and cross-linked via UV irradiation. Cross-linked RNA was extracted and pulldown by azide beads and quantified by RT-qPCR. (E) Docking study of **1** against a model of 1 × 1 nucleotide G/G internal loop. Hydrogen bonding interactions were depicted as blue dashed lines and π-π stacking interactions were depicted as yellow solid lines. *, p < 0.05; **, p < 0.01;as determined by a one-way ANOVA with multiple comparisons.

The target engagement was next examined in c9/ALS patient-derived iPSCs. In this disease model, *C9orf72* transcript undergo alternative splicing to yield one isoform containing exon 1a and intron 1, which harbors the repeat expansion, and another containing exon 1b, which lacks the toxic repeats (Fig. 4D). Two sets of primers that amplify specifically intron 1 and exon 1b were designed to evaluate the presence of each transcript. Indeed, **2** enriched the repeat-expansion-containing transcript by 6-fold relative to control probe **3**, but did not pull down the transcript lacking the repeat expansion, as measured by quantitative reverse transcription polymerase chain reaction (RT-qPCR) (Fig. 4D). The RNA engagement in cells was further confirmed by competition with the binder compound **1** (Fig. S6E). Together, these results demonstrate that compound **1** binds the target RNA both *in vitro* and in cells, validating its on-target engagement and selectivity.

### Structure expansion of lead compound 1 to enhance binding affinity

To improve the potency of compound **1** while maintaining its high selectivity, structure-based optimization was carried out. Because **1** was identified from solid-phase DEL, it could be readily modified at its original attachment site to the solid support. Docking analysis of **1** revealed that the quinoline moiety inserts into the 1 × 1 nucleotide G/G internal loop, forming π-π interactions with adjacent guanine and cytosine residues (Fig. 4E). The homopiperazine moiety extends outward from the loop and formed hydrogen bonds with the phosphate backbone, suggesting that this region is amenable to linker attachment and modification.

Since r(G_4_C_2_)^exp^ is a repeat expansion that presents a linear array of potential small-molecule binding sites, it is well suited for dimeric ligands designed to engage multiple pockets simultaneously.^40, 65–67^ Optimizing linker length to align binding modules across adjacent pockets on r(G_4_C_2_)^exp^ was expected to enhance both affinity and avidity. In the FARFAR2-generated^68^ 3D model of r(G_4_C_2_)₈, the two binding pockets are positioned approximately 12.4 Å apart when measured linearly, although the effective separation is likely greater in the native folded RNA due to conformational flexibility (Fig. S7A). The homopiperazine moiety of monomer **1**, which projects from the internal loop, may already function as a partial linker to bridge this span. Based on these considerations, three dimers with linkers containing a central amine and polyethylene glycol (PEG) segments of varying lengths were synthesized (Fig. 5A; compound **11**–**13**). The inter-nitrogen distances between the homopiperazine moieties of the monomer units were 13.2 Å, 19.6 Å, and 27.6 Å, for **11**-**13**, respectively (Fig. S7B). Based on linker length modeling, **11** was predicted to achieve the most favorable alignment of its binding modules with adjacent RNA. Consistent with this prediction, **11** exhibited a >22-fold improvement in binding affinity relative to the monomer (1.7 μM vs. 38 μM) (Fig. 5A and S8A). Compound **12** also showed enhanced binding (3.3 μM; ∼11-fold improvement) whereas **13** did not display saturated binding at concentrations tested (Fig. 5A and S8B and C). These results suggest that dimerization with shorter linkers enhances the binding avidity of lead compound **1**, with compound **11** demonstrating the highest affinity and likely the most favorable binding orientation. All dimer compounds displayed minimal binding to the base-paired control in BLI, similar to compound **1**, indicating that structural dimerization enhanced binding affinity without compromising selectivity (Fig. S8). Notably, dimer **11** showed a superior binding affinity gain (22-fold), compared to previous examples targeting r(CUG)^exp^ (15- and 4-fold)^66, 67^ and r(G_4_C_2_)^exp^ (13-fold)^40^. These findings support an avidity-driven binding mechanism, whereby multiple modules on a single library bead simultaneously engage the multimeric r(G_4_C_2_)₈ RNA, increasing complex stability and persistence during washing and FACS procedures. This phenomenon was also observed in protein-targeted OBOC screenings.^69^

**Figure 5.**
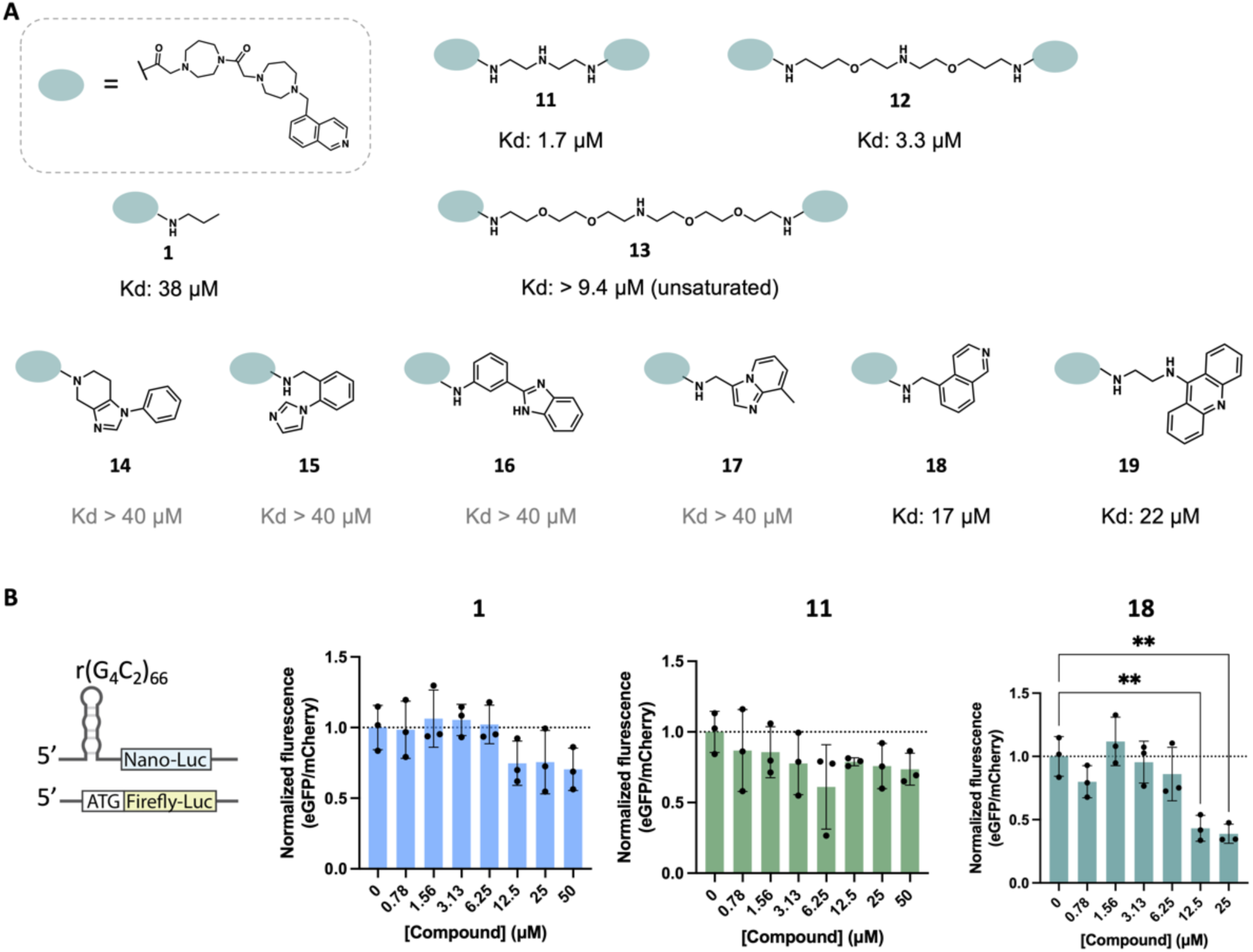
Dimerization and structural expansion of compound 1. (A) Structures and binding affinities of derivatized compounds (**11-19**) as measured by BLI. (B) Luciferase-based RAN translation assay to assess compound bioactivity. Left to right: Compound **1**, **11** and **18**. *, p < 0.05; **, p < 0.01;as determined by a one-way ANOVA with multiple comparisons.

### Optimization of Compounds to Enhance Inhibition of Repeat-Associated Non-AUG (RAN) Translation

The bioactivity of the lead compound **1** and dimers **11**-**13** was evaluated using a luciferase-based RAN translation assay. A plasmid encoding the G_4_C_2_ repeat expansion fused to a NanoLuc luciferase open reading frame lacking a start codon (No ATG - r(G_4_C_2_)_66_ - NanoLuc luciferase) was co-transfected into HEK293T cells along with a control plasmid (SV40 - Firefly luciferase) that undergoes canonical translation. After normalization to firefly luciferase, NanoLuc luciferase signal was dose-dependently reduced by compound **1** (Fig. 5C), inhibiting 25% luciferase signal at 12.5 μM. This indicated that compound **1** was affecting RAN translation but not canonical translation. Despite its markedly enhanced binding affinity *in vitro*, compound **11** did not show stronger RAN translation inhibition than **1** (Fig. 5B), possibly due to reduced cellular permeability.

To improve cellular activity while reducing molecular weight, a small-scale structure expansion of **1** was undertaken. Docking analysis suggested that the homopiperazine moiety of **1** could act as a natural linker, enabling conjugation of aromatic and heteroaromatic scaffolds capable of forming additional π-π interactions near the binding pocket (Fig. 4E). A focused series of amine-containing analogs (compounds **14-19**) was therefore synthesized via peptide coupling to the acid derivative of **1** (**1-COOH**) (Fig. 5A). Among these, compound **18** and **19**, bearing either a second quinoline moiety or an acridine moiety, showed improved binding affinity (Fig. S8D). Compound **18** showed the strongest binding to biotin-r(G_4_C_2_)_8_ by BLI, representing a two-fold affinity improvement (Kd = 17 μM; Fig. S8D). In RAN translation assays, compound **18** enhanced inhibition by ∼2.6-fold relative to **1**, suppressing 66% of NanoLuc signal at 12.5 µM (Fig. 5B). To further assess their biological activity in a disease-relevant context, compounds **1** and **18** were tested in c9ALS patient derived iPSCs, where inhibition of RAN translation was monitored by western blot analysis of toxic dipeptide repeats. Although compound **1** showed a dose-dependent trend toward reducing poly(GA) and poly(GP) levels, the effects were not statistically significant (Fig. S9A). In contrast, the optimized compound **18** dose-dependently reduced poly(GA) abundance, with a 53% ± 4.7% decrease observed at the 20 μM treatment group (p < 0.05) (Fig. S9B), and mildly decreased poly(GP) levels by about 20% at the same concentration, although not statistically significant. Neither compound had an effect on C9ORF72 protein abundance, indicating that they selectively inhibit RAN translation of the dipeptide repeats without interfering with canonical translation of *c9orf72* mRNA. Taken together, these results demonstrate that compound **18**, through rational structure expansion, enhances the activity of **1** by strengthening RNA binding and suppressing RAN translation in both reporter and patient-derived models.

## Conclusion

In this work, a rationally designed, diversified, and RNA-focused DNA-encoded library (DEL) was developed and applied to discover small-molecule ligands targeting the toxic RNA structure r(G_4_C_2_)^exp^ implicated in ALS/FTD. A two-color fluorescence-activated sorting strategy, followed by a series of computational analysis, was utilized to evaluate the binding preferences and interaction modalities of hit compounds with the r(G_4_C_2_)^exp^ hairpin. The top hits were synthesized and evaluated both *in vitro* and in patient-derived cell models. Compound **1** emerged as a promising binder that engaged r(G_4_C_2_)^exp^ hairpin both *in vitro* and *in cellulo*, and alleviated ALS-associated phenotypes by reducing RAN translation in a luciferase reporter assay and in patient-derived iPSCs. Previously, several works have identified bioactive r(G_4_C_2_)^exp^ ligands.^39, 40^ However, these compounds often feature multiple rigid ring structures and high molecular weight. In contrast, compound **1** has low molecular weight and is synthetically accessible due to the robust chemistry of DEL construction. To further enhance the affinity and potency of **1**, dimerization and structure-expansion strategies were implemented. Dimerization improved binding affinity by ∼22-fold without compromising selectivity, whereas structural expansion enhanced bioactivity by ∼2.6-fold in a luciferase-based RAN translation assay.

DNA-encoded library technology and its application to RNA are undergoing rapid development and advancement. The focus of DEL synthesis has shifted from generating billion-scale chemical diversity toward the strategic selection of building blocks and DNA-compatible reaction, enabling high-quality compounds tailored to specific molecular recognition events.^70–73^ To date, limited attention has been devoted to the to the rational design of RNA-targeted DELs. To address this gap, reported analyses of RNA-binding chemical features were consolidated and incorporated into the design of the present RNA-focused DEL. Furthermore, the high information content of DEL selection was leveraged to elucidate RNA-small molecule binding mechanisms specific to r(G_4_C_2_)^exp^, through integrated SAR analysis, structural docking, and physicochemical property evaluation. The integration of DEL selection results with computational docking provided a complementary framework for identifying and prioritizing hit candidates, thereby increasing confidence in their binding potential prior to experimental validation. In future studies, we aim to implement a weighted scoring framework that integrates redundancy scores, docking energies, building-block enrichment, and physicochemical property enrichment (relative to the full library), in combination with machine learning approaches, to further guide hit selection and experimental validation.^74, 75^

Collectively, this work advanced the implementation of DEL technology for RNA therapeutics by (1) designing and constructing a diverse, RNA-focused DEL; (2) integrating computational approaches into ligand identification; (3) discovering a low-molecular weight, selective r(G₄C₂)exp ligand; and (4) optimizing this lead compound through docking-guided dimerization and structural expansion to achieve bioactivity in ALS/FTD disease-model cell lines. Taken together, these advances establish Ribo-DEL as a generalizable discovery platform for identifying and optimizing small molecules that modulate RNA structure and function.

## Acknowledgements

This study was funded by the NIH (R35NS116846 to M.D.D.) and the Muscular Dystrophy Association Development Grant 963835 (to A.T.). We thank Q. M. R. Gibaut for providing the linker used to make compound **12**-**13**.

