## Supplemental data for "RNA-Focused DNA-Encoded Library Construction, Screening, and Integration of Docking Identify Bioactive Ligands of Pathogenic r(G_4_C_2_)^exp^ RNA"

This PDF file includes:

- Supplementary Text Figs. S1 to S8
- Tables S1 to S3
- Detailed Methods for all experimental procedures
- References for SI reference citation

### SUPPLEMENTARY FIGURES AND TABLES

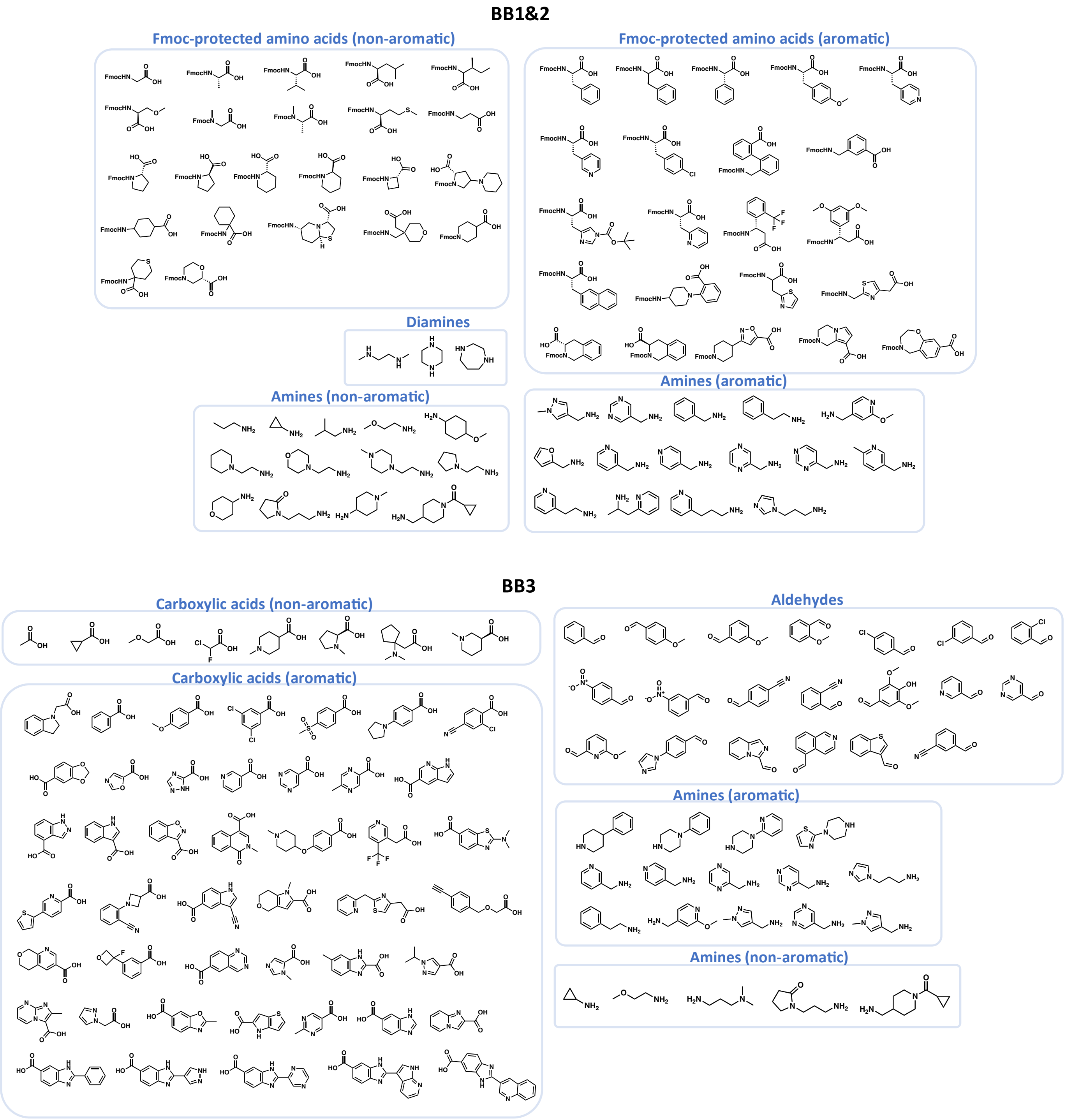


**Figure S1**. Building blocks of DEL-2G. Building blocks 1 and 2 are composed of the same pool of compounds, including Fmoc-protected amino acids, amines and diamines. Building blocks 3 contain carboxylic acids, aldehydes and amines.

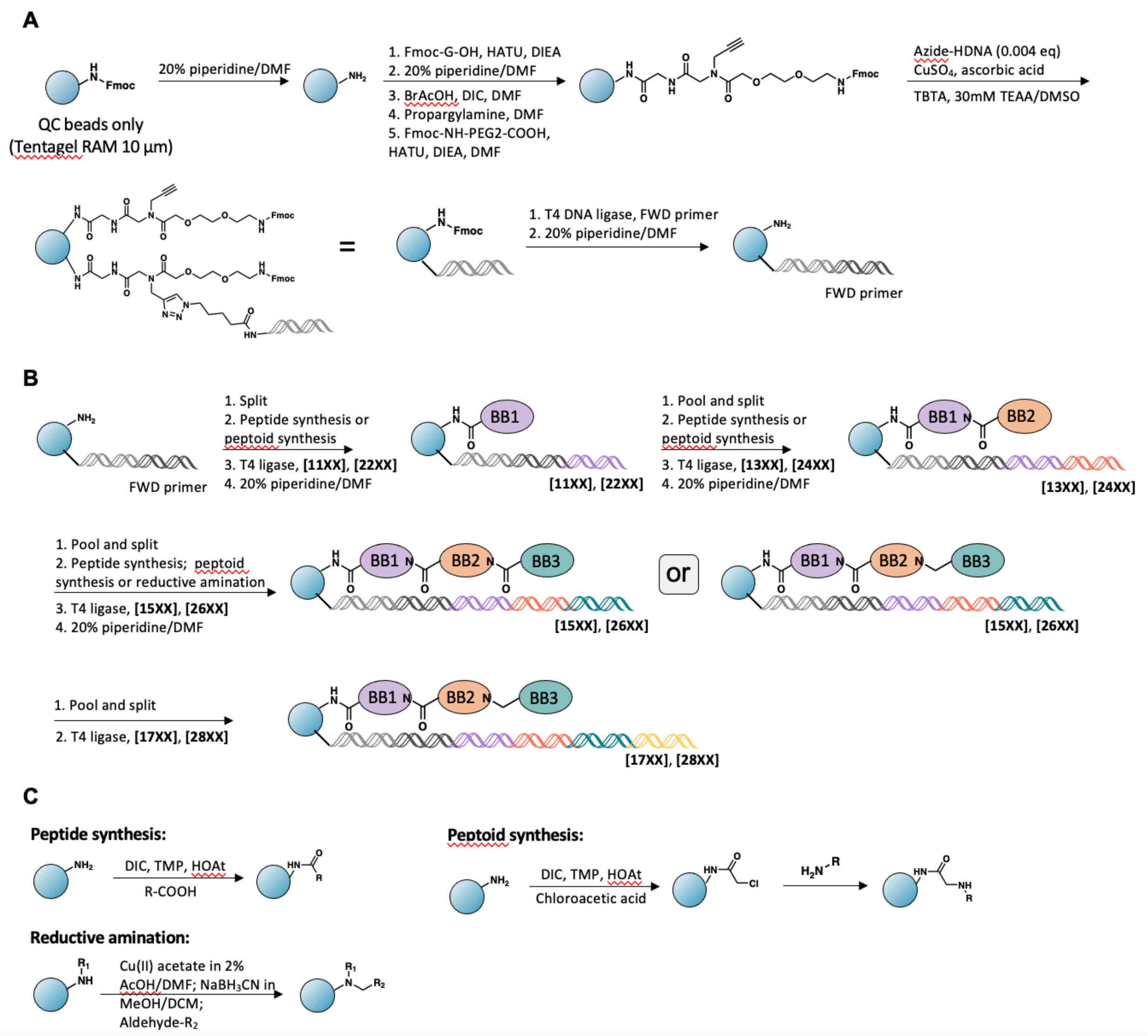


**Figure S2**. Synthesis of DEL-2G. (A) Schematic of head piece synthesis for solid-phase DEL. (B) Schematic of split and pool synthesis. (C) DNA-compatible reactions used in this study.

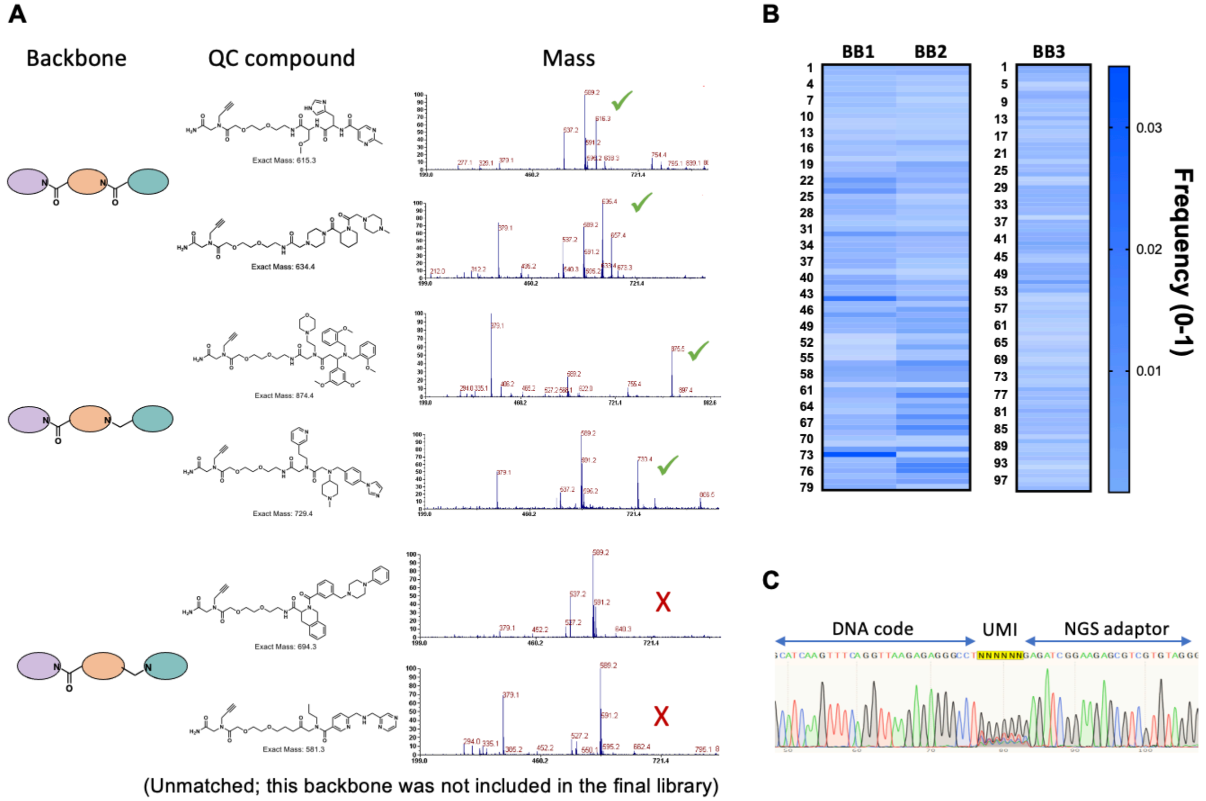


**Figure S3**. Quality control of DEL-2G. (A) Mass detection of quality control beads, as measured by matrix-assisted laser desorption ionization time-of-flight (MALDI-TOF). (B) Next-generation sequencing assessment of native DEL library. The frequency of each building block was plotted as a heat map. (C) Sanger sequencing of quality control beads.

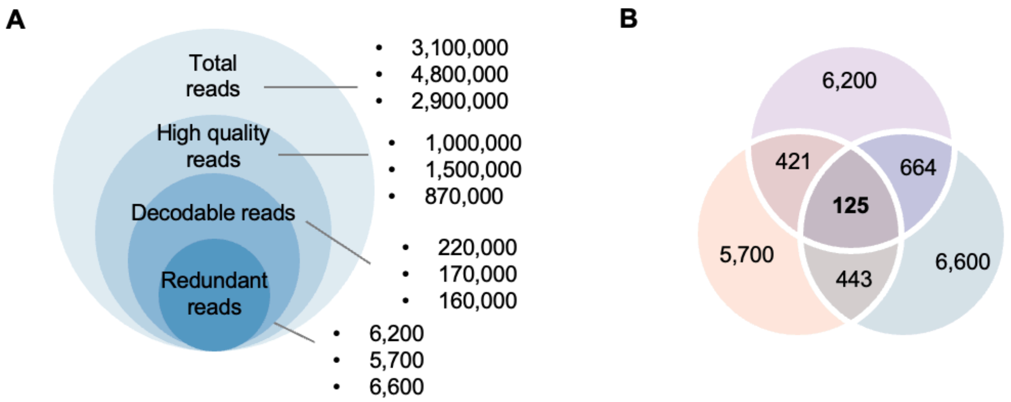


**Figure S4.** Decoding of $G_4C_2$ DEL screening. (A) Filtration of sequencing reads. For each technical replicate, approximately 3 million reads were obtained, of which 1 million were high-quality, 0.2 million were decodable, and ~6,000 were redundant and unique. (B) Overlap of three replicates. A total of 125 redundant reads were present in all three replicates and were used for computational analysis.

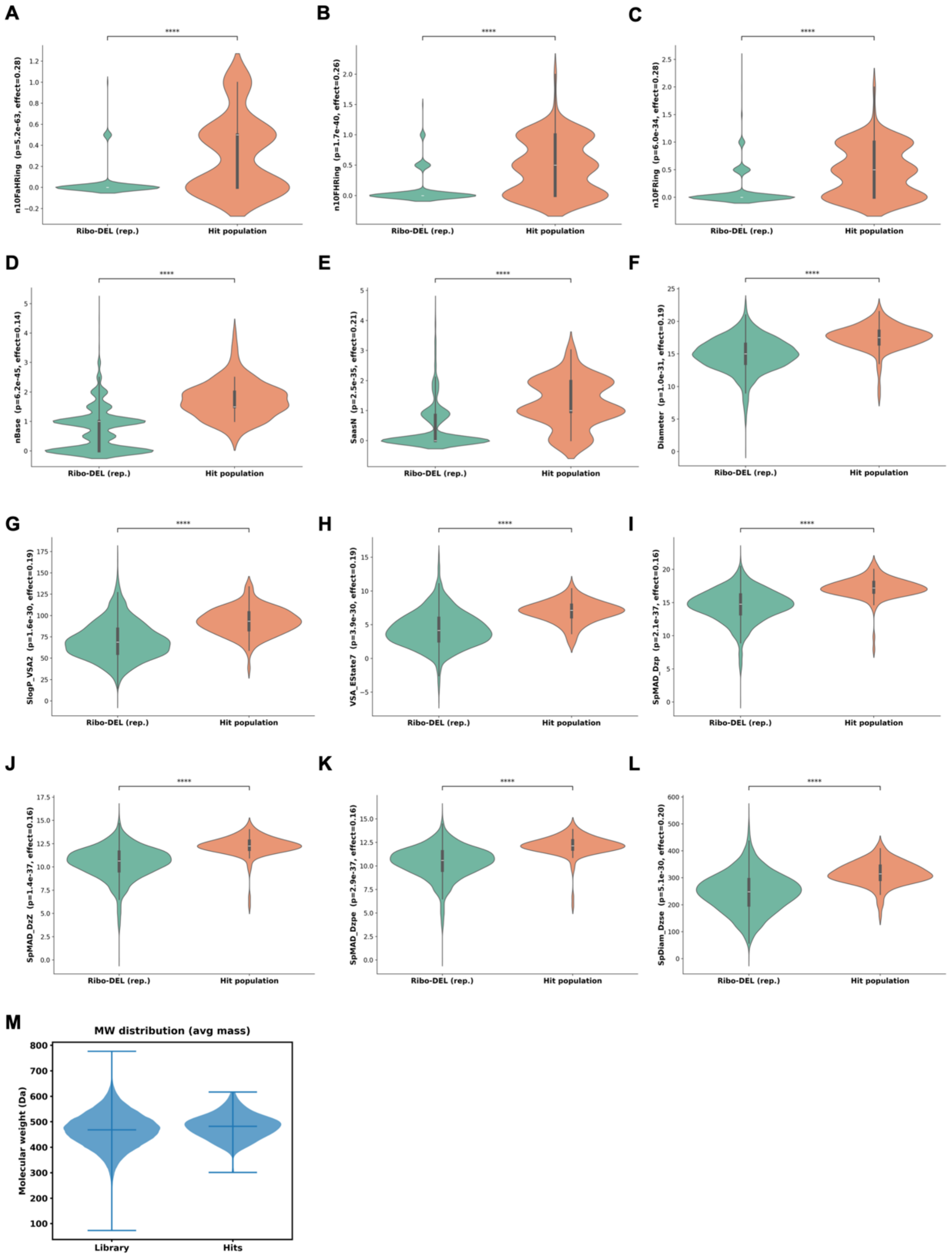


**Figure S5.** Representative physicochemical property comparisons between Ribo-DEL (representative or full library) and hit population. (A) n10FaHRing: 10-membered aliphatic fused hetero ring count. (B) n10FHRing: 10-membered fused hetero ring count. (C) n10FRing: 10-membered fused ring count). (D) nBase: basic group count. (E) SaasN: sum of aasN (nitrogen atom bonded to two aliphatic (single-bonded) carbons (“aa”) and one other substituent). (F) Diameter: geometric diameter. (G) SLogP_VSA2: MOE logP VSA Descriptor 2 (-0.40 <= x < -0.20). (H) VSA_EState7: VSA EState Descriptor 7 ( 6.07 <= x < 6.45). (I) SpMAD_Dzp: spectral mean absolute diviation from Barysz matrix weighted by polarizability. (J) SpMAD_DzZ: spectral mean absolute diviation from Barysz matrix weighted by atomic number. (K) SpMAD_Dzpe: spectral mean absolute diviation from Barysz matrix weighted by pauling EN. (L) SpDiam_Dzse: spectral diamiter from Barysz matrix weighted by sanderson EN. (M) Molecular weight distributions of the full Ribo-DEL and hit populations.

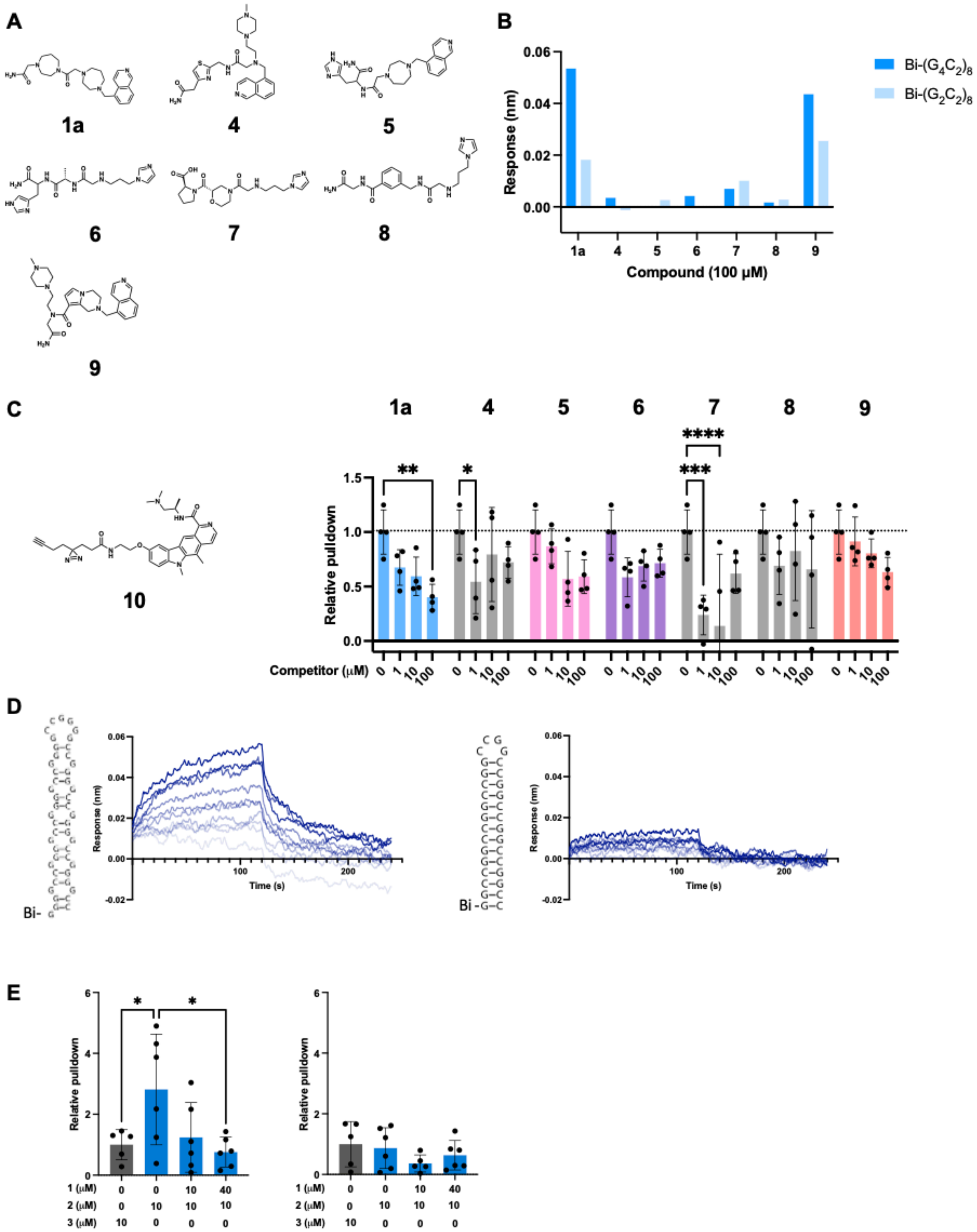

**Figure S6.** *In vitro* binding validation of hit compounds. (A) Structures of hit compounds. (B) Single-dose biolayer interferometry response for compounds binding to Bi-$(G_4C_2)_8$ and Bi-$(G_2C_2)_8$ RNA. (C) *In vitro* Comp-ChemCLIP to assess target engagement. (D) Average response over time for compound **1** binding to Bi-$(G_4C_2)_8$ (left) and Bi-$(G_2C_2)_8$ (right) constructs (from three replicates). Concentration tested ranged from 4.6 to 266 µM (1:1 serial dilution, light to dark blue). (E) Competitive Chem-CLIP in cells. *, $p < 0.05$; **, $p < 0.01$; as determined by a one-way ANOVA with multiple comparisons.

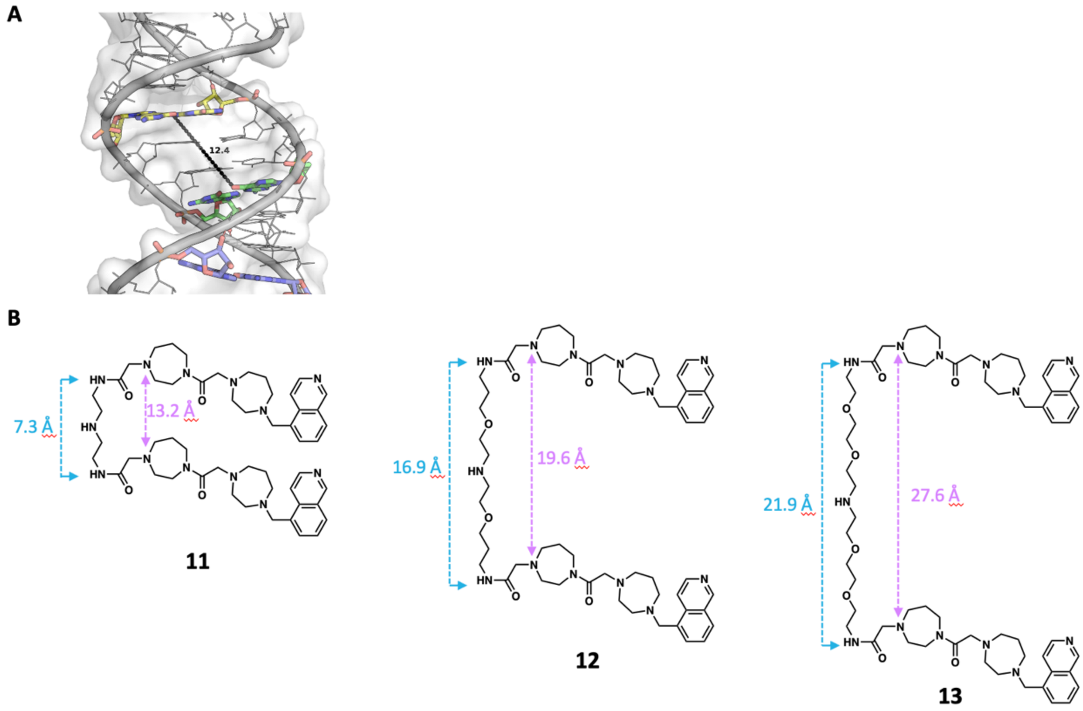


**Figure S7.** Linker design of dimeric compounds. (A) Distance between the two binding pockets measured in a FARFAR2 generated $(G_4C_2)_8$ RNA model. (B) Linker length of measurement in 3D space. Compounds **11-13** were converted to 3D structure and geometry-optimized using Psi4 (B3LYP/6-31G*). For each dimer, the distances between the two nitrogen atoms on the amide (blue arrows) or homopiperiazine (purple arrows) groups were measured to assess linker span.

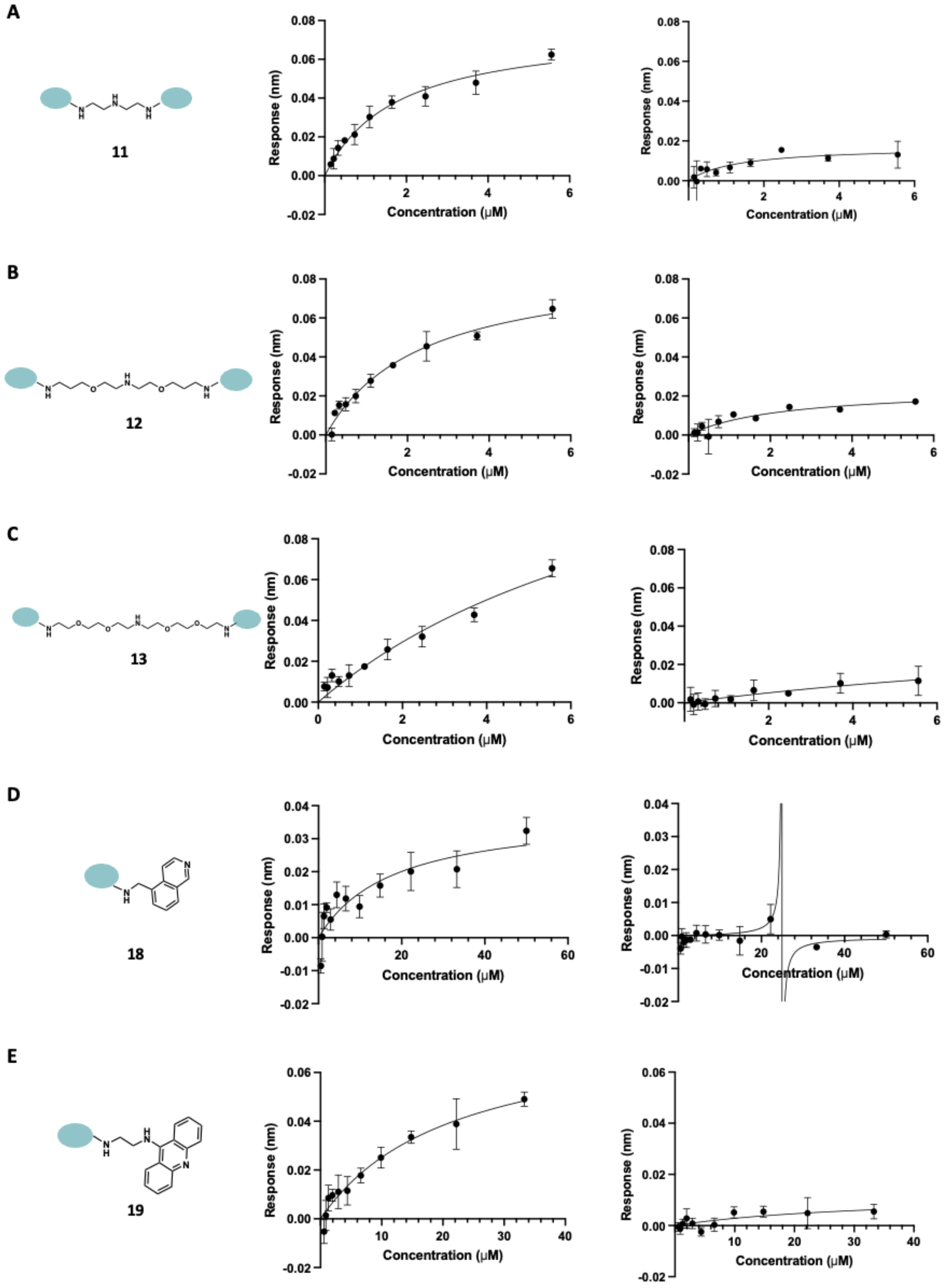


**Figure S8.** Compound affinity measured by biolayer Interferometry**.** (A-E) Response–concentration plots of compounds **11 - 13**, **18** and **19** binding to Bi-$(G_4C_2)_8$ (left) and Bi-$(G_2C_2)_8$ (right) constructs.

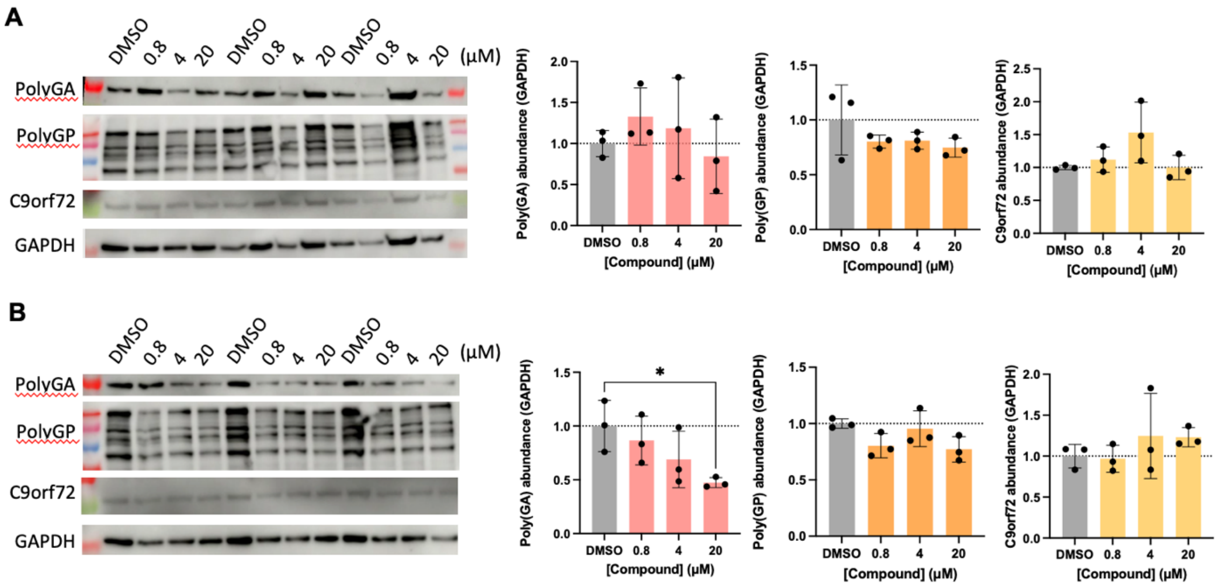


**Figure S9** Bioactivity of **1** and **18** in patient derived iPSCs. The effect of **1** (A) and **18** (B) on poly(GA), poly(GP), and c9orf72 protein abundance, as measured by western blotting and quantified by ImageJ. *P < 0.05, **P < 0.005, ***P < 0.001 , and ****P < 0.0001, as determined by a one-way ANOVA with multiple comparisons. Error bars indicate SD.

**Table S1.** Chemotype analysis of DEL building blocks

| Chemotype name | Count |
|---|---|
| Benzene | 36 |
| Pyridine | 19 |
| Piperidine | 14 |
| Pyrimidine | 8 |
| Pyrrolidine | 8 |
| 1H-Benzimidazole | 7 |
| 1H-Pyrazole | 7 |
| Piperazine | 6 |
| 1H-Imidazole | 4 |
| Cyclohexane | 3 |
| Cyclopropane | 3 |
| Morpholine | 3 |
| Pyrazine | 3 |
| Thiazole | 3 |
| 1,2,3,4-Tetrahydroisoquinoline | 2 |
| Azetidine | 2 |
| 1,4-Diazepane | 2 |
| 1H-Indole | 2 |
| Oxane 2,3,4,5-tetrahydropyran | 2 |
| 1,2,3,4-Tetrahydropyrrolo[1,2-a]pyrazine | 1 |
| 1H-Pyrrolo[2,3-b]pyridine | 1 |
| 2,3-Dihydro-1H-indole | 1 |
| 2,3,4,5-tetrahydro-1,4-benzoxazepine | 1 |
| Thieno[3,2-b]pyrrole | 1 |
| 5,8-Dihydro-6H-pyrano[3,4-b]pyridine | 1 |
| 1,3-Benzodioxole | 1 |
| 1,3-benzothiazole | 1 |
| 1-Benzothiophene | 1 |
| 1,3-benzoxazole | 1 |
| 1,2-Benzoxazole | 1 |
| Cyclopentane | 1 |
| 1,2-dihydroisoquinoline | 1 |
| 1,2-dihydropyridine | 1 |
| Furan | 1 |
| Imidazo[1,2-a]pyrimidine | 1 |
| Imidazo[1,5-a]pyridine | 1 |

| | |
|---|---|
| Imidazo[1,2-A]pyridine | 1 |
| 1H-Indazole | 1 |
| Isoquinoline | 1 |
| Naphthalene | 1 |
| ((8aS)-3,5,6,7,8,8a-hexahydro-2H-[1,3]thiazolo[3,2-a]pyridine) | 1 |
| 1,4,6,7-tetrahydropyrano[4,3-b]pyrrole | 1 |
| 1,2-oxazole | 1 |
| 1,3-oxazole | 1 |
| Quinazoline | 1 |
| Thiane | 1 |
| Triazole | 1 |

**Table S2.** DNA codes used in DEL synthesis

| DNA code | + | - |
|---|---|---|
| 1101 | /5Phos/ATGGAAGAGAGG | /5Phos/TGACCTCTCTT |
| 1102 | /5Phos/ATGGACGGAGCA | /5Phos/TGATGCTCCGT |
| 1103 | /5Phos/ATGGACAAAGAG | /5Phos/TGACTCTTTGT |
| 1104 | /5Phos/ATGGAAGGAGGT | /5Phos/TGAACCTCCTT |
| 1105 | /5Phos/ATGGAGAAAGCA | /5Phos/TGATGCTTTCT |
| 1106 | /5Phos/ATGGATAAAGGT | /5Phos/TGAACCTTTAT |
| 1107 | /5Phos/ATGGATAGAAGG | /5Phos/TGACCTTCTAT |
| 1108 | /5Phos/ATGGATGGGAGT | /5Phos/TGAACTCCCAT |
| 1109 | /5Phos/ATGGGCAAAGGA | /5Phos/TGATCCTTTGC |
| 1110 | /5Phos/ATGGTTGAGGAT | /5Phos/TGAATCCTCAA |
| 2201 | /5Phos/TCAAGTTTCAG | /5Phos/AACCTGAAACT |
| 2202 | /5Phos/TCAAACCTCAA | /5Phos/AACTTGAGGTT |
| 2203 | /5Phos/TCAAATCCCAT | /5Phos/AACATGGGATT |
| 2204 | /5Phos/TCAAACCCTAC | /5Phos/AACGTAGGGTT |
| 2205 | /5Phos/TCAATCCTCTC | /5Phos/AACGAGAGGAT |
| 2206 | /5Phos/TCAATTCTCCG | /5Phos/AACCGGAGAAT |
| 2207 | /5Phos/TCACGCCTTCA | /5Phos/AACTGAAGGCG |
| 2208 | /5Phos/TCACGTTCCTG | /5Phos/AACCAGGAACG |
| 2209 | /5Phos/TCACTCTCCAC | /5Phos/AACGTGGAGAG |
| 2210 | /5Phos/TCATCCTCTTA | /5Phos/AACTAAGAGGA |
| 1301 | /5Phos/GTTAAGAGAGG | /5Phos/TAGCCTCTCTT |
| 1302 | /5Phos/GTTACGGAGCA | /5Phos/TAGTGCTCCGT |
| 1303 | /5Phos/GTTACAAAGAG | /5Phos/TAGCTCTTTGT |
| 1304 | /5Phos/GTTAAGGAGGT | /5Phos/TAGACCTCCTT |
| 1305 | /5Phos/GTTAGAAAGCA | /5Phos/TAGTGCTTTCT |
| 1306 | /5Phos/GTTATAAAGGT | /5Phos/TAGACCTTTAT |
| 1307 | /5Phos/GTTATAGAAGG | /5Phos/TAGCCTTCTAT |
| 1308 | /5Phos/GTTATGGGAGT | /5Phos/TAGACTCCCAT |
| 1309 | /5Phos/GTTGCAAAGGA | /5Phos/TAGTCCTTTGC |
| 1310 | /5Phos/GTTTTGAGGAT | /5Phos/TAGATCCTCAA |
| 2401 | /5Phos/CTAAGTTTCAG | /5Phos/GAACTGAAACT |
| 2402 | /5Phos/CTAAACCTCAA | /5Phos/GAATTGAGGTT |
| 2403 | /5Phos/CTAAATCCCAT | /5Phos/GAAATGGGATT |
| 2404 | /5Phos/CTAAACCCTAC | /5Phos/GAAGTAGGGTT |
| 2405 | /5Phos/CTAATCCTCTC | /5Phos/GAAGAGAGGAT |
| 2406 | /5Phos/CTAATTCTCCG | /5Phos/GAACGGAGAAT |
| 2407 | /5Phos/CTACGCCTTCA | /5Phos/GAATGAAGGCG |
| 2408 | /5Phos/CTACGTTCCTG | /5Phos/GAACAGGAACG |
| 2409 | /5Phos/CTACTCTCCAC | /5Phos/GAAGTGGAGAG |

| | | |
|---|---|---|
| 2410 | /5Phos/CTATCCTCTTA | /5Phos/GAATAAGAGGA |
| 1501 | /5Phos/TTCAAGAGAGG | /5Phos/GCGCCTCTCTT |
| 1502 | /5Phos/TTCACGGAGCA | /5Phos/GCGTGCTCCGT |
| 1503 | /5Phos/TTCACAAAGAG | /5Phos/GCGCTCTTTGT |
| 1504 | /5Phos/TTCAAGGAGGT | /5Phos/GCGACCTCCTT |
| 1505 | /5Phos/TTCAGAAAGCA | /5Phos/GCGTGCTTTCT |
| 1506 | /5Phos/TTCATAAAGGT | /5Phos/GCGACCTTTAT |
| 1507 | /5Phos/TTCATAGAAGG | /5Phos/GCGCCTTCTAT |
| 1508 | /5Phos/TTCATGGGAGT | /5Phos/GCGACTCCCAT |
| 1509 | /5Phos/TTCGCAAAGGA | /5Phos/GCGTCCTTTGC |
| 1510 | /5Phos/TTCTTGAGGAT | /5Phos/GCGATCCTCAA |
| 2601 | /5Phos/CGCAGTTTCAG | /5Phos/AACCTGAAACT |
| 2602 | /5Phos/CGCAACCTCAA | /5Phos/AACTTGAGGTT |
| 2603 | /5Phos/CGCAATCCCAT | /5Phos/AACATGGGATT |
| 2604 | /5Phos/CGCAACCCTAC | /5Phos/AACGTAGGGTT |
| 2605 | /5Phos/CGCATCCTCTC | /5Phos/AACGAGAGGAT |
| 2606 | /5Phos/CGCATTCTCCG | /5Phos/AACCGGAGAAT |
| 2607 | /5Phos/CGCCGCCTTCA | /5Phos/AACTGAAGGCG |
| 2608 | /5Phos/CGCCGTTCCTG | /5Phos/AACCAGGAACG |
| 2609 | /5Phos/CGCCTCTCCAC | /5Phos/AACGTGGAGAG |
| 2610 | /5Phos/CGCTCCTCTTA | /5Phos/AACTAAGAGGA |
| 1701 | /5Phos/GTTAAGAGAGG | /5Phos/TAGCCTCTCTT |
| 1702 | /5Phos/GTTACGGAGCA | /5Phos/TAGTGCTCCGT |
| 1703 | /5Phos/GTTACAAAGAG | /5Phos/TAGCTCTTTGT |
| 1704 | /5Phos/GTTAAGGAGGT | /5Phos/TAGACCTCCTT |
| 1705 | /5Phos/GTTAGAAAGCA | /5Phos/TAGTGCTTTCT |
| 1706 | /5Phos/GTTATAAAGGT | /5Phos/TAGACCTTTAT |
| 1707 | /5Phos/GTTATAGAAGG | /5Phos/TAGCCTTCTAT |
| 1708 | /5Phos/GTTATGGGAGT | /5Phos/TAGACTCCCAT |
| 1709 | /5Phos/GTTGCAAAGGA | /5Phos/TAGTCCTTTGC |
| 1710 | /5Phos/GTTTTGAGGAT | /5Phos/TAGATCCTCAA |
| 2801 | /5Phos/CTAAGTTTCAG | /5Phos/AGGCCTGAAACT |
| 2802 | /5Phos/CTAAACCTCAA | /5Phos/AGGCTTGAGGTT |
| 2803 | /5Phos/CTAAATCCCAT | /5Phos/AGGCATGGGATT |
| 2804 | /5Phos/CTAAACCCTAC | /5Phos/AGGCGTAGGGTT |
| 2805 | /5Phos/CTAATCCTCTC | /5Phos/AGGCGAGAGGAT |
| 2806 | /5Phos/CTAATTCTCCG | /5Phos/AGGCCGGAGAAT |
| 2807 | /5Phos/CTACGCCTTCA | /5Phos/AGGCTGAAGGCG |
| 2808 | /5Phos/CTACGTTCCTG | /5Phos/AGGCCAGGAACG |
| 2809 | /5Phos/CTACTCTCCAC | /5Phos/AGGCGTGGAGAG |
| 2810 | /5Phos/CTATCCTCTTA | /5Phos/AGGCTAAGAGGA |

**Table S3.** Primers used in library preparation and RT-qPCR studies

| Primer | Sequence (5' – 3') |
|---|---|
| FWD | CAGACGTGTGCTCTTCCGATCT |
| RVS | ACACGACGCTCTTCCGATCT |
| P7 index 1 | CAAGCAGAAGACGGCATACGAGATCGTGATGTGACTGGAGTT |
| P7 index 2 | CAAGCAGAAGACGGCATACGAGATACATCGGTGACTGGAGTT |
| P7 index 3 | CAAGCAGAAGACGGCATACGAGATGCCTAAGTGACTGGAGTT |
| P5 adaptor | AATGATACGGCGACCACCGAGATCTACACTCTTTCCCTACACGACGCTCTTCCGATCT |
| P7 | CAAGCAGAAGACGGCATACGAGAT |
| P5 | AATGATACGGCGACCACCGAGATCT |
| 18S fw | GTAACCCGTTGAACCCCATT |
| 18S rv | CCATCCAATCGGTAGTAGCG |
| Intron 1a fw | ACGCCTGCACAATTTCAGCCCAA |
| Intron 1a rv | CAAGTCTGTGTCATCTCGGAGCTG |
| Exon1b fw | AGCTGGAGATGGCGGTGGGC |
| Exon 1b rv | CTGCGGTTGCGGTGCCTGC |
| Exon 2-3 fw | ACTGGAATGGGGATCGCAGCA |
| Exon 2-3 rv | ACCCTGATCTTCCATTCTCTCTGTGCC |

### MATERIALS & METHODS

**Materials.** All RNAs (Dye-labeled, biotinylated, and unmodified) were purchased from Dharmacon (GE Healthcare). Biotinylated and unmodified RNAs were deprotected and desalted using PD-10 columns (GE Healthcare) following the vendor's recommended procedure. Dye-labeled RNAs were HPLC purified by the Vendor and used without further processing. RNA concentrations were measured using Beckman Coulter DU 800 UV/vis spectrophotometer. The measured UV absorbances at 260 nm were converted to concentrations using the extinction coefficients provided by Dharmacon. DNA primers were purchased from Integrated DNA Technologies (IDT, Inc.).

**Abbreviations.** AcOH, Acetic acid; BSA, bovine serum albumin; BTPBB, Bis-tris propane breaking buffer; BTPLB, Bis-tris propane ligation buffer; BTPWB, Bis-tris propane washing buffer; Cu(OAc)2, copper(II) acetate; CuSO4, copper(II) sulfate; DCM, dichloromethane, DIC, diisopropylcarbodiimide, DIEA, N,N-diisopropylethylamine; DMF , N,N-dimethylformamide; DMSO, dimethylsulfoxide; DPBS, dulbecco's phosphate-buffered saline; EDTA, ethylenediaminetetraacetic acid; Fmoc, fluorenylmethyloxycarbonyl, Glycine; HATU, 1-[bis(dimethylamino)methylene]-1H-1,2,3-triazolo[4,5-b]pyridinium 3-oxide hexafluorophosphate; HDNA, headpiece DNA; HEPES, 4-(2-hydroxyethyl)-1-piperazineethanesulfonic acid; HOAt, 1-hydroxy-7-azabenzotriazole; HPLC, high-performance liquid chromatography; LiCl, lithium chloride; MALDI-TOF, Matrix-assisted laser desorption ionization time-of-flight; MeOH, methanol; MS, mass spectrometry; NaBH3CN, sodium cyanoborohydride; NGS, next generation sequencing; OP stock, oligonucleotide paired stock; PAGE, polyacrylamide gel electrophoresis; PCR, polymerase chain reaction; QC, quality control; SDS, sodium dodecyl sulfate; TBTA, tris(benzyltriazolylmethyl)amine; TCEP, tris(2-carboxyethyl)phosphine; TEAA, triethylammonium acetate; TFA, trifluoroacetic acid; THPTA, tris(3-hydroxypropyltriazolylmethyl)amine; TIS, triisopropylsilane; TMP, 2,4,6-trimethylpyridine.

### DNA-ENCODED LIBRARY SYNTHESIS AND SELECTION

**Materials.** TentaGel® RAM and $NH_2$ beads were purchased from Rapp Polymere GmbH. The amino-modified HDNA (/5Phos/GAGTCA/iSp9//iUniAmM//iSp9/TGACTCCC) was purchased as an HPLC-purified oligonucleotide from IDT, Inc. All encoding DNA sequences were obtained from IDT, Inc. and listed in **Table S2**. The azide-modified HDNA was prepared from amino-modified HDNA as previously described.[1]

**T4 DNA ligation.** DNA duplexes that encode building blocks information were prepared by mixing the indicated single stranded DNA pairs **[XXXX]** [+] and **[XXXX]** [-] (see **Table S2**) in OP Stock Buffer (1 mM Bis-Tris, pH 7.6, 50 mM NaCl), then annealing at 60℃ for 5 min and slowly cooling to room temperature. Ligations were carried out with 150 µM DNA duplex folded in OP Stock Buffer.

To perform DNA code ligation, Tentagel beads were washed with DMF/water (1:1, ×1), BTPWB (10 mM Bis-Tris, pH 7.6, 50 mM NaCl, 0.04% (v/v) Tween-20) (×3), and BTPLB (100 mM Bis- Tris propane, pH 7.6, 500 mM NaCl, 100 mM MgCl2, 10 mM ATP, 0.2% (v/v) Tween-20) (×3), followed by equilibration in BTPLB for 30 min. After that, ligation mixture containing DNA duplexes and T4 DNA ligase (400 units) in BTPLB were prepared and added to corresponding beads (300 µL total). The ligation mixture was shaken at room temperature overnight and the ligated beads were washed with BTPWB (×3), DMF/water (1:1, ×1), and DMF (×3).

**Standard procedures for DNA-compatible solid-phase synthesis.** *Fmoc deprotection.* Tentagel beads were swelled in DMF for 5 min before the addition of 300 µL 20% (v/v) piperidine in DMF and incubation at room temperature for 5 min with shaking (2 cycles). Beads were then washed with DMF (×3) to remove excess reagents.

*Peptide coupling.* Tentagel beads were treated with carboxylic acid (0.4 M, 70 µL, 4 equiv.) pre-activated with DIC (0.4 M, 70 µL, 4 equiv.), HOAt (0.4 M, 70 µL, 4 equiv.), and TMP (0.4 M,

70 μL, 4 equiv.) in DMF and incubated at room temperature for 2 hrs with shaking, followed by washing with DMF (×6).

*Peptoid synthesis*. Tentagel beads were treated with a solution of chloroacetic acid (0.4 M, 70 μL, 4 equiv.), HOAt (0.4 M, 70 μL, 4 equiv.,), and TMP (0.4 M, 70 μL, 4 equiv.) in DMF at room temperature for 1 hr. After the reaction completion, the beads were washed with DCM (×3) and DMF (×3), and a solution of amine (1M, 280 μL, 40 equiv.) in DMF was added to the resin. The suspension was incubated at 37 °C overnight followed by washing with DMF (×6).

*Reductive amination*. Aldehyde (1 M, 20 equiv.) was incubated with $Cu(OAc)_2$ (0.05 M, 1 equiv.) in 150 μL of 2% AcOH/DMF. The mixture was added to Tentagel beads and incubated with shaking for 2 hrs. Subsequently, a solution of NaBH3CN (1 M, 90 μL in DCM/MeOH (1:1), 12 equiv.) was added to the reaction mixture and incubated at RT overnight, followed by washing with DMF (×6).

**Click reaction of azide-HDNA with substrate beads.** To Tentagel substrate beads (10 μm – 75 μmol), a solution of $CuSO_4$ (173 μL of 500 mM stock; 86.3 μmol), TBTA (15 μL of 10 mM stock; 0.15 μmol), and ascorbic acid (288 μL of 1500 mM stock; 431 μmol) in water was mixed sequentially. DMSO was then added to reach a final water/DMSO ratio of (2:1), and the mixture was shaken at 37 ℃ for 5 min. After that, a solution of azide-HDNA (100 μL of 3 mM stock; 0.3 mmol, 0.004 equiv.), ascorbic acid (2.3 μL of 500 mM stock; 1.15 μmol), and TEAA buffer (473 μL, 200 mM, pH 7.0) in DMSO/water (1:1) was added to the bead suspension. The suspension was shaken at 37 ℃ for 4 hrs. The beads were washed with BTPBB (10 mM Bis-Tris, pH 7.6, 100 mM NaCl, 10 mM EDTA, 1% (m/v) SDS, 1% (v/v) Tween-20) (×3) and incubated with BTPBB (2 mL) overnight. On the next day, the beads were washed again with BTPBB (×3) and BTPWB (×3), then stored in BTPWB at 0 ℃.

**Combinatorial split-and-pool synthesis of DNA-encoded library beads.** Beads loaded with HDNA were first washed with BTPWB (×3) and BTPLB (×3), and incubated in BTPLB for 30 min. After draining, the beads were incubated with FWD primer (90 nmol) and T4 DNA ligase (4000 unit) in BTPLB (2 mL) at room temperature overnight with shaking. The ligated beads were subsequently washed with BTPWB (×3), DMF/water (1:1, ×1), and DMF (×3). The beads were subjected to Fmoc deprotection as described in “standard procedures for DNA-compatible solid-phase synthesis”. For reaction with BB1, the beads were split into 96-well filter plate (pre-wetted with DCM and DMF, Millipore MultiScreen Solvinert 0.45 μm Hydrophobic PTFE), and washed with DMF (×3). Separately, BB1s were pre-activated and transferred to the corresponding wells of the filter plate that contain beads. Building blocks that are Fmoc protected amino acids were reacted with library beads using peptide coupling conditions; amines were reacted with library beads using peptoid synthesis conditions. Upon reaction completion, library beads were washed with DMF (×3), DMF/water (1:1, ×1), BTPWB (×3), and BTPLB (×3), incubated in BTPLB for 30 min, and resuspended in BTPLB (150 μL). Ligation mixture for BB1 containing **[11XX]** (7.5 nmol), **[22XX]** (7.5 nmol), and T4 DNA ligase (600 units) in BTPLB (150 μL) were prepared separately and were transferred to the corresponding wells of the filter plate that contains beads (300 μL total volume per well), and the filter plate was shaken at room temperature overnight. The beads were then washed with BTPWB (x3), DMF/water (1:1, ×1), and DMF (×3). Post ligation, the Fmoc protection groups on library beads were deprotected and washed with DMF (×6).

For the second round of synthesis, library beads were first pooled together and evenly split into 96-well filter plate (pre-wetted with DCM and then DMF), and washed with DMF (×3). BB2s were conjugated to library beads using corresponding chemical conditions described in the previous paragraph. Upon reaction completion, beads were washed with DMF (x3), DMF/water (1:1, ×1), BTPWB (×3), and BTPLB (×3), incubated in BTPLB for 30 min, and resuspended in BTPLB (150 μL). DNA codes (**[13XX], [24XX]**) were ligated to DEL beads following standard procedure. Post ligation, the Fmoc protection groups on library beads were deprotected and washed with DMF (×6).

For the third round of synthesis, library beads were pooled together and evenly split into 96-well filter plate (pre-wetted with DCM and then DMF), and washed with DMF (×3). Corresponding peptide, peptoid synthesis, and reductive amination reactions were performed following DNA-compatible synthetic procedure. After reaction with BB3, DNA codes (**[15XX], [26XX]**) were ligated to library beads.

For ligation of the redundancy codes, beads were pooled and split into 10 wells (pre-wetted with DCM, DMF, water, and then BTPWB), and ligated on the first redundancy code (**[17XX], 10 nmol**). The beads were washed, pooled again and split into 10 wells, followed by ligation of the second redundancy code (**[28XX], 10 nmol**). The beads were then washed with BTPWB (×3) and resuspended in BTPWB. All beads were pooled together and the final ligation with the RVS primer with unique molecular identifier (UMI) (**10 nmol**) were performed. The beads were then washed with BTPWB (×3) and stored in BTPWB at 4 ℃.

**Quality control of solid-phase DNA-encoded library synthesis.** 10 μm quality control beads (TentaGel® RAM) were used in this study instead of the 160 μm beads, because a difference in reactivity of 10 and 160 μm beads were previously observed. QC beads were synthesized in separate wells using procedures identical of library beads synthesis. BBs at each position were randomly selected and reacted with QC beads, followed by ligation of the corresponding DNA codes. After the ligation of redundancy codes and reverse primer, QC beads were collected and checked by PCR followed by sanger sequencing, as well as mass spectrometry.

*Quantitative PCR and gel purification on DNA codes. QC* beads were placed in a well of 384-well RT-qPCR plate (MicroAmp® optical 384-well reaction plate with barcode, Life Technologies TM) and supplemented with forward and reverse primers (0.4 μM) and Power SYBR Green Master Mix (×1 , 20 μL/each well; Life Technologies, Inc.). DNA tags were amplified by thermocycling (50℃ for 2 min; 95℃ for 10 min; 25 cycles of 95℃ for 15 s; and 60℃ for 1 min). Amplicons were purified by native PAGE (10%, 110 V, 60 min) and stained with SYBR Gold following vendor's protocol (Life Technologies, Inc.). PCR products at 160 bp were excised and eluted by tumbling the gel slices in 300 μL of 0.3 M NaCl at 4℃ overnight. The supernatants were passed through glass wool filter followed by addition of ethanol to a final concentration of 70% and 10 μg of glycogen (RNA grade, Thermo Fisher Scientific, Inc.). The solutions were incubated at -20℃ for 2 h and centrifuged at maximum speed for 15 minutes at 4℃. After removing supernatant, the resulting DNA pellet was air dried for 30 minutes, dissolved in 20 μL of nanopure water and sent for Sanger sequencing (Eton Bioscience Inc.).

*Mass spectrometry analysis of QC beads.* QC beads were transferred to 1.6 mL microcentrifuge tubes, washed with H2O (×1) and MeOH (×3), treated with 100 μL of TFA/TIS/H2O (95:2.5:2.5), and dried by air-blowing. Residues were dissolved with 20 μL of MeOH/H2O (1:1) and analyzed on AB SCIEX 4800 Plus MALDI-TOF/TOF (matrix: a-cyano-4-hydroxycinnamic acid).

**Fluorescence-activated cell sorting to identify $(G_4C_2)^{exp}$ ligand.** Library beads were aliquoted to two mobicols (Boca Scientific Inc.) as control and $(G_4C_2)^{exp}$ groups (1M beads for control sample and 20M beads for $r(G_4C_2)^{exp}$ sample). Beads were washed with 6 mL screening buffer x3 (10 mM sodium phosphates, pH 7.0, 100 mM LiCl, 0.05% Tween 20) and equilibrated in 6 mL screening buffer for 30 min with shaking at room temperature. Beads were then incubated in blocking buffer (screening buffer supplemented with 200 nM BSA) and 200 nM bulk yeast tRNA (ThermoFisher Scientific)) for 1 hr with shaking at room temperature. Alexa750-$r(GC)_8$ and Dy647-$r(G_4C_2)_8$ were folded in 1x folding buffer (10 mM sodium phosphates, pH 7.0, 100 mM LiCl) by heating up to 95℃ for 4 minutes and slowly cooling down to room temperature. Both Alexa750-$r(GC)_8$ and Dy647-$r(G_4C_2)_8$ were supplemented to $r(G_4C_2)^{exp}$ sample at final concentrations of 200 nM of 20 nM, respectively. After shaking at room temperature for 2 hrs, control and $r(G_4C_2)^{exp}$

sample were washed with screening buffer ×3 and filtered into 5 mL round-bottom tubes through cell-strainer caps at concentration of 2 × $10^6$ beads/mL.

Samples were sorted using BD FACS Aria3 cell sorter. First, 1× diversity (550,000) beads were recorded for both samples. Samples with high fluorescence signals at both 660 nm (Dy647) and 750 nm (Alexa750) were gated, yielding a hit rate of 0.02%. This gate was then used to sort hit beads in $(G_4C_2)^{exp}$ samples in three technical replicates, each containing 8x library diversity. The sorted beads were collected in 1.6 mL Eppendorf tubes and the DNA codes were amplified via PCR.

**Library preparation and next generation sequencing.** Beads were washed twice with nanopure water, and diluted in 50 µL nanopore water. The first round of PCR amplification was carried out using protocol described in "*Quantitative PCR and gel purification on DNA codes*". The PCR products were resuspended in 20 µL nanopure water and amplified again with NGS adaptor and barcoding primers (**Table S3**). The reaction mixture containing 1× Power SYBR Green Master Mix, 2 µL DNA products, 0.4 µL forward and reverse NGS adaptor primers were thermo-cycled (50℃ for 2 min; 95℃ for 10 min; 25 cycles of 95℃ for 15 s; and 60℃ for 1 min). The PCR products were gel purified and amplicon ~ 225 bp were extracted using QIAEX® II following manufacturer's protocol. The prepared library was sequenced using Illumina Miseq system (122 bp read length).

### COMPUTATIONAL METHODS

**Software and Computational Environment:**

Biopython v1.79 – Structure parsing and manipulation

PyMOL v2.5+ – Structure selection and extraction in headless mode

Open Babel v3.1.1 – Format conversion and charge calculation

AutoDockTools v1.5.7 – Grid and docking parameter generation

AutoGrid v4.2.6 – Grid map calculation

AutoDock-GPU v1.5.3 – GPU-accelerated molecular docking

SLURM v20.11+ – Job scheduling on HPC

Python 3.8+ and Bash 5.0+ were used to execute the pipeline.

**Structural enumeration.** The full DEL and hit population were enumerated using a custom Python script implemented with RDKit.[2] BBs at each position were classified as Fmoc-protected amino acids, amines, diamines, carboxylic acids, or aldehydes. SMILES codes were converted to SMARTS, and reactions were defined based on SMARTS codes. BBs were sequentially reacted based on their functional groups, and the resulting library was converted back to SMILES and exported to Excel.

**Calculation of molecular descriptors**. Molecular structures for Ribo-DEL were provided as SMILES strings, organized into a single-column input file. SMILES strings were parsed into RDKit Mol objects. Canonical RDKit SMILES were generated when requested to ensure consistent representation of molecular structures. For each valid molecule, a complete set of 2D molecular descriptors available in RDKit was computed. Descriptor functions were obtained from the RDKit Descriptors module and executed in a stable, alphabetical order. Each descriptor produced a scalar numeric value (or NaN if unavailable for a given molecule). Any unexpected non-scalar outputs were safely stringified to maintain consistent formatting. Descriptors were calculated one at a time across the full set of molecules to reduce memory overhead. An option to skip descriptors that yielded exclusively missing values (NaN) was included. Results were saved as separate CSV files, one per descriptor, in the designated output directory. Each CSV contained the following fields:

SMILES: the original or canonical SMILES string,

NAME: optional molecule identifier if supplied in the input file, and

Descriptor value: the calculated property for that descriptor.

Descriptor file names were sanitized to ensure compatibility with common file systems. When enabled, a progress bar (via tqdm) reported calculation progress across the descriptor set.

**Chemical space mapping with UMAP.**[3] Molecular structures for the Ribo-DEL, Inforna and an RNA focused library were provided as SMILES strings in three input files, each representing a distinct library. By default, the script reads the first column of each file, but other column indices or delimiters can be specified. Molecules were the transformed into numerical feature vectors using RDKit. Three fingerprinting schemes were supported: Morgan[4] (circular) fingerprints with configurable radius and bit size, RDKit substructure fingerprints,[5] and AtomPair fingerprints.[6] And Morgan fingerprints were used in this study. Fingerprints were computed as binary bit vectors, then converted to NumPy arrays for downstream analysis. To improve computational efficiency and reduce noise, the fingerprint matrices could optionally be preprocessed using Principal Component Analysis (PCA), retaining a specified number of components. The reduced (or raw) fingerprints were then embedded into a two-dimensional space using Uniform Manifold Approximation and Projection (UMAP). All SMILES sets were concatenated and projected jointly to generate a shared embedding space, which was subsequently split back into individual sets for visualization and analysis.

The resulting 2D coordinates for each molecule were combined with their corresponding SMILES strings and dataset labels, and saved to a CSV file. Invalid molecules retained their SMILES strings but were assigned missing coordinate values (NaN). For visualization, scatter plots of the UMAP embeddings were generated with bold, publication-ready styling. Each input set were distinguished using customizable colors, marker shapes, transparencies (alpha), and marker sizes, with corresponding legends. Figures were saved as PNG and/or SVG files.

**NGS decoding.** A custom Python script was used to decode the sequencing results. All reads were filtered by quality, retaining only those with a minimum Phred score above 37. Duplicated reads were removed to avoid PCR bias. A look-up table of all building block combinations and their corresponding DNA codes was generated from a building block code table. The encoding region of each read was then compared against this table. When a match was found, the redundancy code region was inspected, and the redundancy score (number of unique redundancy code combinations) was recorded. Overlapping hits from three technical replicates with redundancy score > 1 were outputted as the final hit population.

**Clustering to generate a representative Ribo-DEL sub-population**. A two-step clustering procedure was employed to efficiently reduce the chemical space of the library. First, the parent library comprising approximately 580,000 compounds was subjected to K-means clustering based on molecular fingerprints to group structurally similar compounds. From these clusters, representative compounds were selected to generate a downsized library of 5,000 molecules. This reduced set preserved the overall chemical diversity of the parent library while enabling more computationally tractable downstream analyses.

**Mordred 2D and 3D descriptor calculations**.[7] Two molecular libraries (5,000-member Ribo-DEL representative and hit population) were provided as plain-text SMILES files with one entry per line (an optional second column for names is ignored by the analysis). For each line, SMILES were parsed with RDKit (SanitizeMol), and explicit hydrogens were added when generating 3D structures. Entries that failed SMILES parsing (e.g., malformed/partial strings) were excluded from downstream steps.

To accelerate geometry preparation at library scale, 3D conformers were generated in parallel using Python's ProcessPoolExecutor (one molecule per worker; up to --n_jobs concurrent workers). For each valid SMILES:

A 3D embedding was produced with RDKit ETKDG v3 (AllChem.ETKDGv3) with a user-settable random seed (--seed) for reproducibility. The embedded structure was relaxed by UFF (AllChem.UFFOptimizeMolecule) for a user-settable number of iterations (--uff_max_iters; default 200). Workers returned MolBlocks (strings) to the main process to avoid inter-process object issues; the main process reconstructed molecules and wrote a single SDF file (structures.sdf) sequentially. Set 1 molecules were written first, followed by set 2, preserving input order. Progress was reported via a tqdm progress bar (stderr) compatible with batch logging. Molecular descriptors were computed with Mordred using its built-in multiprocessing (nproc = --n_jobs). Two panels were calculated:

2D descriptors: computed on sanitized RDKit molecules (no coordinates required).

3D descriptors: computed on the ETKDG+UFF structures written above.

For robustness, descriptor columns with missing values (NaN) or with all zeros across samples were removed prior to analysis. Within each set (set 1 and set 2) and descriptor panel (2D or 3D), features were min–max scaled to [0,1] using MinMaxScaler. The scaled 2D and 3D panels were then concatenated column-wise to form the final feature matrices for set 1 (binders) and set 2 (nonbinders) used in the statistical comparison and visualization.

Note: normalization was applied independently per set to match the behavior of the original analysis script and to facilitate comparison of distribution shapes in plots. For each descriptor present in both sets, we assessed whether the distributions differed between set 1 and set 2. The script supports two tests: Welch's t-test (default; unequal variances) on the scaled values and Mann–Whitney U test (optional; --test_type mannwhitneyu) for a non-parametric alternative. P-values are reported unadjusted and used to rank descriptors by evidence of separation. (If control of the false discovery rate is desired, an FDR procedure such as Benjamini–Hochberg can be applied to the output table externally.) For the top-ranked descriptors (user-defined, --num_properties), the script produces box plots or violin plots (--plot_type) comparing set 1 vs set 2 distributions. Plots are generated with Seaborn/Matplotlib, and statistical annotations (significance stars) are overlaid using statannotations. One image file per descriptor is written to --output_dir, and a CSV table of ranked descriptors with P-values is saved to the path provided as output_file. All heavy steps are parallelized:

Conformer generation: one molecule per worker process (up to --n_jobs).

Descriptor calculation: Mordred’s internal multiprocessing (nproc = --n_jobs).

To avoid thread oversubscription on shared HPC nodes, BLAS/OpenMP threads were limited at runtime (e.g., OMP_NUM_THREADS=1, MKL_NUM_THREADS=1, OPENBLAS_NUM_THREADS=1). The workflow was implemented in Python (≥3.10) using RDKit for cheminformatics and Mordred for descriptors. Reproducibility of 3D structures is ensured by fixing --seed; changing the seed allows probing conformer variability if desired.

Molecules that failed SMILES parsing, 3D embedding, or minimization were omitted from the 3D panel; molecules that produced NaNs in Mordred were excluded on a per-descriptor basis during column filtering.

**Model construction for molecular docking.** An RNA construct containing a 1×1 GG internal loop was modeled based on canonical A-form RNA helical geometry. Standard base-pairing and helical parameters were assigned using NAB module of Amber.[8] To relieve steric clashes and explore energetically favorable conformations of the 1×1 GG loop, simulated annealing was carried out using a molecular dynamics (MD) engine (AMBER).[9] The RNA model was first minimized under positional restraints on the helical regions to preserve global architecture.

The annealing cycle consisted of (1) heating phase: The system was gradually heated from 0 K to 600 K over ~50 ps while applying harmonic restraints to non-loop regions. (2) High-temperature sampling: The structure was propagated at 600 K for ~200 ps to enable sampling of alternative backbone torsions and base stacking conformations in the loop. (3) Cooling phase: The system was slowly cooled back to 300 K over ~200 ps in decrements of 50 K, allowing the loop to settle into low-energy conformations. (4) Final minimization: Energy minimization was performed with reduced restraints to relax bond geometries and optimize local interactions.

Throughout, implicit solvent models were used and counterions were added to neutralize the system.[10] The force field was chosen to ensure accurate representation of RNA conformational energetics (e.g., AMBER ff99bsc0χOL3).[11] From the annealing trajectory, representative low-energy conformations were clustered, and the centroid structure of the most populated cluster was selected.

**Molecular docking implementation**. To prepare RNA-ligand complexes for structure-based docking, a Python-based preprocessing pipeline that processes mmCIF structures was developed. For each RNA-ligand complex, the pipeline performs the following steps. This automated pipeline enables the efficient preparation of large numbers of RNA-ligand complexes for virtual screening.

1. Binding Pocket Definition: A spherical binding pocket was defined as all atoms within a 10 Å radius of the ligand. The selected atoms were extracted using a PyMOL script and saved in a separate file. This pocket was used to define the receptor region for docking.
2. Grid Parameter Calculation: The spatial coordinates of atoms in the binding pocket were used to calculate the grid center and box dimensions. These values were written to a .gpf file for downstream use with AutoDock grid map generation.
3. Format Conversion and Charge Assignment: Both the ligand and the receptor (binding pocket) were converted from mmCIF to PDBQT format using Open Babel (v3.1.1), with Gasteiger partial charges assigned for ligand and Amber partial charges for RNA.
4. Parallel Processing: All processing steps were executed in parallel using Python's multiprocessing module, allowing batch preparation across multiple CPU cores. Output files were organized into individual subdirectories based on the PDB ID.

To perform docking simulations, a companion Bash script that automatically generates and submits SLURM jobs for each RNA-ligand complex prepared in the previous step was developed. This docking pipeline streamlines the execution of GPU-accelerated AutoDock simulations across hundreds of RNA-ligand pairs with minimal manual intervention.[12] The workflow proceeds as follows:

1. Input Parsing and Directory Matching: A CSV file listing complex IDs is provided as input. For each ID, the script performs a case-insensitive search to identify the corresponding directory containing the docking input files.
2. SLURM Job Generation: For each subdirectory, a SLURM job script is created. The job requests one GPU on the appropriate partition, loads required software modules and sets the working directory to the subdirectory.
3. Docking Pipeline, within each SLURM job:
    a. prepare_gpf4.py and prepare_dpf4.py (AutoDockTools) are used to generate grid and docking parameter files, respectively.
    b. autogrid4 computes the receptor grid maps using the .gpf file.
    c. autodock_gpu_64wi (AutoDock-GPU v1.5.3) performs docking simulations using the precomputed grid maps and ligand PDBQT file.
4. Docked Pose Extraction: Docked conformations are extracted from the AutoDock.dlg log file by isolating lines starting with DOCKED. The coordinates are then converted to SDF format using Open Babel to facilitate downstream analysis.
5. Job Submission: Each SLURM script is submitted automatically using sbatch, enabling high throughput docking on an HPC cluster.

**Structure-based RNA model building and dimer design.** Three-dimensional models of the $(G_4C_2)_8$ RNA construct were generated from the primary nucleotide sequence using the FARFAR2 protocol within the Rosetta framework.[13] Starting from the one-dimensional input sequence and secondary structure constraints, FARFAR2 employs fragment assembly combined with Monte Carlo sampling and energy-based scoring to explore conformational space and build candidate 3D structures. The lowest-energy models were clustered to identify representative conformations, and the top-scoring structures were selected as putative models for downstream analysis, including docking and functional annotation.

3D Structures of **11-13** were geometry optimized using the open-source quantum chemistry package Psi4. Initial 3D coordinates were generated from SMILES strings and pre-optimized using force-field methods to provide reasonable starting geometries. Subsequent geometry optimization was carried out at the density functional theory (DFT) level, employing the B3LYP exchange-correlation functional in conjunction with a standard split-valence basis set (e.g., 6-31G*). The B3LYP functional was chosen due to its balance of computational efficiency and accuracy in describing electronic structure and molecular geometries. Convergence criteria for both energy and gradient were set to default Psi4 thresholds to ensure fully optimized minima on the potential energy surface. The resulting optimized geometries were then used to measure the distance between the two nitrogen atoms on the amides or the homopiperazine building blocks.

### IN VITRO METHODS

**Biolayer interferometry (BLI).** BLI studies were performed using either an Octet RED96 system or Octet BLI Discovery 12.2 as previously described.[14] Briefly, 5'-biotinylated nucleic acids (bi-$(G_4C_2)_8$ or bi-$(G_2C_2)_8$) were folded by heating at 95 °C for 4 min in 1× Folding Buffer (10 mM sodium phosphates, pH 7.0 and 100 mM LiCl), followed by slowly cooling to room temperature. To the folded RNA were added Tween-20 (0.05% (v/v) final concentration), and DMSO (1% final concentration) in a final volume of 200 µL per well, with the final concentration of RNA being 0.5 µM. For quenching streptavidin binding sites on the probe, biotin (CAS# 58-85-5) was diluted to 10 µg/mL in assay buffer (10 mM sodium phosphates, pH 7.0, 100 mM LiCl, 0.05% (v/v) Tween-20, 1% DMSO) in a final volume of 200 µL per well. Serial dilutions of the compound of interest were prepared in assay buffer with indicated final concentration. Super streptavidin sensors (SSA, ForteBio, Catalog #: 18-5057,) were equilibrated with assay buffer for 10 min at room temperature in 96-well black microplates (Greiner; Catalog #: 655209). The following time intervals were used during data acquisition (24 °C with shaking at 1000 rpm): baseline step, 60s; loading of RNA, 300s; quenching with biotin, 60s; washing, 100s; association of compound, 120 s; and dissociation of compound, 120 s. The resulting curves were processed and analyzed using Octet Data Analysis software version 12.2. Response from RNA loaded sensors were subtracted by response from no RNA loaded samples (parallel reference) to obtain processed data. For binding affinity measurements, response values at binding equilibrium were calculated by averaging the signal from 105 to 115 s during the association phase. Three technical replicates were combined, and the responses were plotted against compound concentrations using Prism by fitting to equation below:

$$Y = Bmax * X/(Kd + X)$$

where Y is the response at equilibrium and X is the compound concentration.

***In vitro* Chemical Cross-linking and Isolation by Pull-Down (Chem-CLIP) and Competitive Chemical Cross-linking and Isolation by Pull-Down (C-Chem-CLIP).** $r(G_4C_2)_8$ was 5'-end labeled with [g-$^{32}$P] ATP using T4 polynucleotide kinase and gel purified as previously described.[15] The radioactively labeled RNA (~2000 CPM/sample) was folded in 1× Folding Buffer at 95 °C for 4 min and slowly cool to room temperature (20 µL per sample). To folded RNA was added Chem-CLIP probe **2** and control probe **3** to reach a final concentration of 10, 20, 50, and 100µM. For C-Chem-CLIP studies, the parent compound **1** (1, 10, and 100µM) was incubated with RNA for 1 h before the addition of the Chem-CLIP probe **2** (25 µM). Following incubation for 18 h at room temperature, samples were irradiated with UV light (365 nm) for 15 min to allow for diazirine photo-crosslinking. Click reaction mixture (1.0 µL 10 mM biotin azide (Sigma, Catalog #: 762024); 1.0 µL 50 mM HEPES, pH 7; 1.0 µL 10 mM CuSO4; 1.0 µL 50 mM THPTA; 1.0 µL 250 mM sodium ascorbate) were prepared and added to cross-linked RNA. After 2 hrs incubation at 37 °C, 15 µL of Dynabeads™ MyOne™ Streptavidin C1 (ThermoFisher, Catalog #: 65001; pre-washed 3 times with 1× DPBS and resuspended in 1× DPBS) were added to each sample and incubated for 1 hr to allow affinity enrichment of biotin linked RNA. Beads were captured using a magnetic separation rack and washed three times with 1× PBST (DPBS supplemented with 0.1% (v/v) Tween-20). For each sample, the washout and beads were collected separately, and the radioactivity was measured using Beckman Coulter LS6500 Liquid Scintillation Counter.

**NMR Spectroscopy to detect compound binding**. NMR spectra for Carr–Purcell–Meiboom–Gill (CPMG)[16] were acquired on a Bruker Advance III 600 MHz spectrometer equipped with a cryoprobe. All spectra for were processed using TopSpin 4.1.1 (Bruker). The model of the RNA has the following sequence: 5'-GACGGCCGGGUGSSSSCCGGGGCGUC-3' where the GG loop is underlined (purchased from Dharmacon as an HPLC-purified and desalted oligonucleotide). The RNA was diluted in NMR Buffer (5 mM KH2PO4/K2HPO4, 50 mM NaCl, pH 6.0, Temp: 25°C) and folded by heating at 95 ˚C for 3 min followed by snap cooling on ice.

Samples for CPMG experiments contained 5% (v/v) D2O (Cambridge Isotope Labs), 300 µM compound, and 5 mM RNA in a final volume of 600 µL. Experiments were carried out by first collecting spectra of ligand alone at a concentration of 300 mM, followed by addition of folded RNA (5 mM), affording a final ratio of RNA/compound of 60:1. To quantify transverse relaxation times ($T_2$) of RNA-ligand complexes, a pseudo-2D CPMG pulse sequence combined with excitation sculpting-based water suppression, implemented as the cpmg_esgp2d program on a Bruker Avance NMR spectrometer was employed. This sequence enables reliable detection of spin echo decay while effectively suppressing the water signal.

The experiment was configured such that a series of 1D experiments were recorded with incrementally increasing numbers of spin echoes using the default settings (cpmglist). The echo spacing (d20) was chosen to be significantly shorter than 1/J but long enough to accommodate the 180° refocusing pulses, ensuring well-resolved echo formation. The number of echoes in each train (controlled by COUNTER1) was incremented across the indirect dimension (td1), enabling exponential $T_2$ decay curves to be reconstructed for each signal. Water suppression was achieved via excitation sculpting, which uses a pair of shaped 180° pulses bracketed by pulsed field gradients (gp1, gp2). This approach selectively inverts and refocuses water magnetization, while leaving solute signals unaffected. Phase cycling schemes were applied to suppress artifacts and select desired coherence pathways.

### CELLULAR METHODS

**Cell Culture.** HEK293T cells (CRL-3216) were acquired from American Type Culture Collection (ATCC). iPSCs were generated through Answer ALS and obtained from the Cedars Sinai iPSC core.

HEK293T cells were maintained in Dulbecco's Modified Eagle Medium (Corning, Catalog #: 15-017-CI) supplemented with 10% (v/v) fetal bovine serum (Gibco, Catalog #: A56707-01), 1%

penicillin-streptomycin (Corning, Catalog #: 30-002-CI) and 1% glutagro supplement (Corning, Catalog #: 25-015-CI) at 37 °C and 5% $CO_2$.

C9orf72 ALS/FTD patient derived iPSCs (passage number < 18) were maintained in Matrigel (Corning, Catalog #: 356234) coated plates with mTeSR1 feeder-free medium (STEMCELL Technologies; Catalog #: 85850) according to manufacturer's instructions. IPSCs were treated for 5 days in Matrigel-coated plates with mTeSR1 feeder-free medium. Cells were treated in 6-well plates on day 1 and re-dosed on day 3 with compounds in DMSO stock, with 0.1% final DMSO concentration.

**RAN translation assay.** RAN translation assay was completed in HEK293T cells as previously described.[15] Briefly, HEK293T cells were plated in clear-bottom 96-well plate (40,000 cells per well) in FluoroBrite DMEM media (Gibco, Catalog #: A1896702) and incubated overnight. On the second day, cells at 50% confluency were transfected with RANT plasmid ($r(G4C2)_{66}$-No-ATG-eGFP; 0.02 µg per well) and control plasmid (SV40-mCherry; 0.01 µg per well) using Lipofectamine 3000 (Life Technologies, Catalog #: L3000015) per the manufacturer's protocol. After 2 hrs of transfection, compounds in DMSO stocks were added to each well, ensuring the final DMSO concentration was less than 1%. After 24 hrs compounds treatment, eGFP and mCherry signals were measured using an Infinite M1000 Pro Plate Reader (Tecan) with the following settings: (eGFP) excitation: 488 nm; emission: 509 nm; (mCherry) excitation: 587 nm; emission: 610 nm; band width: 5 nm. Fluorescence from eGFP was normalized to mCherry. The resulting fluorescence ratio was normalized to DMSO-treated samples.

**Cellular Chem-CLIP.** Patient-derived iPSCs were plated at $1\times10^6$ cells per well in 6-well plates and allowed to grow into 80% confluency over two days. Cells were then treated with Chem-CLIP probe in growth medium (final concentration of 0.1% (v/v) DMSO) for 12 hrs. After compound incubation, cells were washed with ice cold DPBS (Fisher Scientific, Catalog #: 21-013-CV) twice and subjected to UV crosslinking (with 365 nm light for 10 min), with 1mL DPBS in each well. The total RNA from each sample was then extracted using Quick-RNA Miniprep Kit (Zymo Research, Catalog #: R1055), per the manufacturer's protocol. Crosslinked RNA was clicked on to disulfide agarose azide beads (Click Chemistry Tools, Catalog #: CCT-1238) by adding 15 µL 50 mM HEPES, pH 7.1; 15 µL 10 mM CuSO4; 15 µL 50 mM THPTA; 15 µL 250 mM sodium ascorbate click reaction mixture and incubated at 37 °C for 2 hrs. Upon reaction completion, the beads were washed 3 times with Chem-CLIP Wash Buffer (10 mM Tris-HCl, pH 7.0, 1 mM EDTA, 4 M NaCl, and 0.2% (v/v) Tween-20), and 3 times with DPBS. The pulled-down RNA was eluted from the beads using a 1:1 mixture of 25 µL of TCEP (200 mM) and 25 µL of $K_2CO_3$ (600 mM), with shaking at 37 °C for 30 min, followed by quenching with 400 nM iodoacetamide incubated with shaking at r.t. for 30 min. The supernatant for each sample was collected and purified with RNA Clean XP beads (Beckman Coulter) per the manufacturer's recommended protocol. RNA obtained from Chem-CLIP pull-down was subjected to RT-qPCR for the determination of *C9orf72* mRNA levels as described below.

**RT-qPCR Analysis.** RNA samples were extracted using Quick-RNA Miniprep Kit (Zymo Research) following the manufacturer's protocol. Purified RNA was quantified using Nanodrop UV spectrophotometer. For each sample, 100 ng RNA was reverse transcribed using qScript™ cDNA Synthesis Kit (Quantabio, Catalog #: 95047-100) per the manufacturer's protocol. QPCR was performed using Power SYBR Green Master Mix (Applied Biosystems, Catalog #: 4367659) on a QuantStudio™ Real-Time PCR Instrument (Applied Biosystems). Expression levels of mRNAs were normalized to *b-actin* or *GAPDH*. See **Table S3** for a list of primers.

**Western Blotting to assess RAN translation in c9/ALS patient derived iPSCs.** Upon conclusion of compound treatment, cells were scraped in DPBS and pelleted in to 1.6 mL Eppendorf tubes. Total protein was extracted by incubating the pellet with Co-IP Buffer (50 mM

Tris-HCl, pH 7.4, 300 mM NaCl, 5 mM EDTA, 1% (v/v) Triton-X 100, 2% (w/v) sodium dodecyl sulfate, and 1% (v/v) protease and phosphatase inhibitors) on ice for 5 min followed by sonication (3 s intervals at 35% power for ~40 s) and centrifugation at 14,000 xg for 15 min. Detergent was removed using Pierce Detergent Removal Spin Columns following the manufacturer's protocol. Protein concentration was measured by Pierce BCA Protein Assay Kit.

Approximately 40 µg of total protein was separated on an SDS-polyacrylamide (8%) gel and transferred to a PVDF membrane. The membrane was washed with 1× TBST and then blocked in 5% (w/v) milk in 1× TBST for 1 hr at room temperature. After washing with 1× TBST, primary antibody was added in 1× TBST containing 5% milk overnight at 4 °C. The membrane was washed with 1× TBST and incubated with a 1:3000 dilution of anti- mouse IgG horseradish-peroxidase secondary antibody conjugate (Cell Signaling Technology, Catalog #: 7076S) in 1× TBST containing 5% milk for 1 h at room temperature. The membrane was washed with 1× TBST, and protein expression was quantified using SuperSignal West Pico Plus Chemiluminescent Substrate (Pierce Biotechnology) per the manufacturer's protocol. To quantify β-actin expression, the membrane was stripped using 1× Stripping Buffer (200 mM glycine, pH 2.2 and 0.1% SDS) followed by washing in 1× TBST. The membrane was re-blocked by 5% milk and the antibody incubation processes were repeated.

Antibodies used in this study include: poly(GA) (Millipore, ABN889; 1:1000 dilution); poly(GP) (Millipore, ABN455; 1:1000 dilution); C9orf72 (GeneTex, GTX634482/GTX119776; 1:3000 dilution GAPDH (Cell Signaling Technology, catalog #: #2118; 1:5000 dilution).

### SYNTHETIC METHODS

**Abbreviations.** $CD_3OD$, methanol-d4; DBU, 1,8-diazabicyclo[5.4.0]undec-7-ene; DCM, dichloromethane; DIEA, N,N-diisopropylethylamine; DMAP, 4-dimethylaminopyridine; DMF, N,N-dimethylformamide; DMSO, dimethyl sulfoxide; EDC, N-ethyl-N'-(3-dimethylaminopropyl)carbodiimide hydrochloride; HATU, hexafluorophosphate azabenzotriazole tetramethyl uronium; HOAt, 1-hydroxy-7- azabenzotriazole; HOBt, 1-hydroxybenzotriazole; HRMS, high-resolution mass spectrometry; LC-MS, liquid chromatography-mass spectrometry; MALDI, matrix-assisted laser desorption/ionization; MeOH, methanol; NMR, nuclear magnetic resonance; PYBOP, benzotriazole-1-yloxytripyrrolidinophosphonium hexafluorophosphate; rt, room temperature; TFA, trifluoroacetic acid. s, singlet; d, duplet; t, triplet; q, quartet; quint., quintet; hept., heptet; m, multiplet.

**General.** *Reversed-phase high-performance liquid chromatography (HPLC).* Purification *via* HPLC was carried out using a system of FlexInject (Waters), 1525 Binary HPLC pump (Waters), 2489 UV/Visible detector (Waters), SunFire Prep C18 OBD 5 µm, 19×150 mm column (Waters) and Fraction Collector III (Waters) with the indicated solvent gradients.

*Analytical HPLC.* Compound purity was analyzed either on a 1100 Series HPLC system (Agilent) or on a 1525 Binary Pumps with 2489 UV-Visible Absorption Spectrometer HPLC system (Waters) with indicated gradient, using a SunFire C18 3.5 µm, 4.6×150 mm column (Waters)

*Nuclear magnetic resonance (NMR) spectrometry.* NMR spectra were acquired on a UltraShield 400 MHz spectrometer (Bruker) or Ascend 600 MHz spectrometer (Bruker). $^1H$-NMR spectra were calibrated to residual solvent signal: $\delta$ = 3.31 ppm (quint.) for $CD_3OD$. $^{13}C$-NMR experiments were run with $^1H$-decoupling, and resulting spectra were calibrated to the respective $^{13}C$-D multiplet: $\delta$ = 49.00 ppm (hept.) for $CD_3OD$.

*Mass spectrometry:* High-resolution mass spectrometry data was acquired with an Orbitrap Exploris 120 (Thermo Fisher Scientific) attached to a Vanquish HPLC system (Thermo

Fisher Scientific) in ESI mode. Alternatively, mass spectrometry data was acquired on a 4800 Plus MALDI TOF/TOF Analyzer (Applied Biosystems) with α-cyano-4-hydroxycinnamic acid matrix.

**General protocol for the re-synthesis of DEL hits.** Fmoc-protected Rink-amide resin (100.0 mg, 1 mmol/g, 0.1 mmol, 1 equiv.) was swollen with DMF, deprotected with 2% (v/v) piperidine / 2% (v/v) DBU in DMF (5 mL, 2 × 1 min) and washed with DMF (6 × 5 mL). For peptide coupling, Fmoc-amino acid (0.4 mmol, 4 equiv.), was pre-activated with HATU (0.4 mmol, 4 equiv.) and DIEA (0.8 mmol, 8 equiv.) in DMF (2 mL) for 5 min, followed by addition to the washed beads. The reaction mixture was shaken at r.t. for 2 hrs and washed with DMF (6 × 5 mL). For peptoid formation reaction, BrAcOH (0.3 mmol, 3 equiv.) and DIC (0.3 mmol, 3 equiv.) were pre-mixed in DMF (2 mL), added to the Rink-amide resin and shaken at 37 °C for 0.5 hr. Subsequently, the beads were washed with DMF (3 × 5 mL) and treated with the corresponding amine (2 mmol, 20 equiv.) in 2 mL DMF. The reaction mixture was shaken at 37 °C for 2 hrs and washed with DMF (6 × 5 mL). For reductive amination, aldehyde building block (0.8 mmol, 8 equiv.) and $Cu(OAc)_2$ (0.05 mmol, 0.5 equiv.) in 2% AcOH in DMF (1 mL) was added to the beads and shaken at r.t. for 0.5 hr. After that, NaBH3CN (0.9 mmol, 9 equiv.) in DCM/MeOH (1:1, 1 mL) was added to the beads and shaken at r.t. overnight, then washed with DMF (6 × 5 mL). For compound cleavage, the reacted beads were washed with MeOH (1 × 5 mL) and DCM (3 × 5 mL), and treated with 20% TFA in DCM (3 mL) at r.t. for 2 hrs. The resulting solution was dried by air blowing and purified by HPLC using indicated gradient to afford the target product.

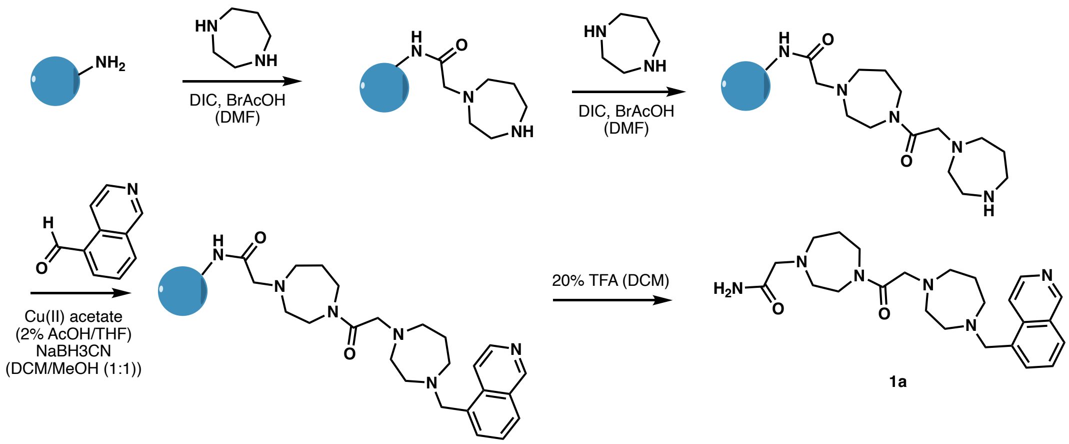


**1a**

**2-(4-(2-(4-(Isoquinolin-5-ylmethyl)-1,4-diazepan-1-yl)acetyl)-1,4-diazepan-1-yl)acetamide (1a).** Homopiperazine (BB1 and BB2) and Isoquinoline-5-carbaldehyde (BB3) were used in peptoid synthesis and reductive amination, respectively, following the general protocol. The cleavage product was purified by HPLC with gradient of 0-50% MeOH in $H_2O$ (0.1% TFA) over 60 min, providing compound **1a** as a yellow oil. $^1$H NMR (400 MHz, $CD_3OD$) δ 9.73 (s, 1H), 8.79 (dd, *J* = 6.7, 0.9 Hz, 1H), 8.62 (dd, *J* = 6.7, 0.8 Hz, 1H), 8.45 (d, *J* = 8.4 Hz, 1H), 8.22 (dd, *J* = 7.2, 1.2 Hz, 1H), 7.99 (dd, *J* = 8.4, 7.1 Hz, 1H), 4.42 (s, 2H), 4.31 (d, *J* = 2.4 Hz, 2H), 4.04 (d, *J* = 12.7 Hz, 2H), 3.98 – 3.77 (m, 2H), 3.77 – 3.41 (m, 10H), 3.25 – 3.18 (m, 2H), 3.04 (t, *J* = 5.8 Hz, 2H), 2.38 – 2.06 (m, 4H). $^{13}$C NMR (150 MHz, $CD_3OD$) δ 167.53, 166.49, 149.77, 139.18, 138.80, 138.77, 134.69, 131.75, 131.22, 129.92, 123.22, 59.74, 58.61, 58.33, 56.33, 55.96, 55.89, 55.45, 55.06, 50.55, 45.91, 41.42, 25.28, 24.79. (Note that two sets of peaks corresponding to two different conformations were found, and only the dominant set was reported). HRMS-ESI (m/z): calcd for $C_{24}H_{34}N_6O_2$ $[M + H]^+$: 439.2816; found: 439.2827.

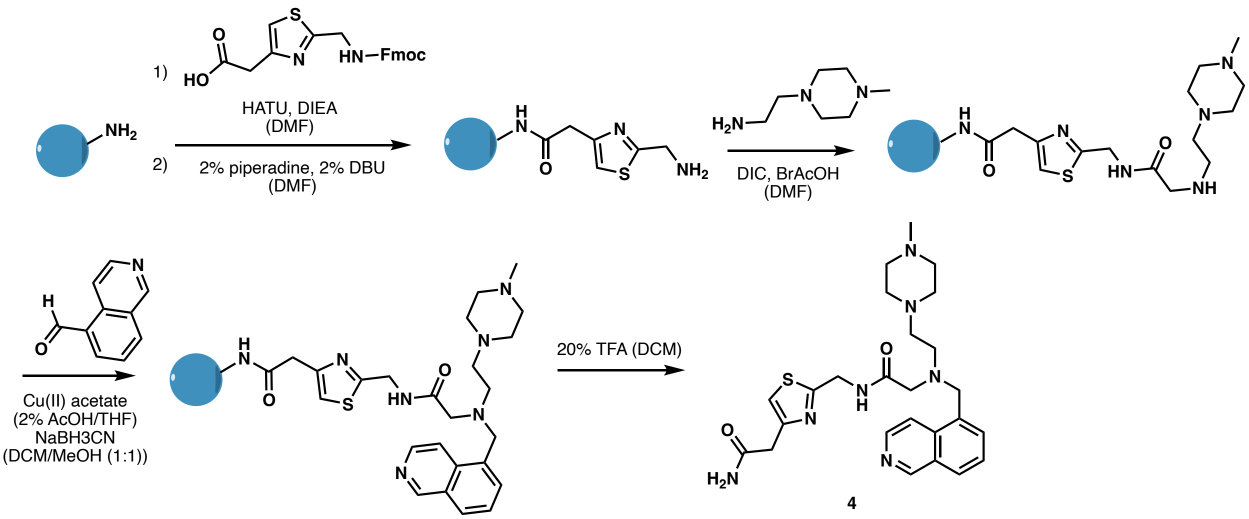


4

**$N$-((4-(2-Amino-2-oxoethyl)thiazol-2-yl)methyl)-2-((isoquinolin-5-ylmethyl)(2-(4-methylpiperazin-1-yl)ethyl)amino)acetamide (4).** Fmoc-aminomehtly-2-thiazole-4-acetic-OH (BB1), *N*-methyl-*N′*-piperazin-2-ethylamine (BB2), and Isoquinoline-5-carbaldehyde (BB3) were used in peptide, peptoid synthesis, and reductive amination, following the general protocol. The cleavage product was purified by HPLC with gradient of 0-80% MeOH in $H_2O$ (0.1% TFA) over 60 min, providing compound **4** as a yellow oil. $^1$H NMR (400 MHz, $CD_3OD$) δ 9.71 (s, 1H), 8.68 (d, $J$ = 6.6 Hz, 1H), 8.62 (d, $J$ = 6.6 Hz, 1H), 8.42 (d, $J$ = 8.3 Hz, 1H), 8.22 (d, $J$ = 7.1 Hz, 1H), 7.97 (dd, $J$ = 8.3, 7.2 Hz, 1H), 7.29 (s, 1H), 4.55 (d, $J$ = 4.3 Hz, 4H), 3.69 (s, 2H), 3.61 (s, 2H), 3.35 (m, 4H), 3.14 (dt, $J$ = 35.6, 5.5 Hz, 8H), 2.89 (s, 3H).

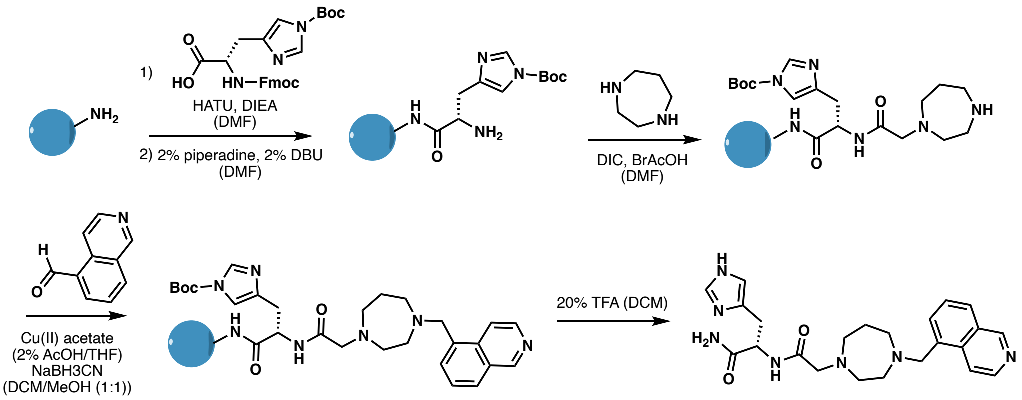


5

**(S)-3-(1*H*-Imidazol-4-yl)-2-(2-(4-(isoquinolin-5-ylmethyl)-1,4-diazepan-1-yl)acetamido)propenamide (5).** Fmoc-His(Boc)-OH (BB1), Homopiperazine (BB2), and Isoquinoline-5-carbaldehyde (BB3) were used in peptide, peptoid synthesis and reductive amination, respectively, following the general protocol. The cleavage product was purified by HPLC with gradient of 0-80% MeOH in $H_2O$ (0.1% TFA) over 60 min, providing compound **5** as a yellow oil. $^1$H NMR (400 MHz, $CD_3OD$) δ 9.70 (s, 1H), 8.82 (dd, *J* = 3.3, 1.4 Hz, 1H), 8.71 (d, *J* = 3.6 Hz, 1H), 8.62 (d, *J* = 6.6 Hz, 1H), 8.44 (d, *J* = 8.4 Hz, 1H), 8.21 (d, *J* = 7.2 Hz, 1H), 7.97 (t, *J* = 7.8 Hz, 1H), 7.37 (dd, *J* = 3.4, 1.4 Hz, 1H), 4.74 (dd, *J* = 7.8, 5.7 Hz, 1H), 4.48 (d, *J* = 3.5 Hz, 2H), 3.89 (dd, *J* = 6.3, 3.3 Hz, 2H), 3.34 (d, *J* = 6.0 Hz, 4H), 3.31 – 3.05 (m, 6H), 2.14 – 2.06 (m, 2H).

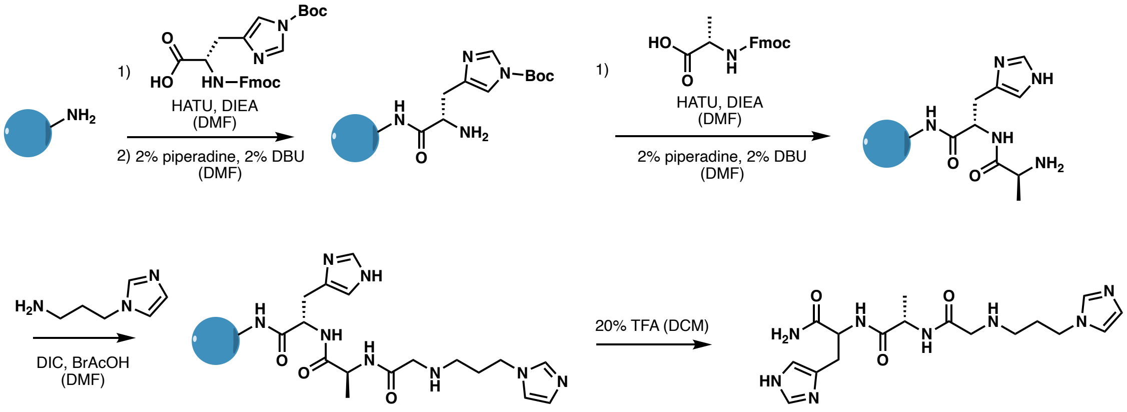

6

**(2*S*)-2-(2-((3-(1*H*-Imidazol-1-yl)propyl)amino)acetamido)-*N*-(1-amino-3-(1*H*-imidazol-4-yl)-1-oxopropan-2-yl)propenamide (6).** Fmoc-His(Boc)-OH (BB1), Fmoc-Ala-OH (BB2) and Isoquinoline-5-carbaldehyde (BB3) were used in peptide synthesis and reductive amination, following the general protocol. The cleavage product was purified by HPLC with gradient of 0-60% MeOH in $H_2O$ (0.1% TFA) over 60 min, providing compound **6** as a yellow oil. $^1$H NMR (400 MHz, $CD_3OD$) δ 8.97 (d, *J* = 1.7 Hz, 1H), 8.80 (d, *J* = 1.4 Hz, 1H), 7.68 (q, *J* = 1.8 Hz, 1H), 7.61 (d, *J* = 1.1 Hz, 1H), 7.39 (dd, *J* = 6.4, 1.4 Hz, 1H), 4.75 (dd, *J* = 8.4, 4.9 Hz, 1H), 4.69 (dd, *J* = 8.3, 5.2 Hz, 1H), 4.40 (t, *J* = 7.2 Hz, 2H), 4.30 (dq, *J* = 14.5, 7.2 Hz, 1H), 3.89 (d, *J* = 5.8 Hz, 2H), 3.20 – 3.06 (m, 3H), 2.39 – 2.28 (m, 2H), 1.36 (t, *J* = 7.3 Hz, 3H).

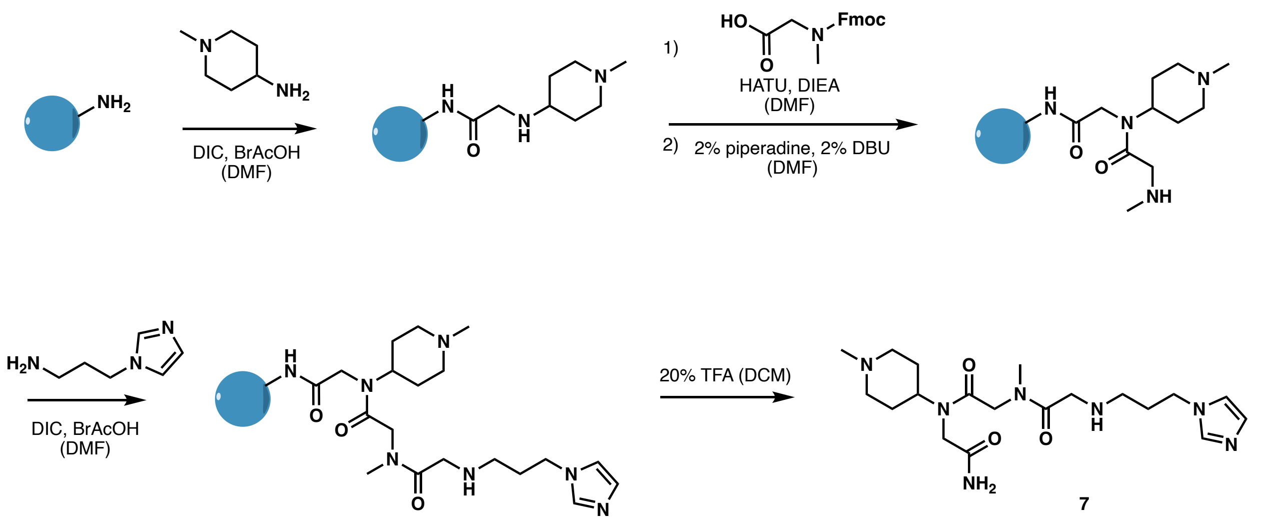

**2-((3-(1*H*-Imidazol-1-yl)propyl)amino)-*N*-(2-((2-amino-2-oxoethyl)(1-methylpiperidin-4-yl)amino)-2-oxoethyl)-*N*-methylacetamide (7).** The 1-methylpiperidin-4-amine (BB1), *N*-(((9*H*-fluoren-9-yl)methoxy)carbonyl)-*N*-methylglycine (BB2), and *3-(1H-imidazol-1-yl)propan-1-amine* (BB3) were used in peptoid, peptide synthesis and reductive amination following the general protocol. The cleavage product was purified by HPLC with gradient of 0-80% MeOH in $H_2O$ (0.1% TFA) over 60 min, providing compound **7** as a colorless oil. $^1$H NMR (400 MHz, $CD_3OD$) δ 9.00 (s, 1H), 7.70 (s, 1H), 7.62 (s, 1H), 4.41 (t, $J$ = 7.2 Hz, 2H), 4.19 (dd, $J$ = 10.7, 2.0 Hz, 2H), 4.16 – 4.05 (m, 4H), 4.05 – 3.92 (m, 2H), 3.88 – 3.77 (m, 1H), 3.76 – 3.62 (m, 1H), 3.58 – 3.46 (m, 1H), 3.35 (d, $J$ = 2.3 Hz, 2H), 3.19 – 3.07 (m, 2H), 3.04 (s, 2H), 2.95 (s, 1H), 2.35 (p, $J$ = 7.4 Hz, 2H), 2.17 – 1.97 (m, 4H).

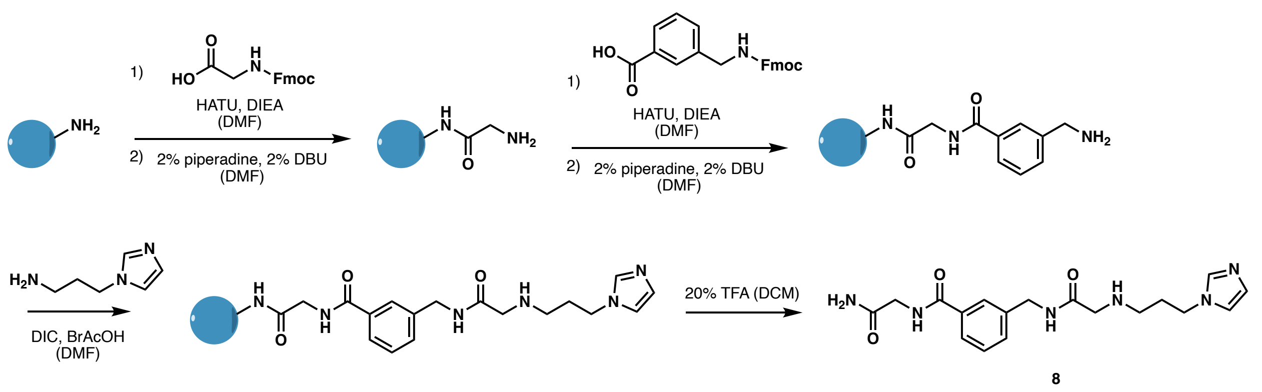

**3-((2-((3-(1*H*-Imidazol-1-yl)propyl)amino)acetamido)methyl)-*N*-(2-amino-2-oxoethyl)benzamide (8).** Fmoc-Gly-OH (BB1), Fmoc-3-aminoMe-Bz-OH (BB2), and *N*-imidazole-3-propylamine (BB3) were used in peptide synthesis and reductive amination following the general protocol. The cleavage product was purified by HPLC with gradient of 0-80% MeOH in $H_2O$ (0.1% TFA) over 60 min, providing compound **8** as a colorless oil. $^1$H NMR (400 MHz, $CD_3OD$) δ 8.94 (s, 1H), 7.82 (d, *J* = 1.9 Hz, 1H), 7.77 (t, *J* = 7.3 Hz, 1H), 7.66 (s, 1H), 7.58 (s, 1H), 7.50 (d, *J* = 7.6 Hz, 1H), 7.45 (t, *J* = 7.6 Hz, 1H), 4.50 (s, 2H), 4.40 (t, *J* = 7.1 Hz, 2H), 4.11 – 4.02 (m, 2H), 3.91 (s, 2H), 3.17 – 3.08 (m, 2H), 2.32 (p, *J* = 7.2 Hz, 2H).

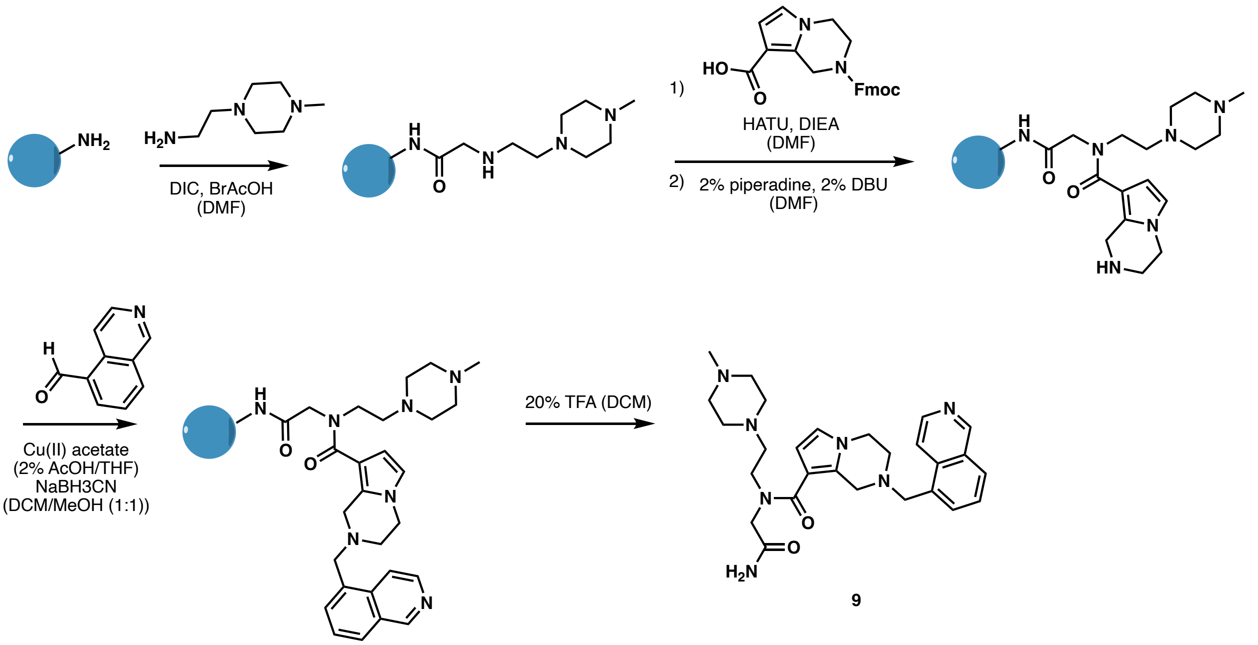


***N*-(2-Amino-2-oxoethyl)-2-(isoquinolin-5-ylmethyl)-*N*-(2-(4-methylpiperazin-1-yl)ethyl)-1,2,3,4-tetrahydropyrrolo[1,2-a]pyrazine-8-carboxamide (9).** *N*-methyl-*N'*-piperazin-2-ethylamine (BB1), Fmoc-tetrahydropyrrolopyrazine-OH (BB2), and Isoquinoline-5-carbaldehyde (BB3) were used in peptoid synthesis and reductive amination following the general protocol. The cleavage product was purified by HPLC with gradient of 0-80% MeOH in $H_2O$ (0.1% TFA) over 60 min, providing compound **9** as a yellow oil. $^1$H NMR (400 MHz, $CD_3OD$) δ 9.79 (s, 1H), 8.82 (d, *J* = 6.8 Hz, 1H), 8.65 (d, *J* = 6.7 Hz, 1H), 8.55 (d, *J* = 8.5 Hz, 1H), 8.40 – 8.29 (m, 1H), 8.12 –

8.00 (m, 1H), 6.75 (d, *J* = 2.9 Hz, 1H), 6.26 (s, 1H), 4.75 (d, *J* = 10.2 Hz, 2H), 4.32 (d, *J* = 12.8 Hz, 4H), 4.18 (d, *J* = 5.5 Hz, 2H), 3.82 (s, 2H), 3.43 (d, *J* = 62.4 Hz, 12H), 2.94 (d, *J* = 2.8 Hz, 3H).

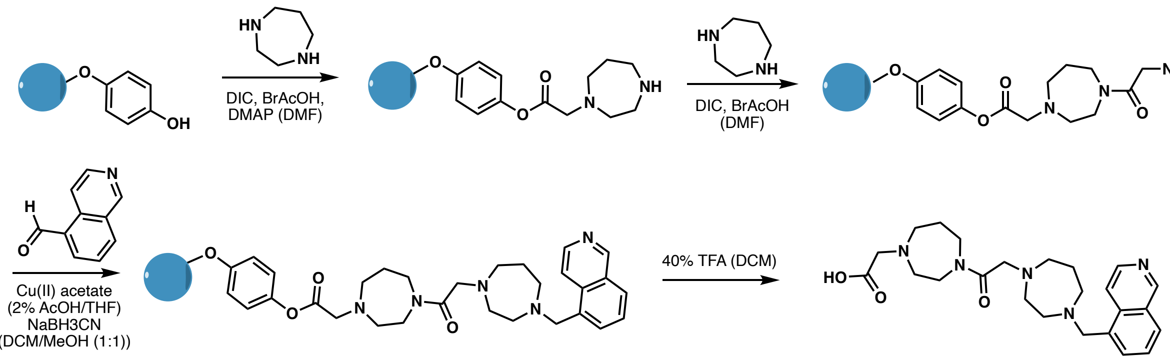


**2-(4-(2-(4-(Isoquinolin-5-ylmethyl)-1,4-diazepan-1-yl)acetyl)-1,4-diazepan-1-yl)acetic acid (1-COOH).** Wang resin (100.0 mg, 0.08 mmol, 1 equiv.) was swollen with DMF. A solution of BrAcOH (33.3 mg, 0.24 mmol, 3 equiv.), DMAP (2.0 mg, 0.016 mmol, 0.2 equiv.), and DIC (30.3 mg, 0.24 mmol, 3 equiv.) in DMF (2 mL) were reacted with the beads at 37 °C for 2 hrs and washed with DMF (6 × 5 mL). Homopiperazine (BB1 and BB2, 160.0 mg, 1.6 mmol, 20 equiv.) in 2 mL DMF were added to the drained beads and reacted at 37 °C for 2 hrs. After washing with DMF (6 × 5 mL), the peptoid synthesis procedure was repeated without the addition of DMAP. Finally, a mixture of isoquinoline-5-carbaldehyde (BB3) (100.0 mg, 0.64 mmol, 8 equiv.) and $Cu(OAc)_2$ (7.3 mg, 0.04 mmol, 0.5 equiv.) in 2% AcOH in DMF (1 mL) was added to the beads and shaken at r.t. for 0.5 hr. After that, NaBH3CN (45.2 mg, 0.72 mmol, 9 equiv.) in DCM/MeOH (1:1, 1 mL) was added to the beads and shaken at r.t. overnight, then washed with DMF (6 × 5 mL). For compound cleavage, the reacted beads were washed with MeOH (1 × 5 mL) and DCM (3 × 5 mL), and treated with 40% TFA in DCM (3 mL) at r.t. for 2 hrs. The resulting solution was dried by air blowing and purified by HPLC with the following gradient: 0-50% MeOH in $H_2O$ (0.1% TFA) over 60 min, providing compound **1-COOH** as a yellow oil. $^1$H NMR (600 MHz, $CD_3OD$) δ 9.21 (s, 1H), 8.48 – 8.43 (m, 1H), 8.26 (d, *J* = 6.0 Hz, 1H), 8.02 (d, *J* = 8.2 Hz, 1H), 7.75 (dd, *J* = 7.0, 4.4 Hz, 1H), 7.63 (t, *J* = 7.6 Hz, 1H), 4.06 (s, 2H), 3.63 (tt, *J* = 33.2, 5.5 Hz, 4H), 3.39 (d, *J* =

14.4 Hz, 2H), 3.15 (d, *J* = 6.9 Hz, 2H), 2.87 (dd, *J* = 6.3, 3.8 Hz, 1H), 2.80 (q, *J* = 5.9 Hz, 5H), 2.75 (qd, *J* = 6.1, 3.4 Hz, 5H), 1.99 – 1.77 (m, 4H). $^{13}$C NMR (150 MHz, $CD_3OD$), δ 177.81, 177.45, 172.21, 172.12, 153.59, 142.62, 136.88, 136.07, 133.36, 130.65, 128.78, 128.38, 119.89, 63.35, 63.11, 61.09, 60.84, 60.75, 57.51, 56.76, 56.43, 56.10, 56.01, 55.66, 55.55, 55.54, 55.49, 55.40, 55.31, 55.28, 48.19, 47.34, 45.89, 28.62, 28.51, 28.35, 27.72. (Note that two sets of peaks corresponding to two different conformations were found). HRMS-ESI (m/z): calcd for $C_{24}H_{33}N_5O_3$ $[M + H]^+$: 440.2656; found: 440.2655.

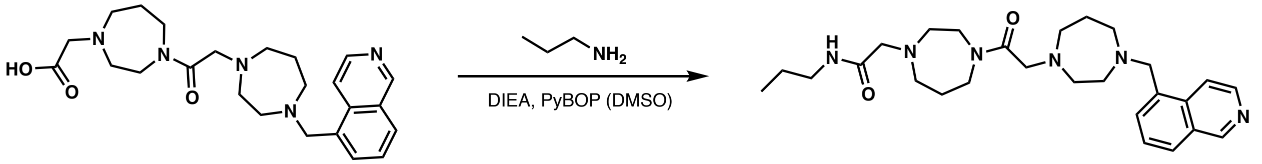


1-COOH

1

**2-(4-(2-(4-(Isoquinolin-5-ylmethyl)-1,4-diazepan-1-yl)acetyl)-1,4-diazepan-1-yl)-*N*-propylacetamide (1).** To a solution of **1-COOH** (4.4 mg, 0.01 mmol, 1 equiv.) in DMSO (50 µL) was added PyBOP (10.4 mg, 0.02 mmol, 2 equiv.) and DIEA (5.2 mg, 0.04 mmol, 4 equiv.). After 30 min, a solution of propylamine (1.2 mg, 0.02 mmol, 2 equiv.) in DMSO (20 µL) was added and the reaction mixture was stirred at r.t. for 2 hrs. The reaction mixture was diluted in 10% MeOH in $H_2O$ (4 mL) and purified using HPLC with a gradient of 0-50% MeOH in $H_2O$ (0.1% TFA) over 60 min, yielding compound **1** as a yellow oil. $^1$H NMR (400 MHz, $CD_3OD$) δ 9.72 (s, 1H), 8.78 (d, *J* = 6.7 Hz, 1H), 8.61 (d, *J* = 6.6 Hz, 1H), 8.44 (d, *J* = 8.3 Hz, 1H), 8.21 (d, *J* = 7.1 Hz, 1H), 8.03 – 7.92 (m, 1H), 4.40 (s, 2H), 4.30 (s, 2H), 4.00 (d, *J* = 11.7 Hz, 2H), 3.95 – 3.69 (m, 2H), 3.70 – 3.38 (m, 10H), 3.25 – 3.10 (m, 4H), 3.02 (dd, *J* = 7.1, 4.5 Hz, 2H), 2.21 (d, *J* = 54.2 Hz, 4H), 1.55 (h, *J* = 7.4 Hz, 2H), 0.94 (t, *J* = 7.4 Hz, 3H). $^{13}$C NMR (151 MHz, $CD_3OD$) δ 166.46, 165.16, 149.67, 139.24, 138.86, 134.44, 131.78, 131.30, 129.90, 123.32, 59.75, 58.62, 58.58, 56.31, 56.05, 55.91, 55.41, 55.08, 50.56, 42.36, 41.42, 25.30, 24.83, 23.47, 11.65. HRMS-ESI (m/z): calcd for $C_{27}H_{40}N_6O_2$ $[M + H]^+$: 481.3286; found: 481.3288.

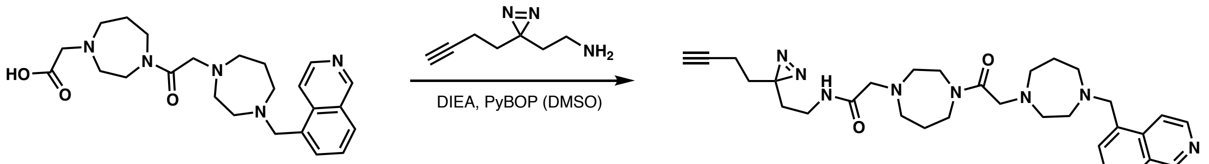


***N*-(2-(3-(But-3-yn-1-yl)-3*H*-diazirin-3-yl)ethyl)-2-(4-(2-(4-(isoquinolin-5-ylmethyl)-1,4-diazepan-1-yl)acetyl)-1,4-diazepan-1-yl)acetamide (2)** To a solution of **1-COOH** (4.4 mg, 0.01 mmol, 1 equiv.) in DMSO (50 µL) was added PyBOP (10.4 mg, 0.02 mmol, 2 equiv.) and DIEA (5.2 mg, 0.04 mmol, 4 equiv.). After 30 min, a solution of 2-(3-(but-3-yn-1-yl)-3*H*-diazirin-3-yl)ethan-1-amine (2.7 mg, 0.02 mmol, 2 equiv.) in DMSO (20 µL) was added and the reaction mixture was stirred at r.t. for 2 hrs. The reaction mixture was diluted in 10% MeOH in $H_2O$ (4 mL) and purified using HPLC with a gradient of 0-50% MeOH in $H_2O$ (0.1% TFA) over 60 min, yielding compound **2** as a yellow oil. $^1$H NMR (600 MHz, $CD_3OD$) δ 9.74 (s, 1H), 8.80 (d, *J* = 6.7 Hz, 1H), 8.62 (d, *J* = 6.7 Hz, 1H), 8.46 (d, *J* = 8.3 Hz, 1H), 8.23 (dd, *J* = 7.2, 1.2 Hz, 1H), 7.99 (dd, *J* = 8.3, 7.1 Hz, 1H), 4.42 (s, 2H), 4.32 (s, 2H), 4.01 (d, *J* = 17.0 Hz, 2H), 3.83 – 3.44 (m, 12H), 3.21 (t, *J* = 5.0 Hz, 2H), 3.16 (t, *J* = 7.0 Hz, 2H), 3.04 (t, *J* = 5.8 Hz, 2H), 2.33 – 2.11 (m, 5H), 2.03 (tt, *J* = 7.4, 2.3 Hz, 2H), 1.71 – 1.61 (m, 4H). $^{13}$C NMR (150 MHz, $CD_3OD$) δ 166.38, 165.27, 149.51, 139.36, 139.16, 134.64, 134.12, 131.95, 131.42, 129.88, 123.48, 83.57, 70.45, 59.66, 58.63, 58.59, 56.30, 55.89, 55.41, 55.09, 50.50, 45.93, 41.43, 35.55, 33.23, 33.06, 27.88, 25.30, 24.68, 13.82. HRMS-ESI (m/z): calcd for $C_{31}H_{42}N_8O_2$ $[M + H]^+$: 559.3504; found: 559.3502.

**Dimeric compound synthesis.**

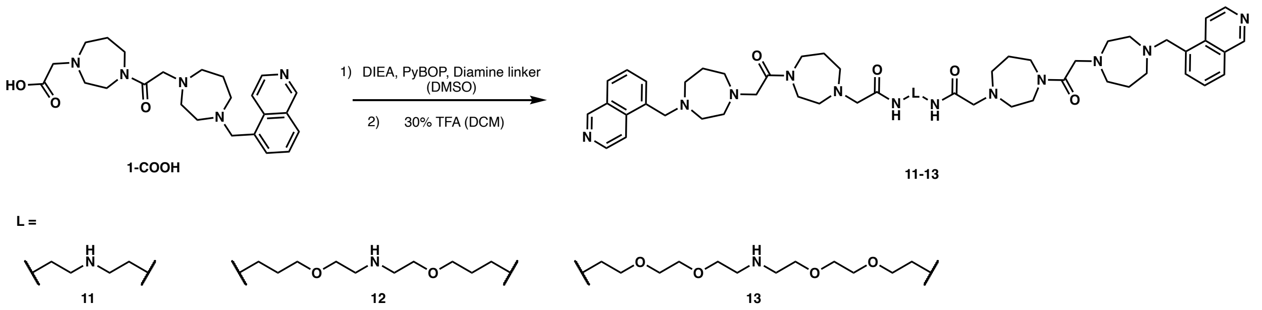


**General.** To a solution of **1-COOH** (9.7 mg, 0.022 mmol, 1 equiv.) in DMF (100 μL) was added PyBOP (22.9 mg, 0.044 mmol, 2 equiv.) and DIEA (11.4 mg, 0.088 mmol, 4 equiv.). After 30 min, a solution of diamine linker (0.01 mmol, 0.45 equiv.) in DMF was added, and the reaction mixture was stirred at r.t. overnight. The reaction mixture was diluted in 10% MeOH in $H_2O$ (4 mL) and purified using HPLC with a gradient of 0-60% MeOH in $H_2O$ (0.1% TFA) over 60 min. A solution of 40% TFA in DCM was added to the dried product and stirred for 2 hrs. And the deprotected compound was purified by HPLC with a gradient of 0-60% MeOH in $H_2O$ (0.1% TFA) over 60 min. The same synthetic method was used to produce compound **11-13**.

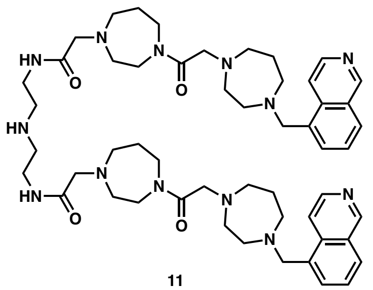

11

***N*,*N'*-(Azanediylbis(ethane-2,1-diyl))bis(2-(4-(2-(4-(isoquinolin-5-ylmethyl)-1,4-diazepan-1-yl)acetyl)-1,4-diazepan-1-yl)acetamide) (11).** Compound **11** was synthesized following the general procedure, using *tert*-butyl bis(2-aminoethyl)carbamate (Aaron Chemicals, Catalog #: AR00IAM8) as the diamine linker. $^1$H NMR (600 MHz, $CD_3OD$) δ 9.75 (s, 2H), 8.81 (d, *J* = 6.7 Hz, 2H), 8.67 – 8.58 (m, 2H), 8.49 (d, *J* = 8.2 Hz, 2H), 8.28 (d, *J* = 6.6 Hz, 2H), 8.02 (dd, *J* = 8.3, 7.1

Hz, 2H), 4.51 (s, 4H), 4.33 (s, 4H), 4.06 (d, $J$ = 13.9 Hz, 4H), 3.98 – 3.42 (m, 28H), 3.32 (d, $J$ = 2.9 Hz, 4H), 3.23 (td, $J$ = 6.0, 1.7 Hz, 4H), 3.12 (t, $J$ = 5.7 Hz, 4H), 2.34 – 2.27 (m, 3H), 2.18 (dq, $J$ = 10.8, 5.0 Hz, 5H). $^{13}$C NMR (151 MHz, $CD_3OD$) δ 166.94, 166.40, 149.40, 139.90, 139.55, 133.92, 133.82, 132.47, 131.69, 129.91, 123.64, 59.43, 58.74, 58.59, 56.62, 55.96, 55.75, 55.33, 55.18, 50.58, 48.32, 46.10, 41.53, 37.04, 25.30, 24.37. HRMS-ESI (m/z): calcd for $C_{52}H_{75}N_{13}O_4$ $[M + 2H]^{2+}$: 473.8106; found: 473.8105.

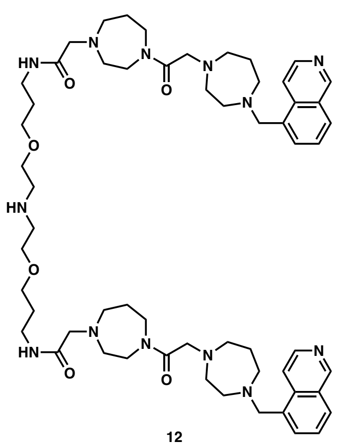

12

***N,N'*-(((Azanediylbis(ethane-2,1-diyl))bis(oxy))bis(propane-3,1-diyl))bis(2-(4-(2-(4-(isoquinolin-5-ylmethyl)-1,4-diazepan-1-yl)acetyl)-1,4-diazepan-1-yl)acetamide) (12).** Compound **12** was synthesized following the general procedure, using *tert-butyl bis(2-(3-aminopropoxy)ethyl)carbamate* as the diamine linker. $^{1}$H NMR (600 MHz, $CD_3OD$) δ 9.75 (s, 2H), 8.81 (d, $J$ = 6.7 Hz, 2H), 8.63 (d, $J$ = 6.7 Hz, 2H), 8.49 (d, $J$ = 8.3 Hz, 2H), 8.27 (d, $J$ = 6.6 Hz, 2H), 8.04 – 7.99 (m, 2H), 4.50 (s, 4H), 4.33 (s, 4H), 4.03 (d, $J$ = 17.9 Hz, 4H), 3.97 – 3.43 (m, 32H), 3.35 (d, $J$ = 6.8 Hz, 4H), 3.32 (m, 4H), 3.28 (t, $J$ = 5.1 Hz, 4H), 2.36 – 2.26 (m, 3H), 2.23 – 2.13 (m, 5H), 1.81 (p, $J$ = 6.6 Hz, 4H). $^{13}$C NMR (150 MHz, $CD_3OD$) δ 166.43, 165.62, 149.40, 139.85, 139.55, 133.91, 132.43, 131.69, 129.91, 123.65, 69.52, 66.55, 59.47, 58.74, 58.65, 56.52, 55.98, 55.57, 55.41, 55.18, 50.61, 48.36, 46.06, 41.49, 37.63, 30.35, 25.26, 24.42. HRMS-ESI (m/z): calcd for $C_{58}H_{87}N_{13}O_6$ $[M + 2H]^{2+}$: 531.8524; found: 531.8525.

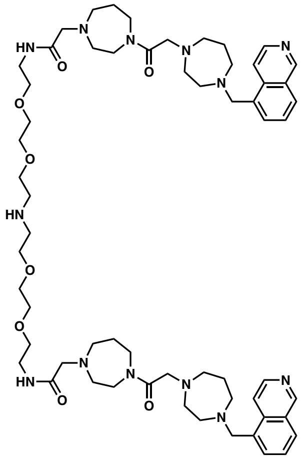

13

***N,N'*-(3,6,12,15-Tetraoxa-9-azaheptadecane-1,17-diyl)bis(2-(4-(2-(4-(isoquinolin-5-ylmethyl)-1,4-diazepan-1-yl)acetyl)-1,4-diazepan-1-yl)acetamide) (13).** Compound **13** was synthesized following the general procedure, using *tert-butyl bis(2-(2-(2-aminoethoxy)ethoxy)ethyl)carbamate*. $^{1}$H NMR (400 MHz, $CD_3OD$) δ 9.71 (s, 2H), 8.77 (d, *J* = 6.7 Hz, 2H), 8.61 (d, *J* = 6.7 Hz, 2H), 8.45 (d, *J* = 8.4 Hz, 2H), 8.22 (d, *J* = 7.1 Hz, 2H), 8.03 – 7.95 (m, 2H), 4.43 (s, 4H), 4.30 (s, 4H), 4.03 (d, *J* = 9.6 Hz, 4H), 3.97 – 3.71 (m, 8H), 3.67 (t, *J* = 1.3 Hz, 8H), 3.63 – 3.39 (m, 28H), 3.27 (t, *J* = 5.0 Hz, 4H), 3.22 (s, 4H), 3.05 (d, *J* = 5.8 Hz, 4H), 2.28 (d, *J* = 6.5 Hz, 3H), 2.15 (d, *J* = 6.1 Hz, 5H). $^{13}$C NMR (150 MHz, $CD_3OD$) δ 166.46, 165.65, 149.53, 139.23, 139.05, 134.41, 134.31, 131.89, 131.36, 129.84, 123.31, 71.27, 71.17, 70.23, 66.76, 59.62, 58.63, 58.59, 56.37, 55.93, 55.89, 55.46, 55.03, 50.58, 48.31, 45.95, 41.43, 40.32, 25.22, 24.66. HRMS-ESI (m/z): calcd for $C_{60}H_{91}N_{13}O_8$ $[M + 2H]^{2+}$: 561.8630; found: 561.8630.

**Compound 1 structural expansion via peptide coupling.**

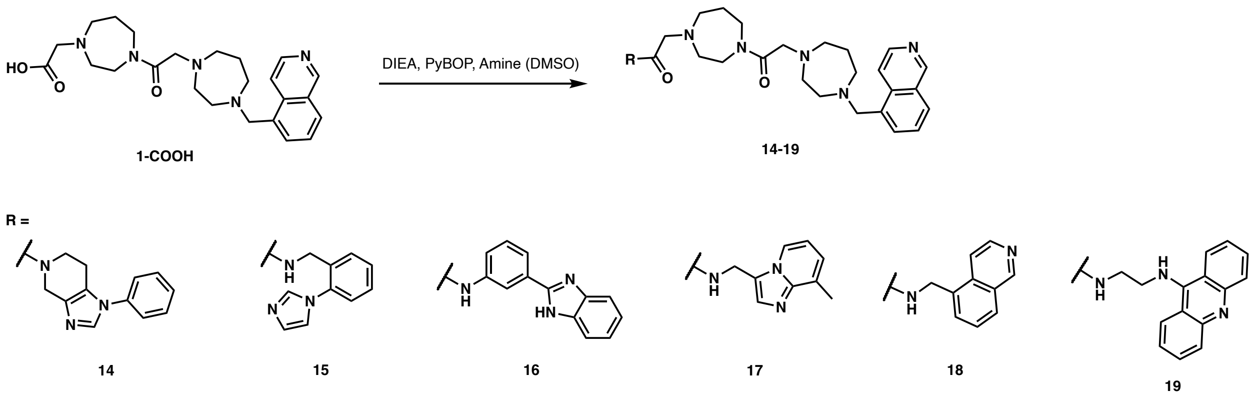


**General.** To a solution of **1-COOH** (8.8 mg, 0.02 mmol, 1 equiv.) in DMF (100 μL) was added PyBOP (41.6 mg, 0.08 mmol, 4 equiv.) and DIEA (20.7 mg, 0.16 mmol, 8 equiv.). After 30 min, a solution of corresponding amine (0.08 mmol, 4 equiv.) in DMF was added, and the reaction mixture was stirred at r.t. overnight. The reaction mixture was then diluted in 10% MeOH in $H_2O$ (4 mL) and purified using HPLC with a gradient of 0-60% MeOH in $H_2O$ (0.1% TFA) over 60 min. The same synthetic method was used to produce compound **14-19**.

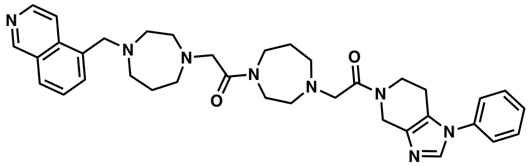

14

**2-(4-(Isoquinolin-5-ylmethyl)-1,4-diazepan-1-yl)-1-(4-(2-oxo-2-(1-phenyl-1,4,6,7-tetrahydro-5*H*-imidazo[4,5-*c*]pyridin-5-yl)ethyl)-1,4-diazepan-1-yl)ethan-1-one (14).** Compound **14** was synthesized following the general procedure, using 1-phenyl-1H,4H,5H,6H,7H-imidazo[4,5-c]pyridine (Enamine, Catalog #: EN300-116793) as the amine component. $^1$H NMR (400 MHz, $CD_3OD$) δ 9.75 (s, 1H), 9.07 (d, *J* = 27.7 Hz, 1H), 8.81 (d, *J* = 6.8 Hz, 1H), 8.62 (d, *J* = 6.7 Hz, 1H), 8.47 (d, *J* = 8.3 Hz, 1H), 8.24 (d, *J* = 7.1 Hz, 1H), 8.00 (t, *J* = 7.8 Hz, 1H), 7.66 (q, *J* = 3.0 Hz, 3H), 7.62 – 7.53 (m, 2H), 4.52 (d, *J* = 13.5 Hz, 2H), 4.43 (s, 2H), 4.34 (s, 2H), 4.04 – 3.89 (m, 2H), 3.89 – 3.43 (m, 14H), 3.23 (s, 2H), 3.09 – 3.01 (m, 2H), 2.83 (d, *J* = 39.7 Hz, 2H), 2.43 – 2.07 (m, 4H). HRMS-ESI (m/z): calcd for $C_{36}H_{44}N_8O_2$ $[M + H]^+$: 621.3660; found: 621.3657.

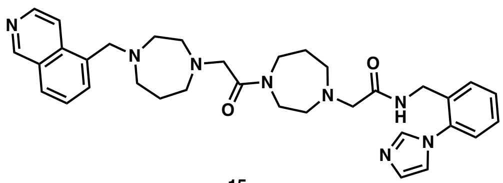

15

***N*-(2-(1*H*-Imidazol-1-yl)benzyl)-2-(4-(2-(4-(isoquinolin-5-ylmethyl)-1,4-diazepan-1-yl)acetyl)-1,4-diazepan-1-yl)acetamide (15).** Compound **15** was synthesized following the general procedure, using [2-(1H-imidazol-1-yl)phenyl]methanamine (Enamine, Catalog #: EN300-53224) as the amine component. $^1$H NMR (600 MHz, $CD_3OD$) δ 9.71 (s, 1H), 9.24 (s, 1H), 8.76 (d, *J* = 6.6 Hz, 1H), 8.61 (d, *J* = 6.6 Hz, 1H), 8.43 (d, *J* = 8.3 Hz, 1H), 8.20 (d, *J* = 7.1 Hz, 1H), 7.97 (dd, *J* = 8.3, 7.1 Hz, 1H), 7.88 (s, 1H), 7.78 (s, 1H), 7.70 – 7.61 (m, 2H), 7.56 (d, *J* = 1.9 Hz, 1H), 7.52 (dd, *J* = 7.9, 1.3 Hz, 1H), 4.39 (s, 2H), 4.29 (s, 2H), 4.02 (d, *J* = 6.1 Hz, 2H), 3.94 – 3.68 (m, 3H), 3.58 – 3.36 (m, 11H), 3.19 – 3.13 (m, 2H), 3.01 (t, *J* = 5.8 Hz, 2H), 2.33 – 2.06 (m, 4H). HRMS-ESI (m/z): calcd for $C_{34}H_{42}N_8O_2$ $[M + H]^+$: 595.3504; found: 595.3502.

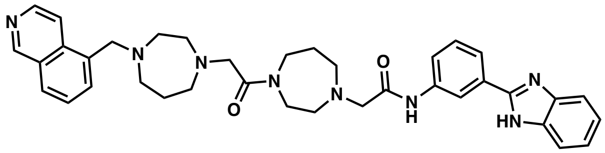

16

***N*-(3-(1*H*-Benzo[*d*]imidazol-2-yl)phenyl)-2-(4-(2-(4-(isoquinolin-5-ylmethyl)-1,4-diazepan-1-yl)acetyl)-1,4-diazepan-1-yl)acetamide (16).** Compound **16** was synthesized following the general procedure, using 3-(1H-1,3-benzodiazol-2-yl)aniline (Enamine, Catalog #: EN300-11470) as the amine component. $^1$H NMR (600 MHz, $CD_3OD$) δ 9.62 (s, 1H), 8.67 (d, *J* = 6.4 Hz, 1H), 8.59 (d, *J* = 6.2 Hz, 1H), 8.58 – 8.54 (m, 1H), 8.36 (d, *J* = 8.5 Hz, 1H), 8.13 (d, *J* = 7.3 Hz, 1H), 7.94 – 7.89 (m, 1H), 7.88 (dt, *J* = 7.8, 1.3 Hz, 1H), 7.85 – 7.77 (m, 3H), 7.70 (dd, *J* = 8.8, 7.2 Hz, 1H), 7.59 (dd, *J* = 6.1, 3.0 Hz, 2H), 4.36 (s, 2H), 4.28 (s, 2H), 4.23 (d, *J* = 20.7 Hz,

2H), 4.02 – 3.72 (m, 2H), 3.68 – 3.37 (m, 10H), 3.13 (s, 2H), 3.00 (t, *J* = 5.8 Hz, 2H), 2.22 (d, *J* = 114.6 Hz, 4H). HRMS-ESI (m/z): calcd for $C_{37}H_{42}N_8O_2$ $[M + H]^+$: 631.3504; found: 631.3501.

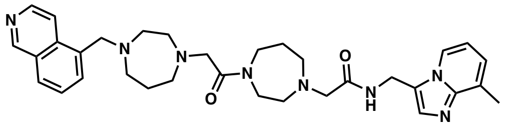

17

**2-(4-(2-(4-(Isoquinolin-5-ylmethyl)-1,4-diazepan-1-yl)acetyl)-1,4-diazepan-1-yl)-*N*-((8-methylimidazo[1,2-*a*]pyridin-3-yl)methyl)acetamide (17).** Compound **17** was synthesized following the general procedure, using 1-(8-Methylimidazo[1,2-a]pyridin-3-yl)methanamine (Sigma Aldrich, Catalog #: CBR01374) as the amine component. $^1$H NMR (600 MHz, $CD_3OD$) δ 9.67 (s, 1H), 8.71 (d, *J* = 6.4 Hz, 1H), 8.67 (d, *J* = 6.8 Hz, 1H), 8.60 (d, *J* = 6.5 Hz, 1H), 8.40 (d, *J* = 8.2 Hz, 1H), 8.16 (d, *J* = 7.1 Hz, 1H), 8.07 (s, 1H), 7.94 (dd, *J* = 8.3, 7.1 Hz, 1H), 7.82 (d, *J* = 7.1 Hz, 1H), 7.46 (t, *J* = 6.7 Hz, 1H), 4.37 (s, 2H), 4.28 (s, 2H), 4.04 (s, 2H), 3.96 – 3.63 (m, 3H), 3.61 – 3.32 (m, 11H), 3.17 – 3.11 (m, 2H), 3.00 (t, *J* = 5.7 Hz, 2H), 2.66 (s, 3H), 2.31 – 2.07 (m, 4H). HRMS-ESI (m/z): calcd for $C_{33}H_{42}N_8O_2$ $[M + H]^+$: 583.3504; found: 583.3500.

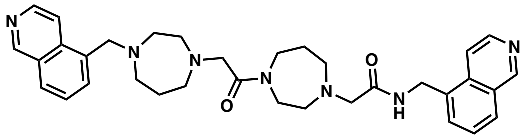

18

***N*-(Isoquinolin-5-ylmethyl)-2-(4-(2-(4-(isoquinolin-5-ylmethyl)-1,4-diazepan-1-yl)acetyl)-1,4-diazepan-1-yl)acetamide (18).** Compound **17** was synthesized following the general procedure, using 5-Isoquinolinemethanamine (Combi-Block, Catalog #: QB-8131) as the amine component. $^1$H NMR (600 MHz, $CD_3OD$) δ 9.67 (s, 2H), 8.72 (d, *J* = 6.5 Hz, 1H), 8.63 – 8.57 (m, 2H), 8.51 (d, *J* = 6.5 Hz, 1H), 8.40 (d, *J* = 8.3 Hz, 1H), 8.37 (d, *J* = 8.3 Hz, 1H), 8.16 (d, *J* = 7.1 Hz, 1H), 8.12 (d, *J* = 7.1 Hz, 1H), 7.94 (td, *J* = 7.2, 5.2 Hz, 2H), 5.00 (s, 2H), 4.37 (s, 2H), 4.28 (s, 2H), 4.07 (s,

2H), 3.93 – 3.38 (m, 12H), 3.15 (d, $J$ = 4.7 Hz, 2H), 3.00 (t, $J$ = 5.8 Hz, 2H), 2.32 – 2.05 (m, 4H). $^{13}$C NMR (150 MHz, $CD_3OD$) δ 166.56, 165.82, 150.20, 150.04, 139.00, 138.38, 137.82, 136.45, 135.71, 135.52, 135.27, 131.55, 131.18, 131.03, 129.97, 129.76, 122.97, 121.88, 59.81, 58.62, 58.60, 56.36, 56.08, 55.90, 55.53, 55.06, 50.61, 45.99, 41.60, 41.07, 25.42, 24.91. HRMS-ESI (m/z): calcd for $C_{34}H_{41}N_7O_2$ $[M + H]^+$: 580.3395; found: 580.3393.

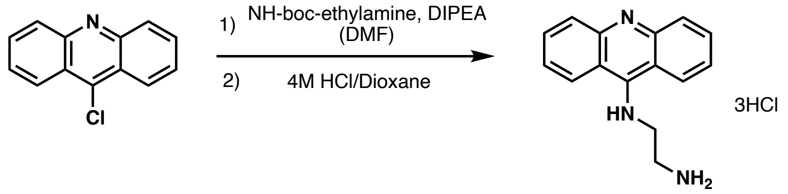


**19a**

***N*[1]-(Acridin-9-yl)ethane-1,2-diamine hydrochloride (19a).** To a solution of 9-chloroacridine (213 mg, 0.997 mmol, 1.3 eq) in DMF (4.0 mL) was added *tert*-butyl (2-aminoethyl)carbamate (206 mL, 1.30 mmol) and DIPEA (348 mL, 2.00 mmol, 2.0 eq) at room temperature, and the solution was heated at 70 °C for 2 d. The reaction mixture was cooled to room temperature, followed by concentration under reduced pressure. The residue was purified by automated silica gel flush column chromatography system (Isolera One) with EtOAc-MeOH (100:0 to 85:15) to obtain *tert*-butyl (2-(acridin-9-ylamino)ethyl)carbamate. The boc-protected amine was treated with 4 M HCl in dioxane (10 mL) and the solution was stirred for 16 h at room temperature. The precipitated salt was filtered off and washed with EtOAc to obtain *N*[1]-(acridin-9-yl)ethane-1,2-diamine hydrochloride (274 mg, 0.790 mmol, 79% in 2 steps) as a yellow solid and directly used for the synthesis of **19**.

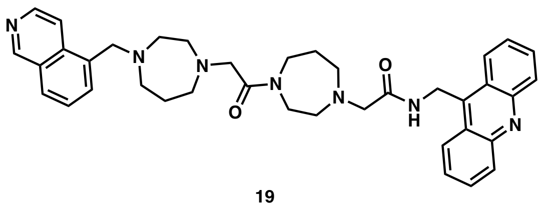

19

***N*-(Acridin-9-ylmethyl)-2-(4-(2-(4-(isoquinolin-5-ylmethyl)-1,4-diazepan-1-yl)acetyl)-1,4-diazepan-1-yl)acetamide (19).** Compound **19** was synthesized following the general procedure, using $N^1$-(acridin-9-yl)ethane-1,2-diamine hydrochloride (**19a**) as the amine component. $^1$H NMR (600 MHz, $CD_3OD$) δ 9.53 (s, 1H), 8.57 (d, $J$ = 6.5 Hz, 1H), 8.48 (dd, $J$ = 9.7, 7.2 Hz, 3H), 8.27 (d, $J$ = 8.3 Hz, 1H), 8.03 (d, $J$ = 7.1 Hz, 1H), 7.89 (ddd, $J$ = 8.3, 6.9, 1.2 Hz, 2H), 7.84 – 7.78 (m, 1H), 7.74 (dd, $J$ = 8.6, 1.2 Hz, 2H), 7.50 (ddd, $J$ = 8.6, 5.8, 1.3 Hz, 2H), 4.31 – 4.21 (m, 4H), 4.15 (d, $J$ = 5.6 Hz, 2H), 3.90 – 3.69 (m, 6H), 3.68 – 3.56 (m, 2H), 3.46 – 3.23 (m, 10H), 3.05 – 2.97 (m, 2H), 2.89 (t, $J$ = 5.8 Hz, 2H), 2.01 (t, $J$ = 42.9 Hz, 4H). HRMS-ESI (m/z): calcd for $C_{39}H_{46}N_8O_2$ $[M + H]^+$: 659.3817; found: 659.3814.

### COMPOUND CHARACTERIZATION

**1a** ($^1$H-NMR, 400 MHz, $CD_3OD$):

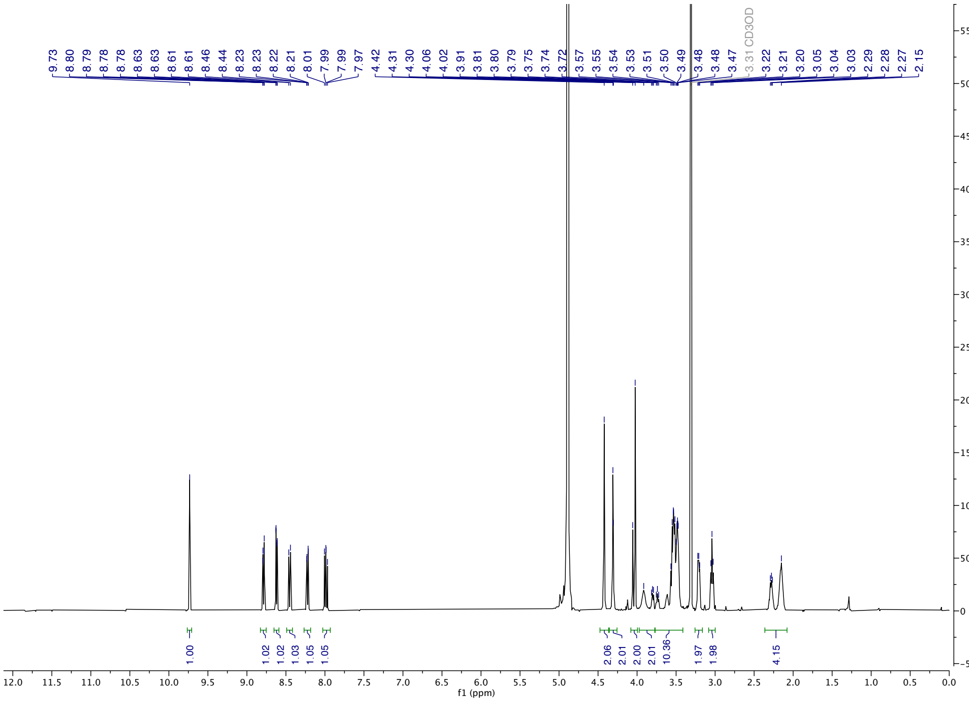

**1a** ($^{13}$C-NMR, 150 MHz, $CD_3OD$):

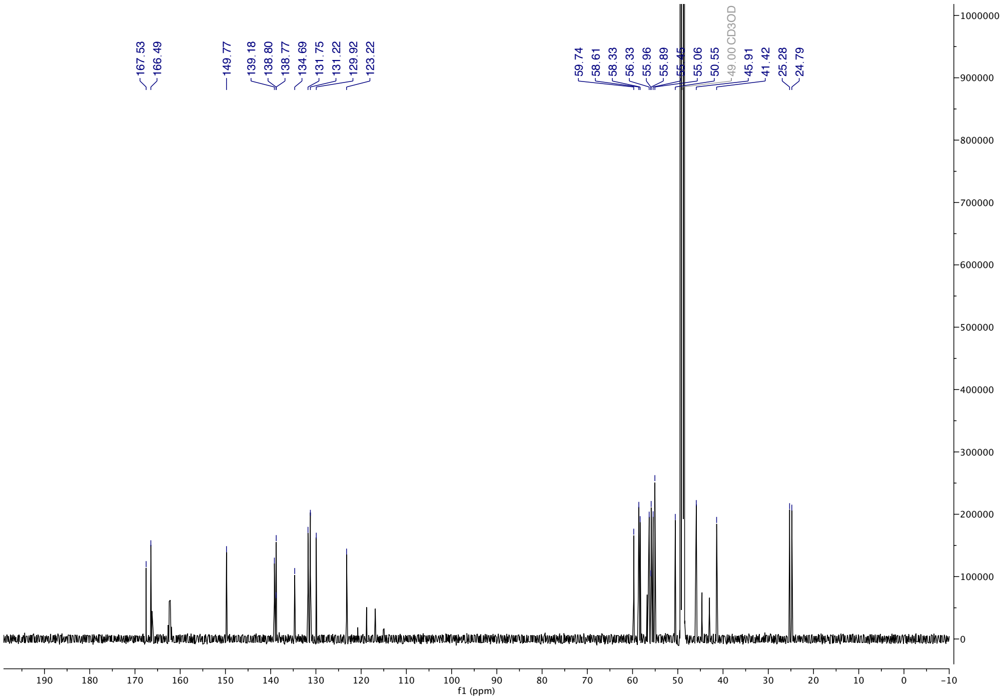

**1a** (HPLC, λ = 254 nm, 0-60% MeOH in $H_2O$, +0.1% TFA, 60 min; 60-100% MeOH in $H_2O$, +0.1% TFA, 5 min;):

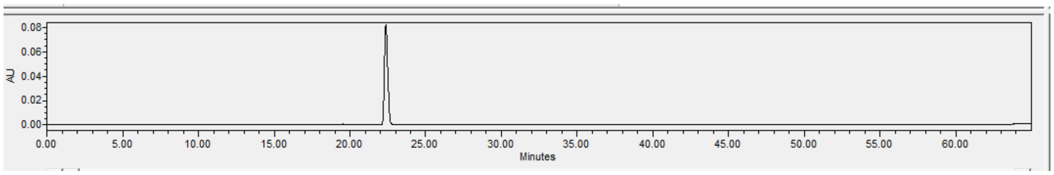


**1a** (ESI-HRMS):

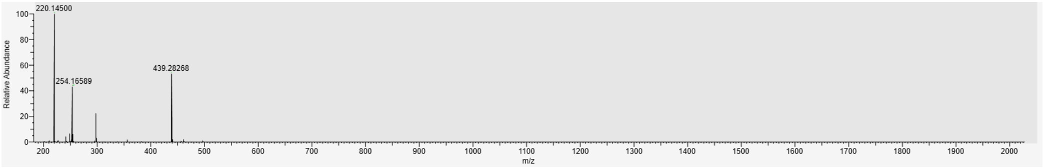

**1-COOH** ($^1$H-NMR, 600 MHz, $CD_3OD$):

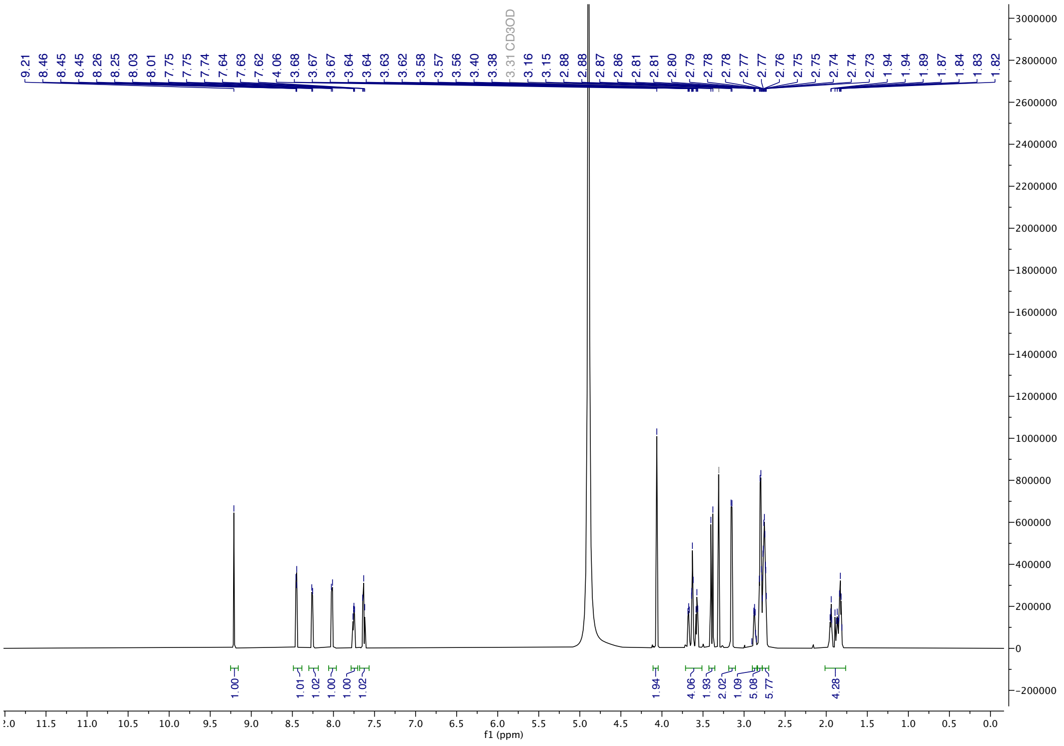

**1-COOH** ($^{13}$C-NMR, 150 MHz, $CD_3OD$):

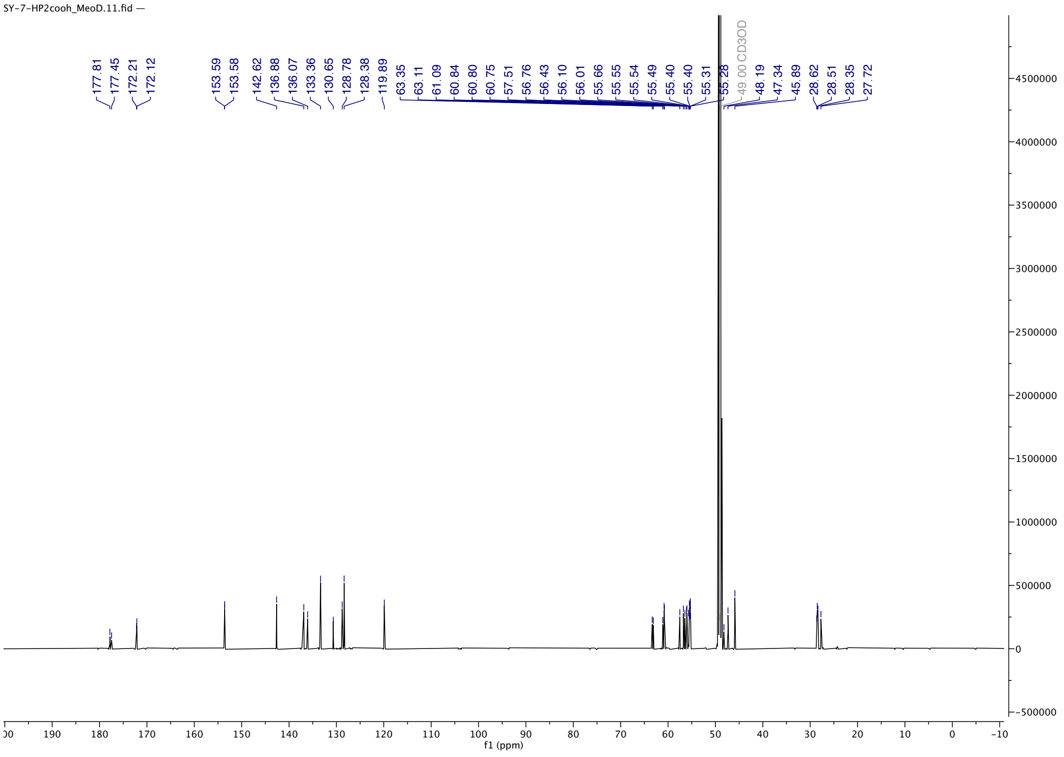

**1-COOH** (HPLC, λ = 254 nm, 0-60% MeOH in $H_2O$, +0.1% TFA, 60 min; 60-100% MeOH in $H_2O$, +0.1% TFA, 5 min):

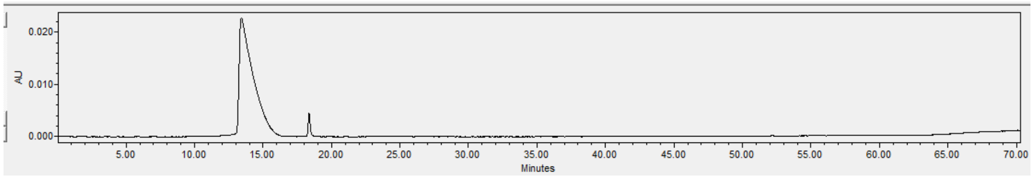


**1-COOH** (ESI-HRMS):

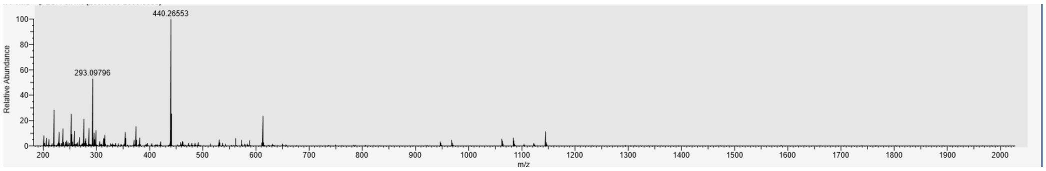

**1** ($^{1}$H-NMR, 600 MHz, $CD_3OD$):

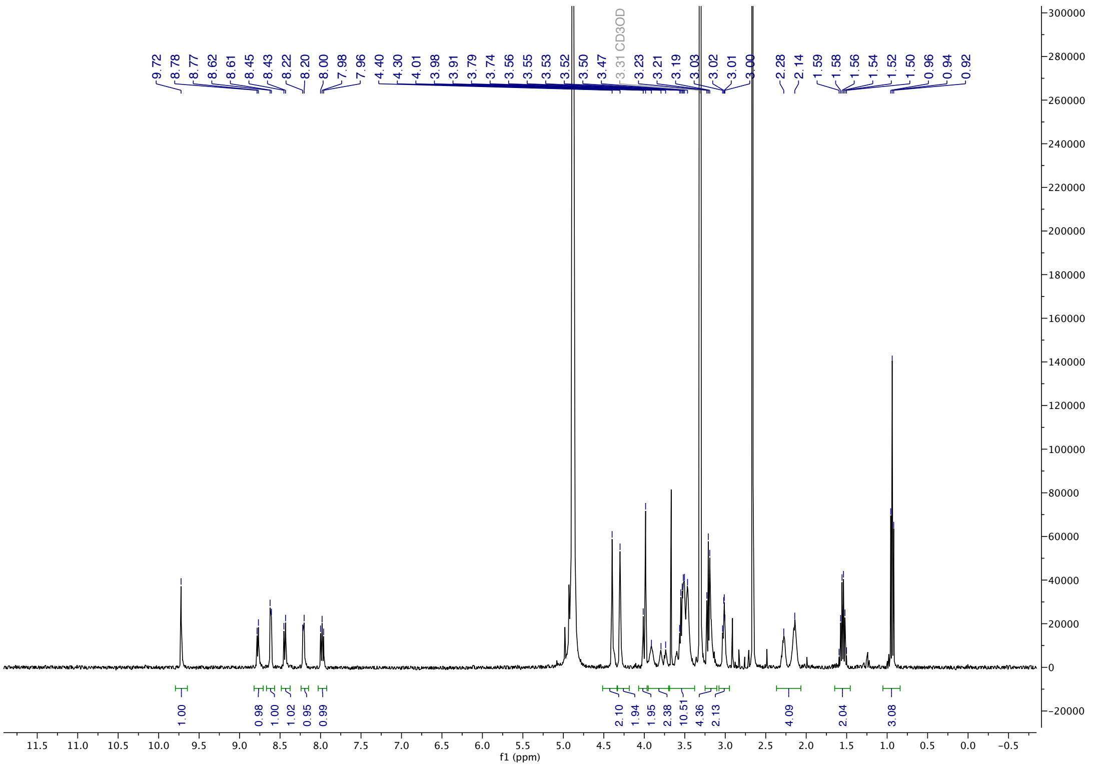


**1** ($^{13}$C-NMR, 150 MHz, $CD_3OD$):

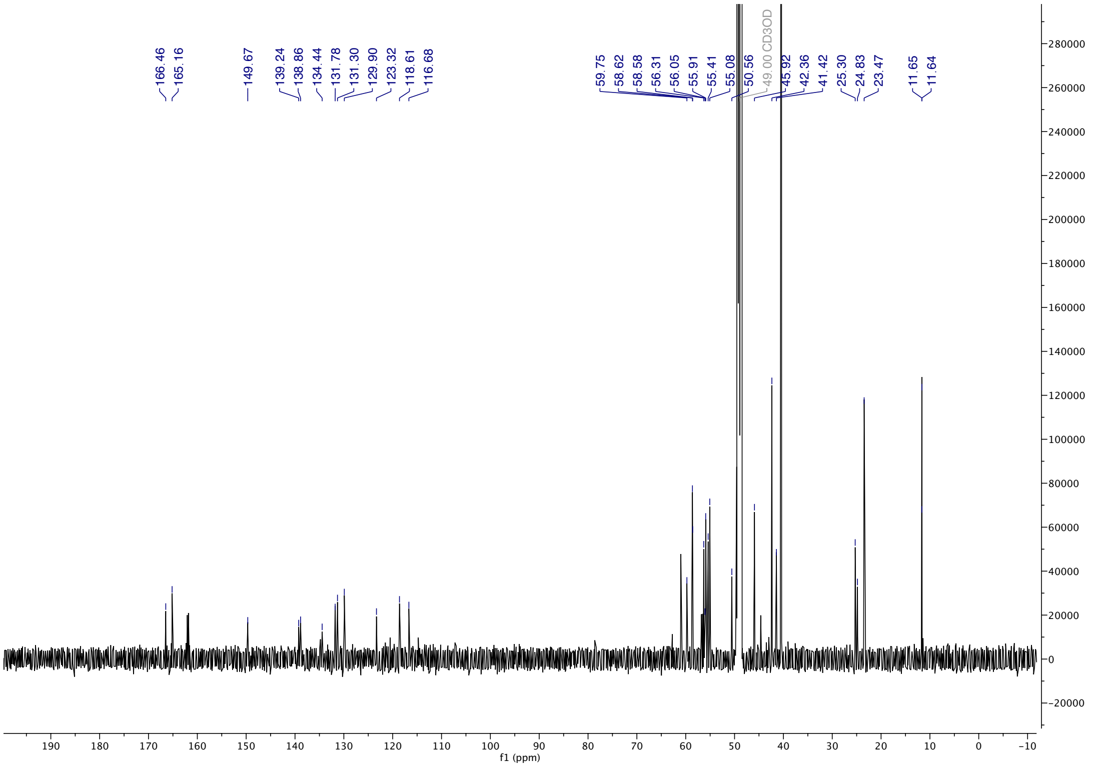

**1** (HPLC, λ = 254 nm, 0-60% MeOH in $H_2O$, +0.1% TFA, 60 min; 60-100% MeOH in $H_2O$, +0.1% TFA, 5 min):

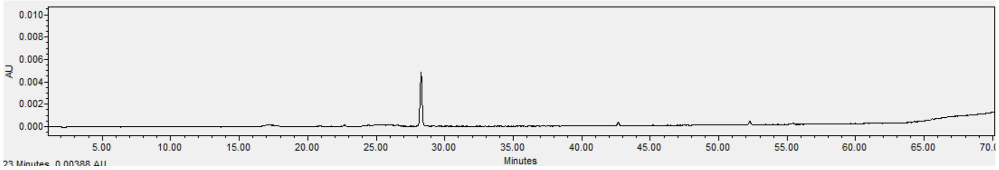


**1** (ESI-HRMS):

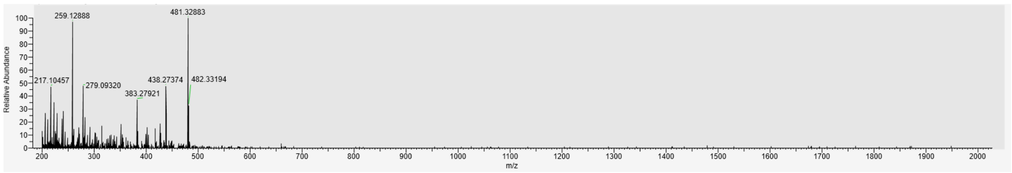

**3** ($^1$H-NMR, 600 MHz, $CD_3OD$):

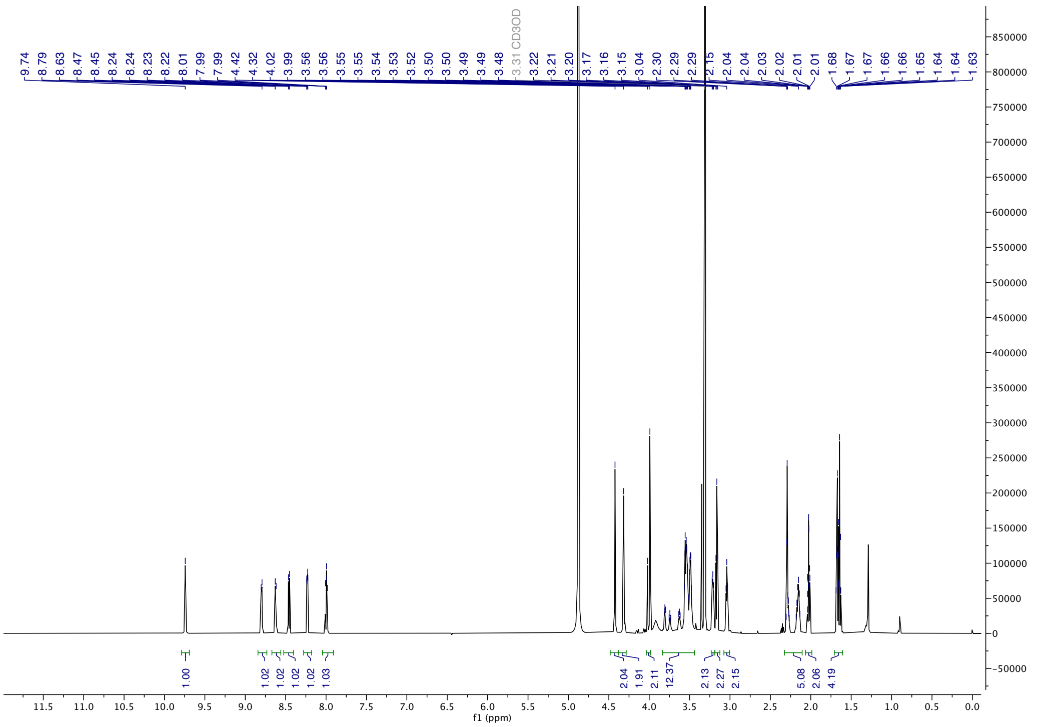

**3** ($^{13}$C-NMR, 150 MHz, $CD_3OD$):

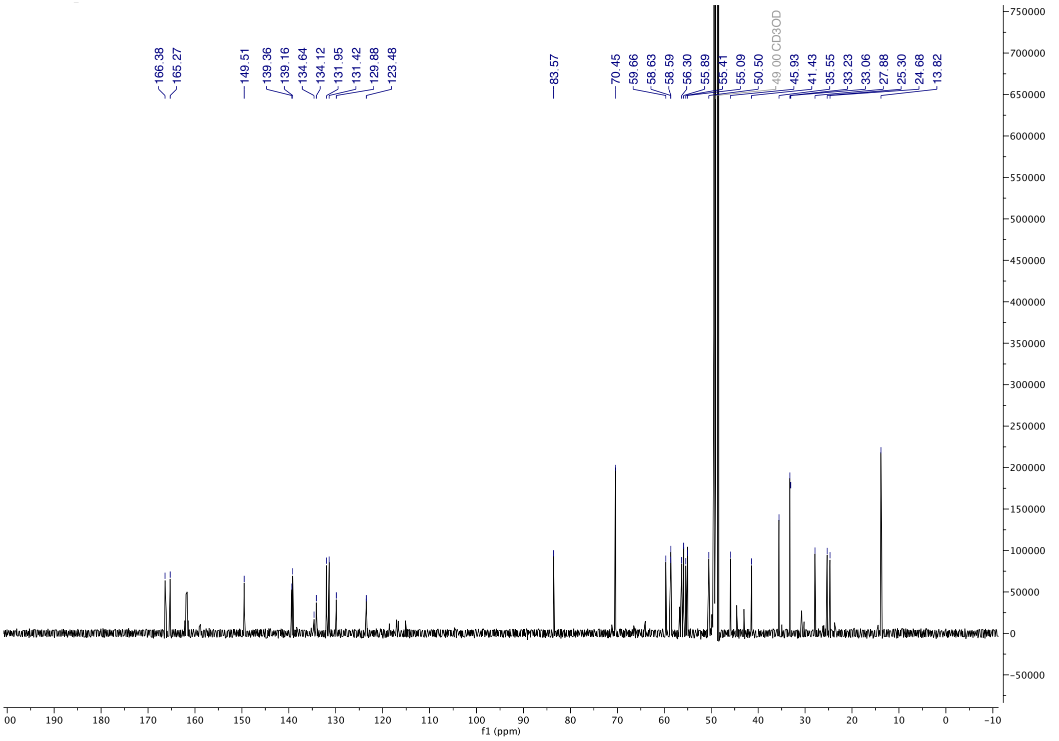

**3** (HPLC, λ = 254 nm, 0-60% MeOH in $H_2O$, +0.1% TFA, 60 min; 60-100% MeOH in $H_2O$, +0.1% TFA, 5 min):

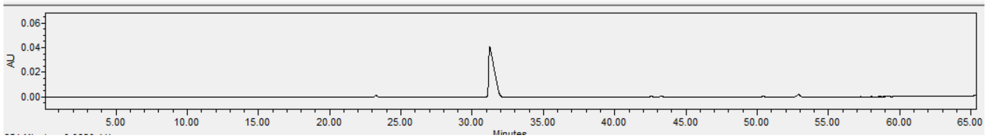


**3** (ESI-HRMS):

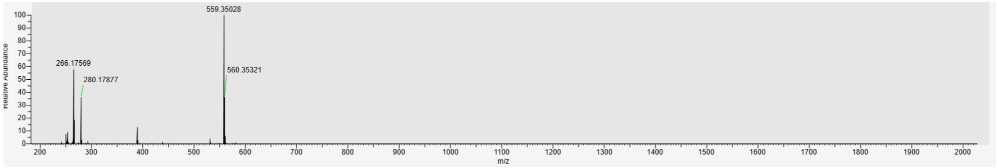

**4** ([1]H-NMR, 600 MHz, $CD_3OD$):

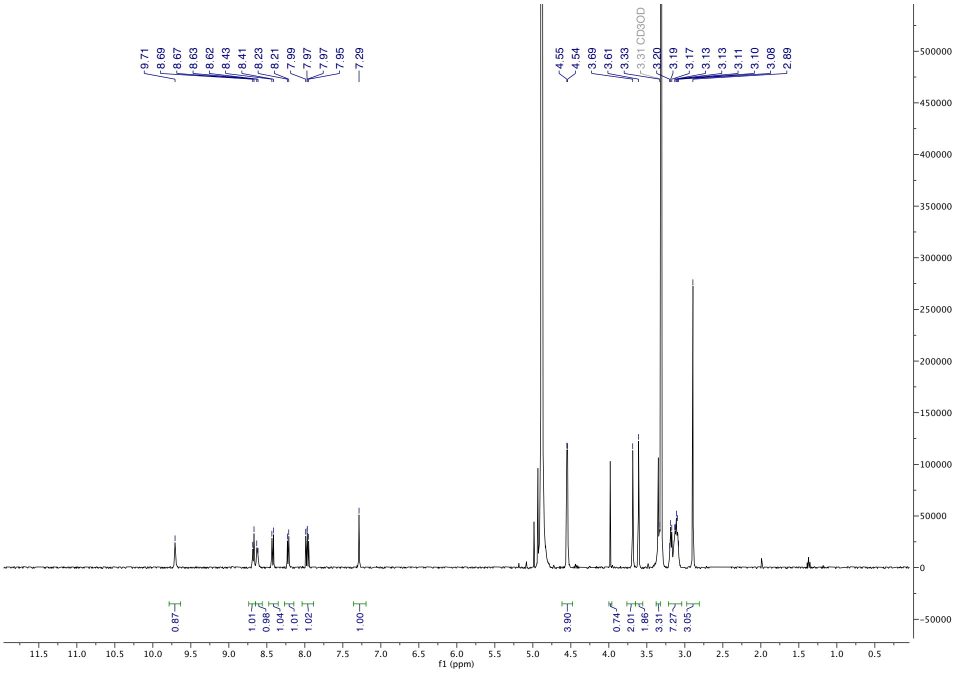

**5** ([1]H-NMR, 600 MHz, $CD_3OD$):

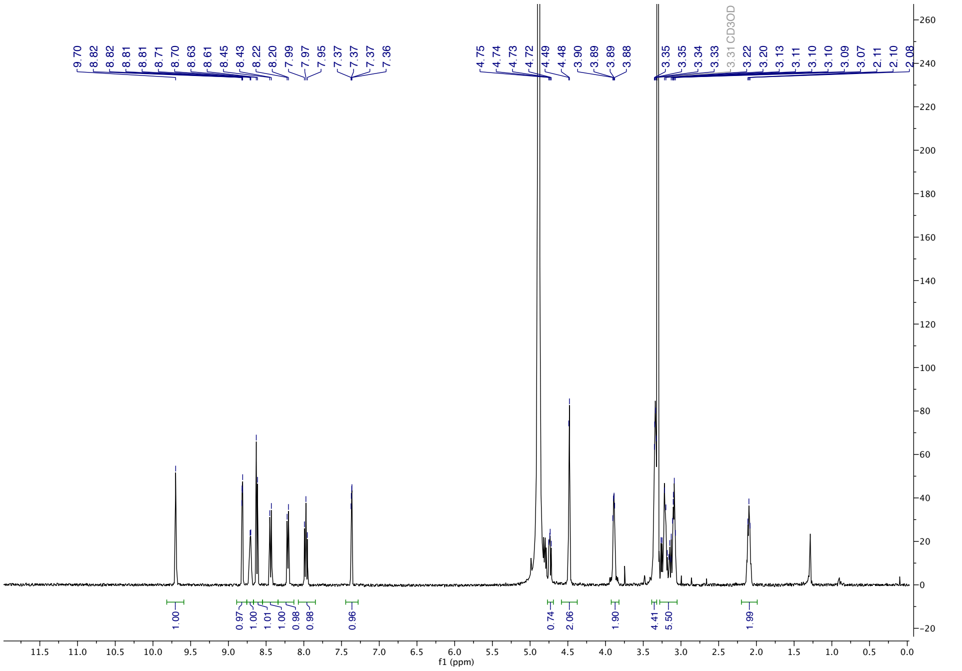

**6** ([1]H-NMR, 600 MHz, $CD_3OD$):

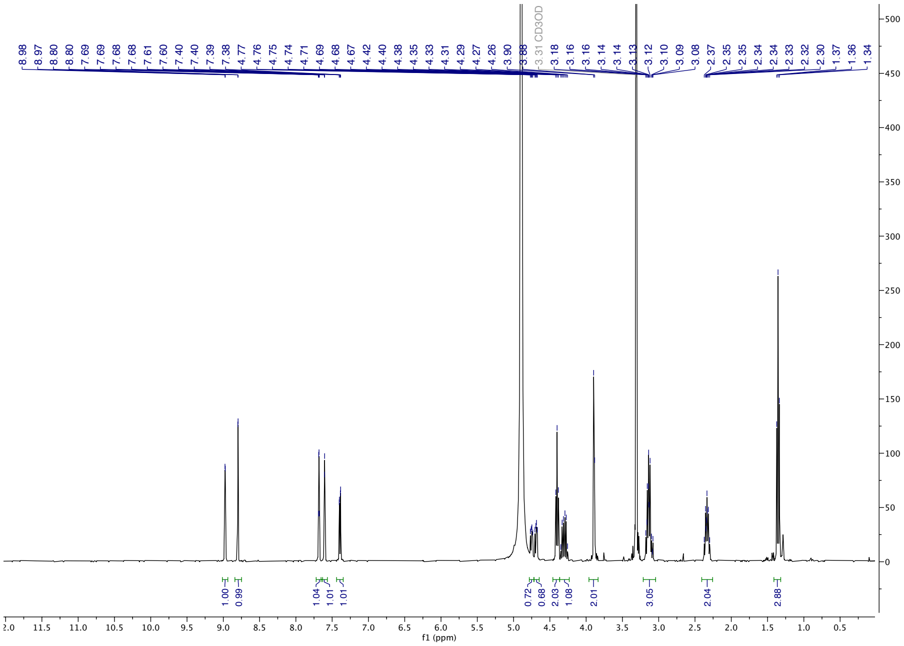

**7** ([1]H-NMR, 600 MHz, $CD_3OD$):

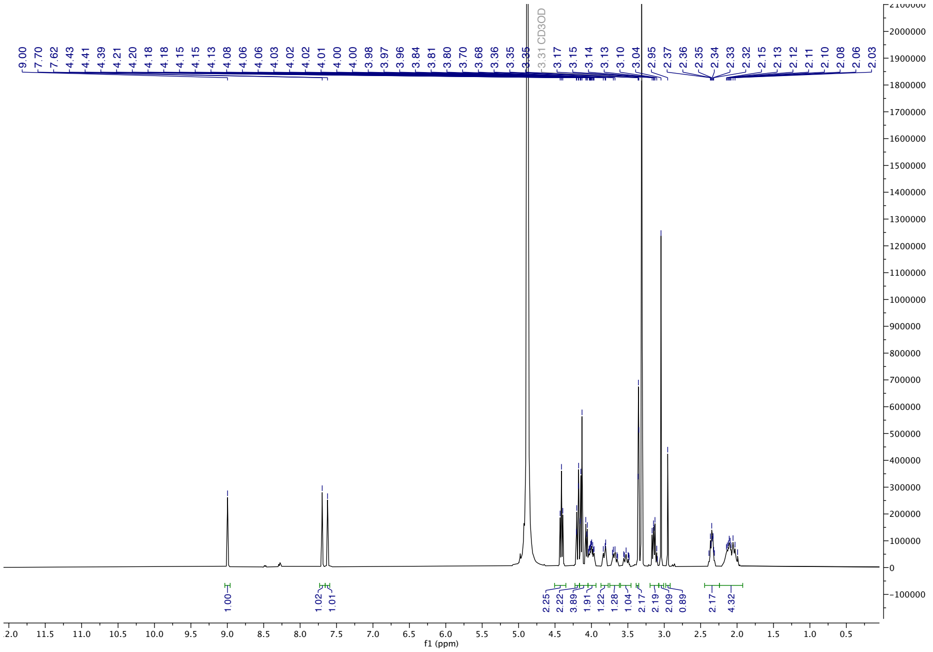

**8** ($^1$H-NMR, 600 MHz, $CD_3OD$):

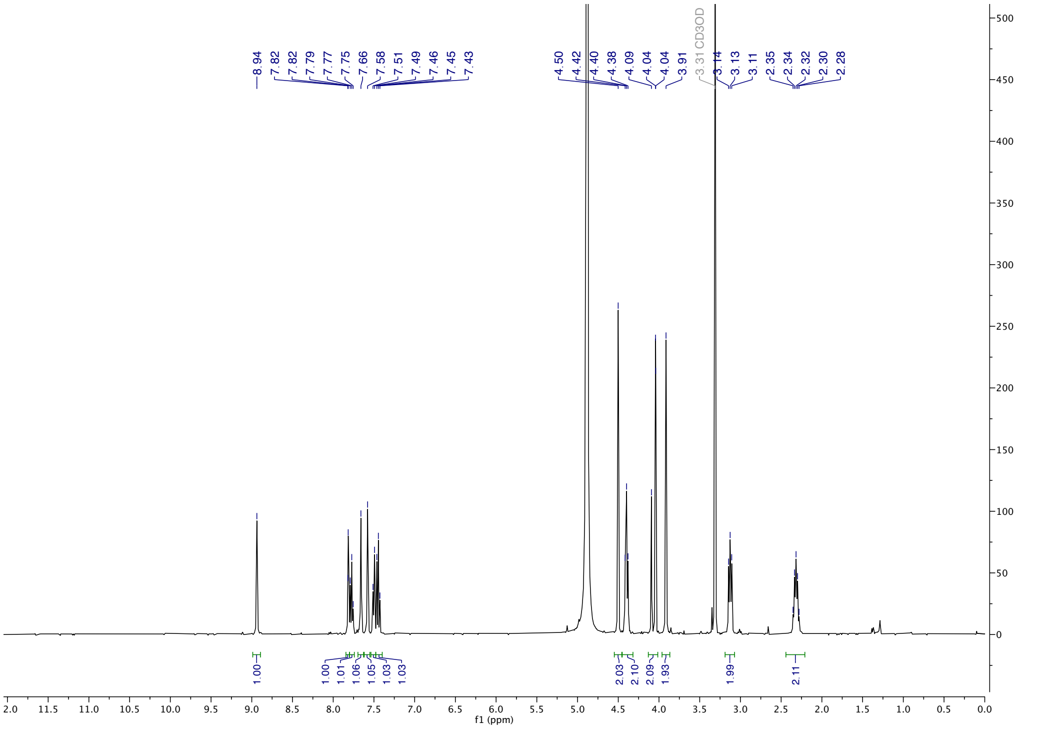

**9** ($^1$H-NMR, 600 MHz, $CD_3OD$):

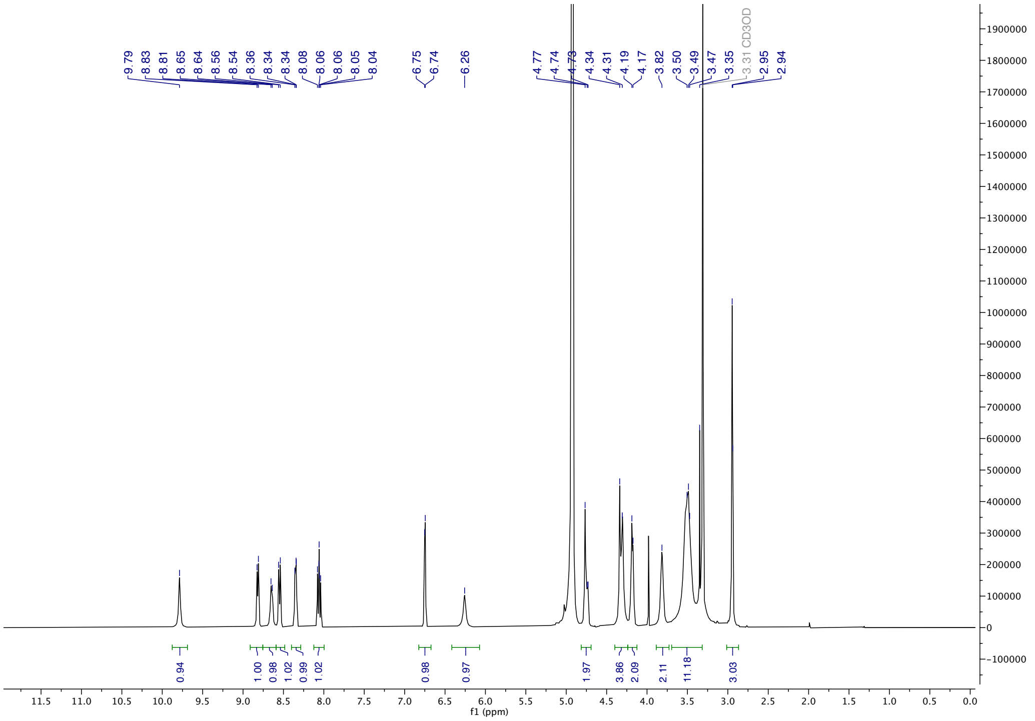

**11** ($^{1}$H-NMR, 600 MHz, $CD_3OD$):

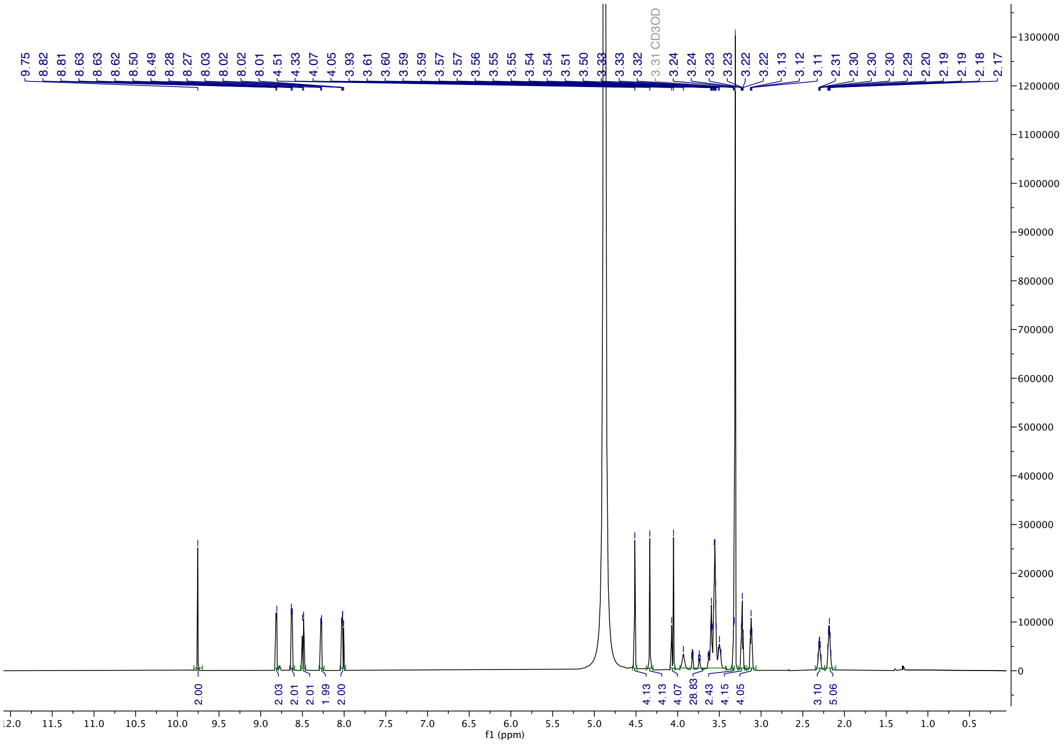


**11** ($^{13}$C-NMR, 150 MHz, $CD_3OD$):

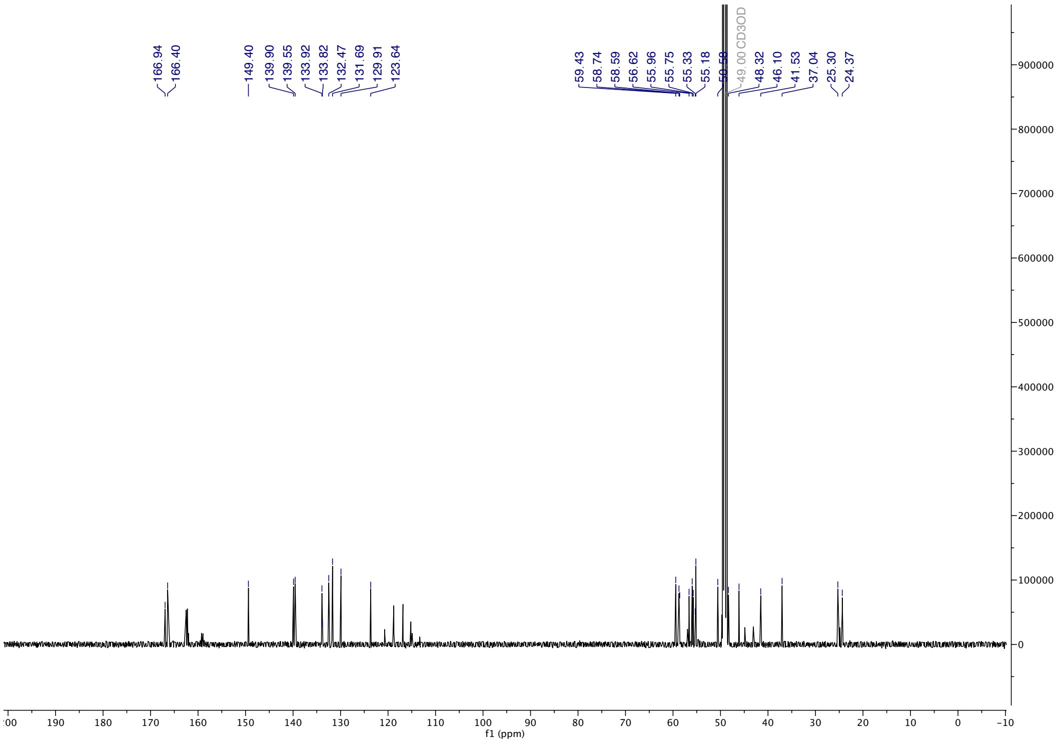

**11** (HPLC, λ = 254 nm, 0-60% MeOH in $H_2O$, +0.1% TFA, 60 min; 60-100% MeOH in $H_2O$, +0.1% TFA, 5 min):

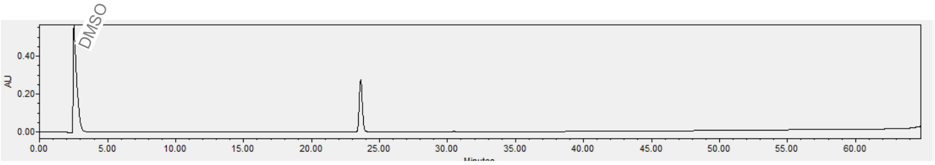


**11** (ESI-HRMS):

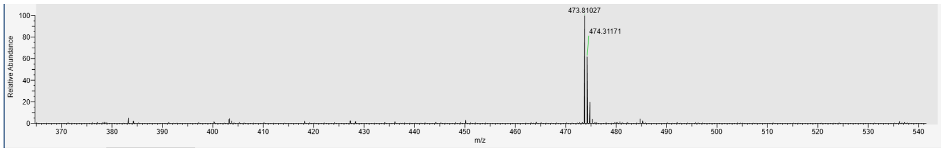

**12** ($^1$H-NMR, 600 MHz, $CD_3OD$):

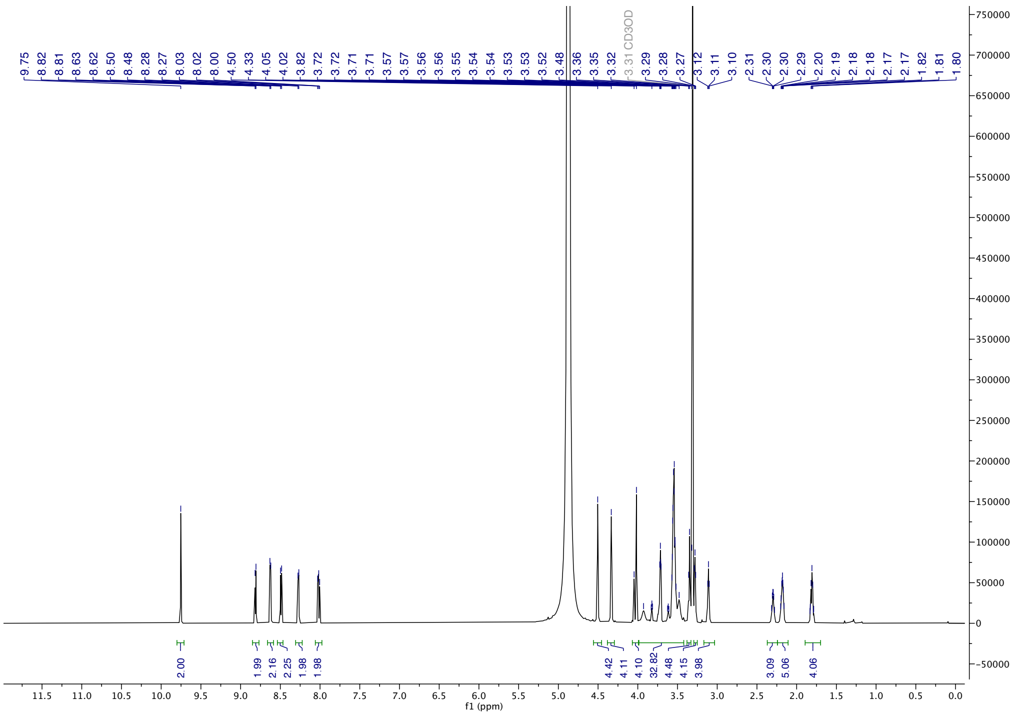


**12** ($^{13}$C-NMR, 150 MHz, $CD_3OD$):

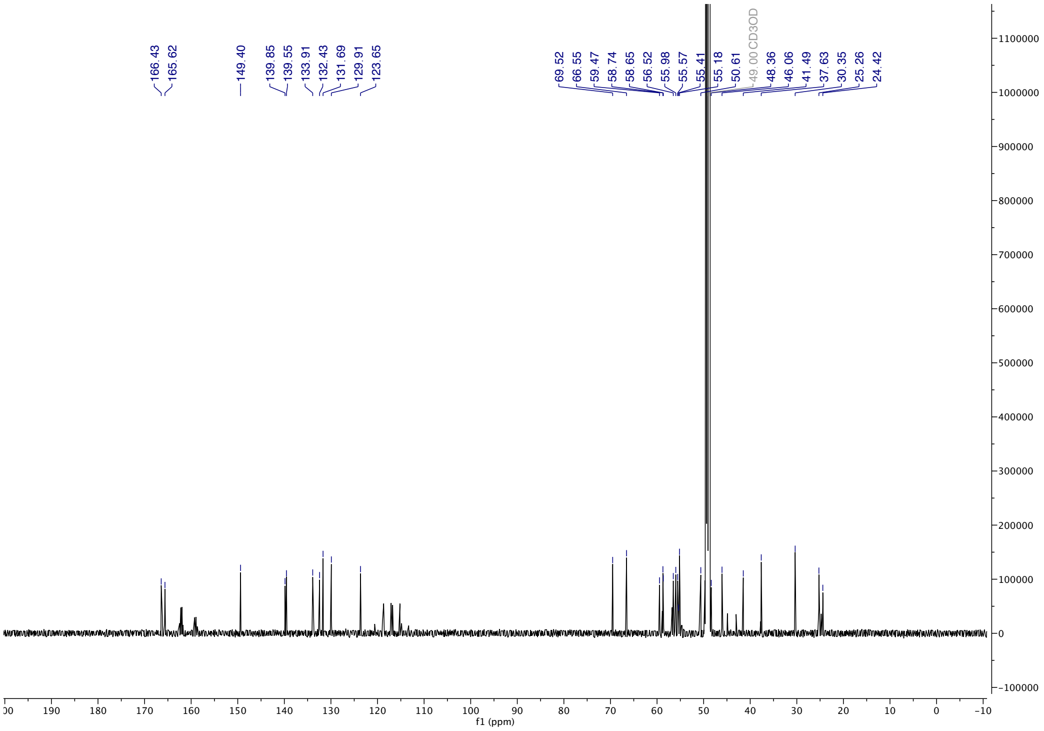

**12** (HPLC, λ = 254 nm, 0-60% MeOH in $H_2O$, +0.1% TFA, 60 min; 60-100% MeOH in $H_2O$, +0.1% TFA, 5 min):

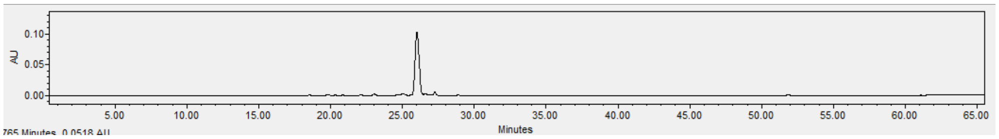


**12** (ESI-HRMS):

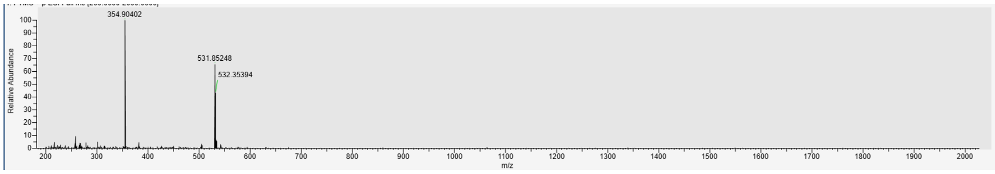

**13** ($^1$H-NMR, 600 MHz, $CD_3OD$):

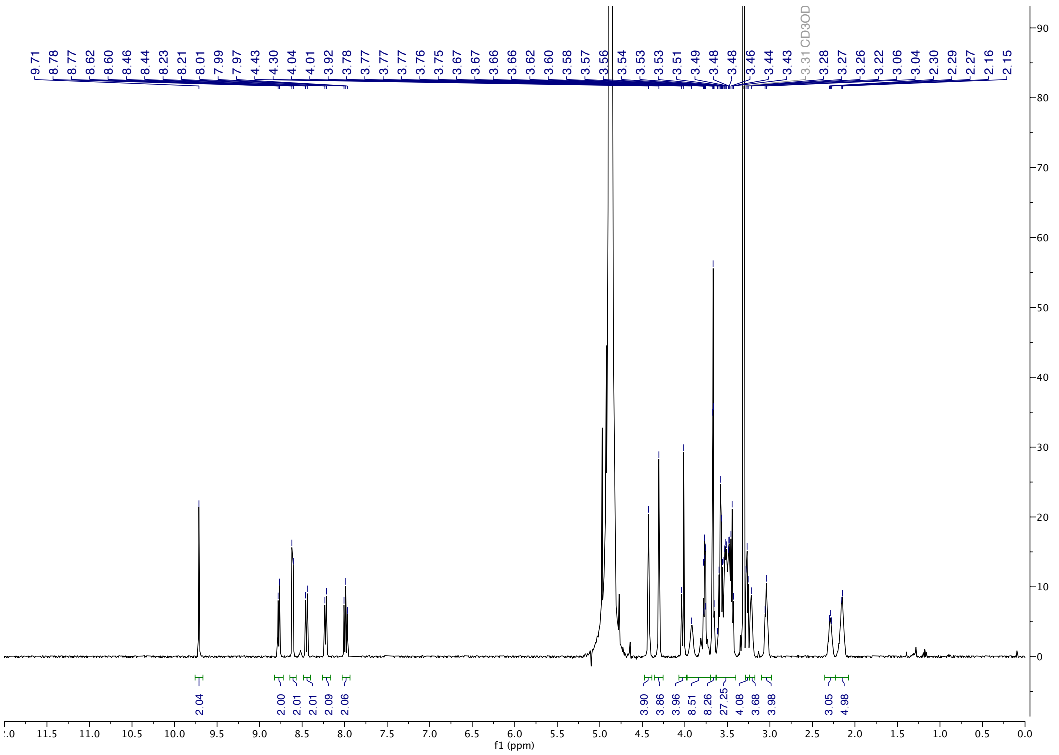

**13** ($^{13}$C-NMR, 150 MHz, $CD_3OD$):

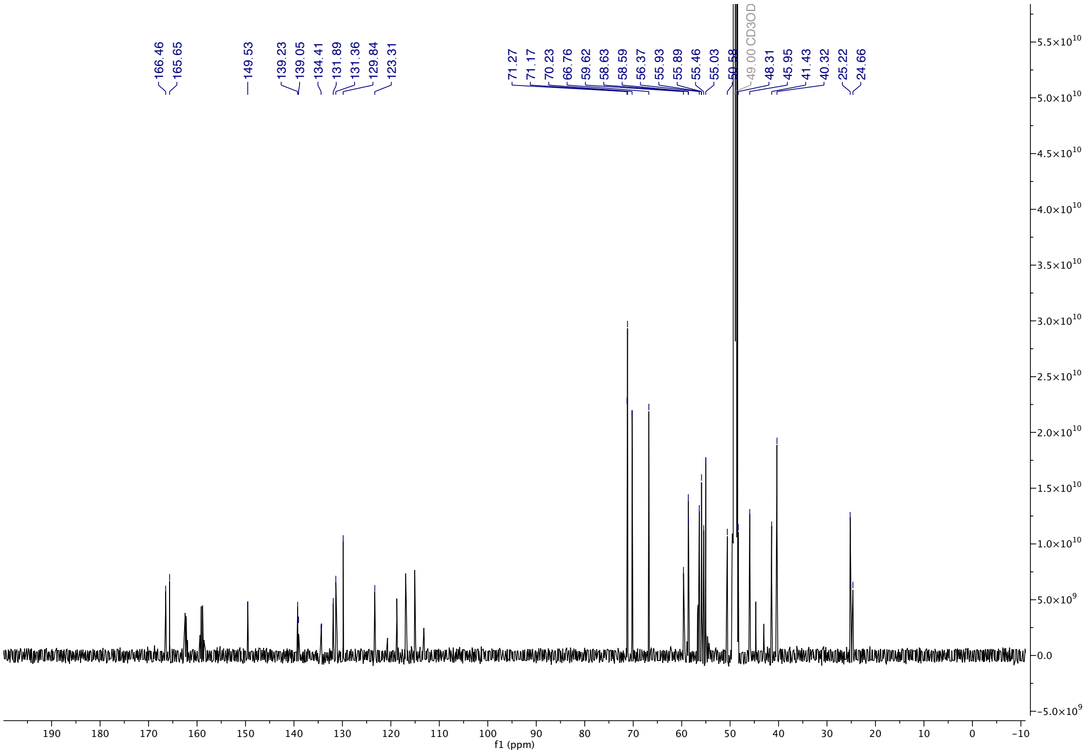

**13** (HPLC, λ = 254 nm, 0-60% MeOH in $H_2O$, +0.1% TFA, 60 min; 60-100% MeOH in $H_2O$, +0.1% TFA, 5 min):

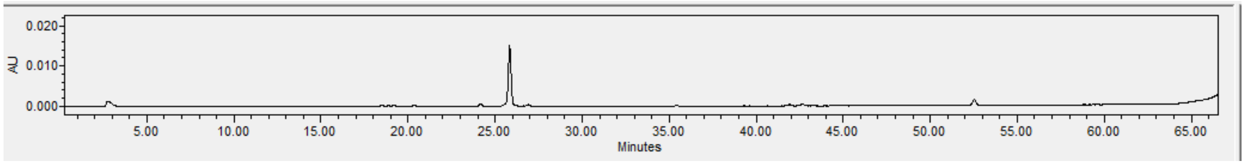


**13** (ESI-HRMS):

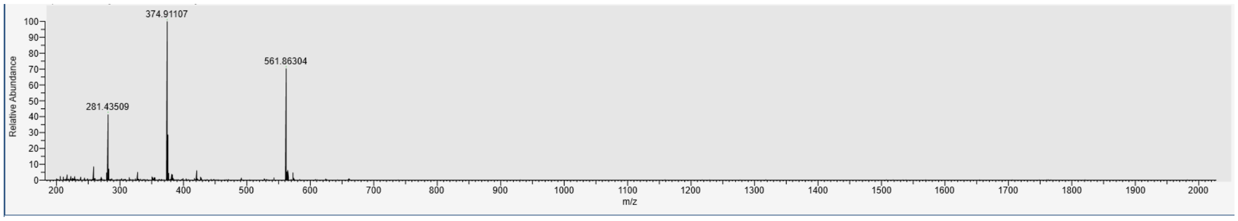

**14** ($^1$H-NMR, 600 MHz, $CD_3OD$):

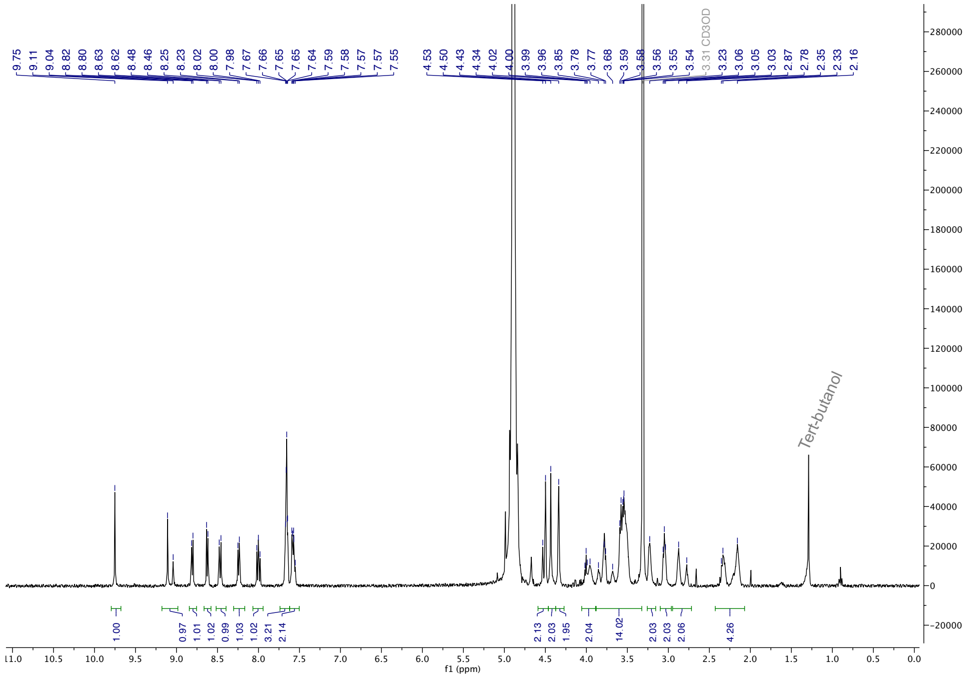


**14** (HPLC, λ = 254 nm, 0-60% MeOH in $H_2O$, +0.1% TFA, 60 min; 60-100% MeOH in $H_2O$, +0.1% TFA, 5 min):

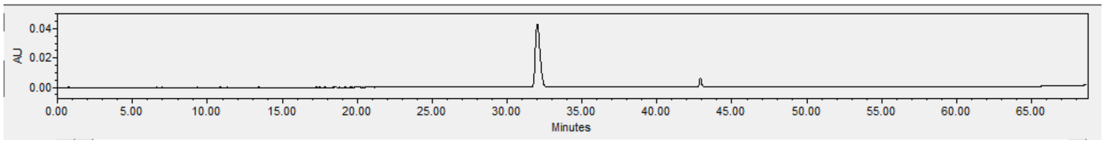


**14** (ESI-HRMS):

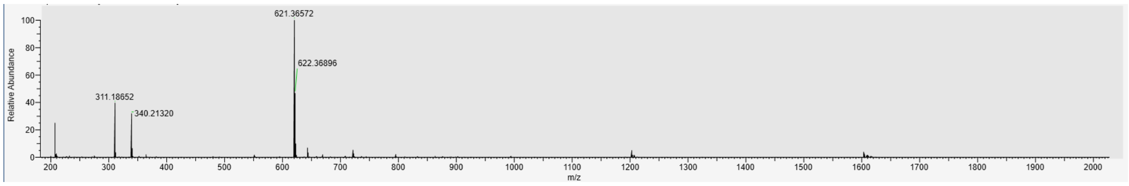

**15** ($^1$H-NMR, 600 MHz, $CD_3OD$):

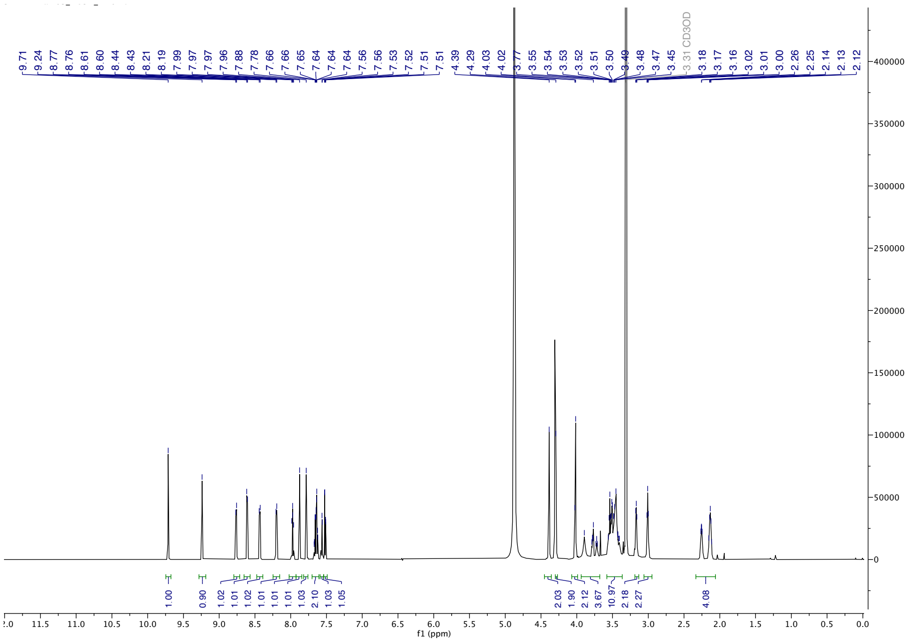

**15** (HPLC, λ = 254 nm, 0-60% MeOH in $H_2O$, +0.1% TFA, 60 min; 60-100% MeOH in $H_2O$, +0.1% TFA, 5 min):

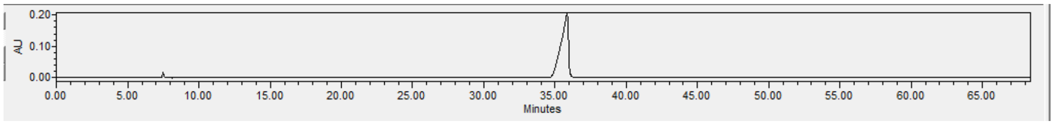


**15** (ESI-HRMS):

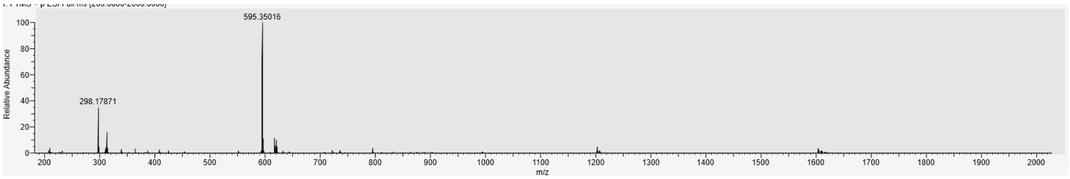

**16** ($^{1}H$-NMR, 600 MHz, $CD_3OD$):

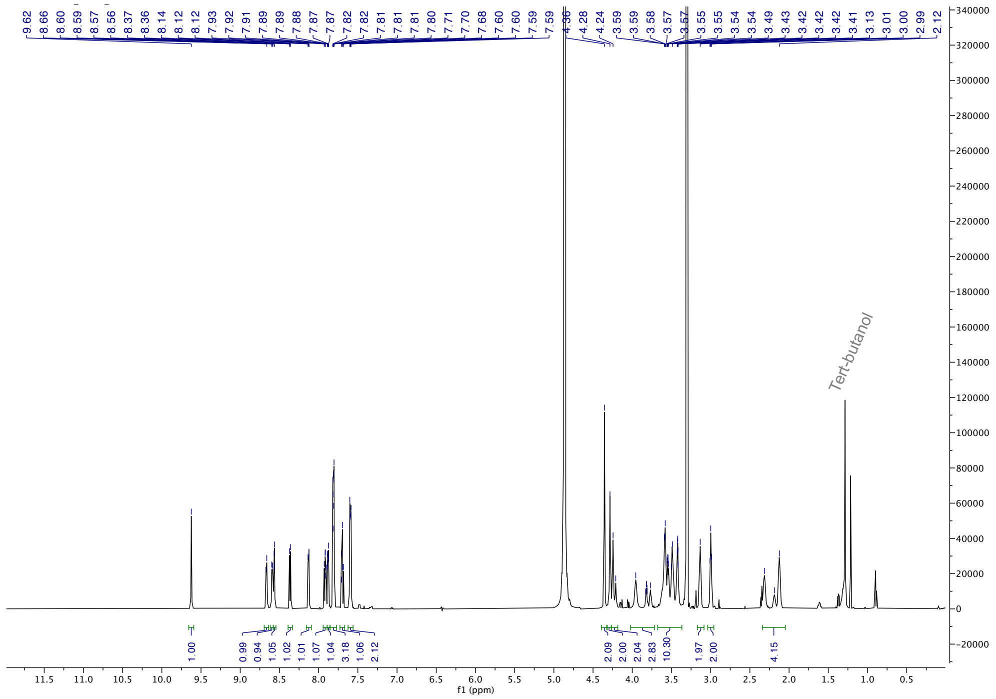

**16** (HPLC, λ = 254 nm, 0-60% MeOH in $H_2O$, +0.1% TFA, 60 min; 60-100% MeOH in $H_2O$, +0.1% TFA, 5 min):

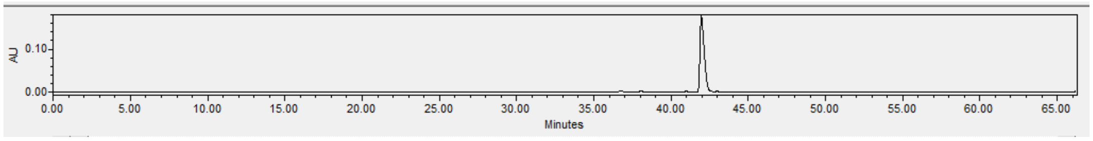

**16** (ESI-HRMS):

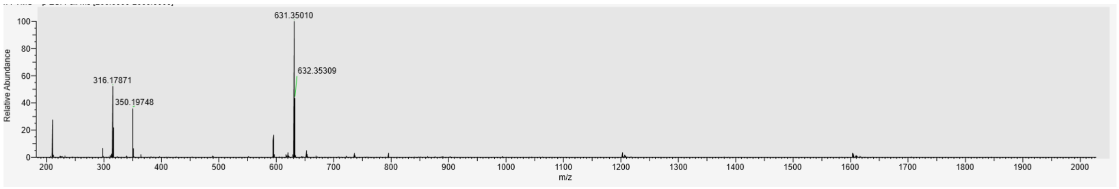

**17** ($^{1}H$-NMR, 600 MHz, $CD_3OD$):

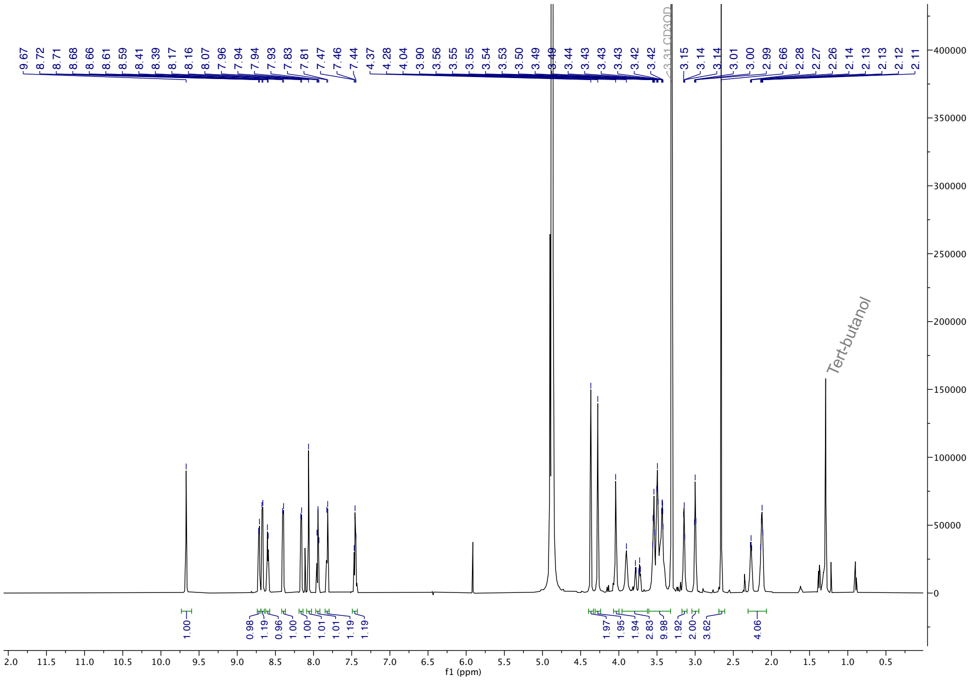

**17** (HPLC, λ = 254 nm, 0-60% MeOH in $H_2O$, +0.1% TFA, 60 min; 60-100% MeOH in $H_2O$, +0.1% TFA, 5 min):

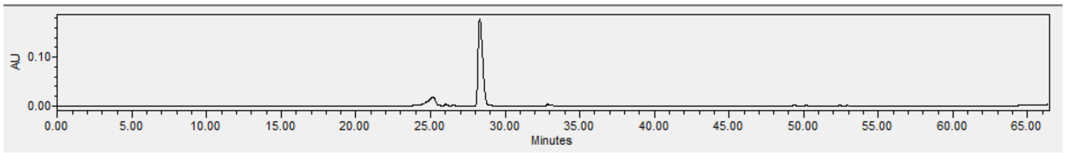

**17** (ESI-HRMS):

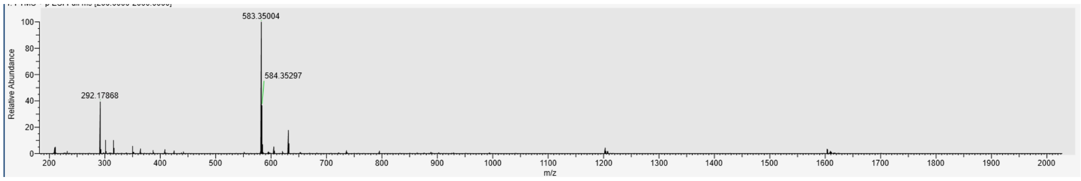

**18** ($^1$H-NMR, 600 MHz, $CD_3OD$):

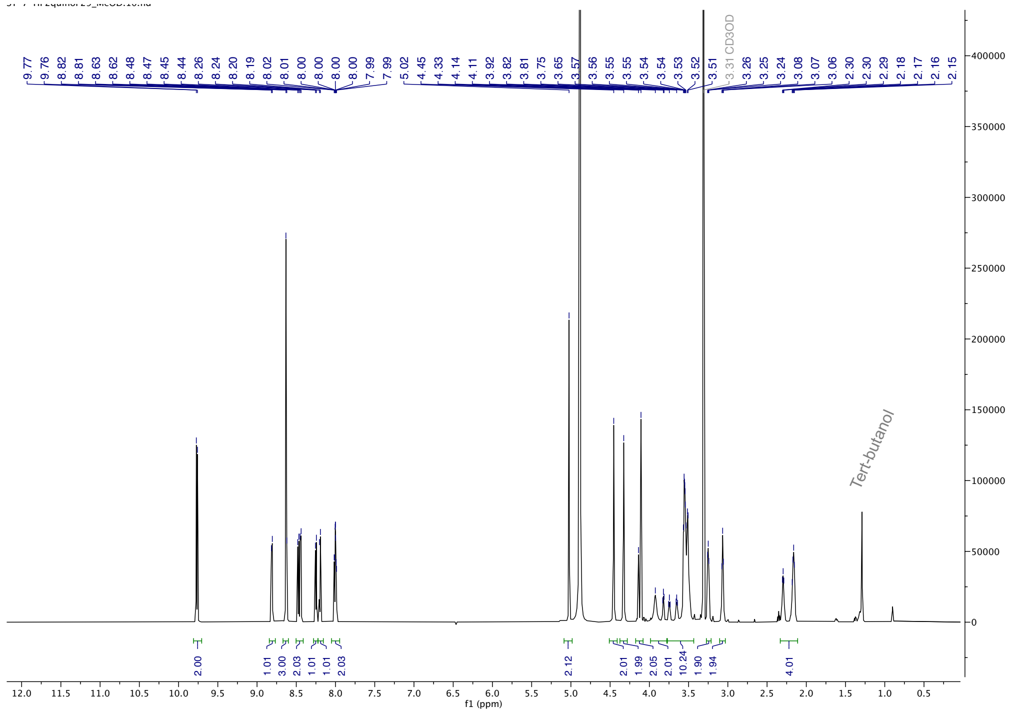

**18** ($^{13}$C-NMR, 150 MHz, $CD_3OD$):

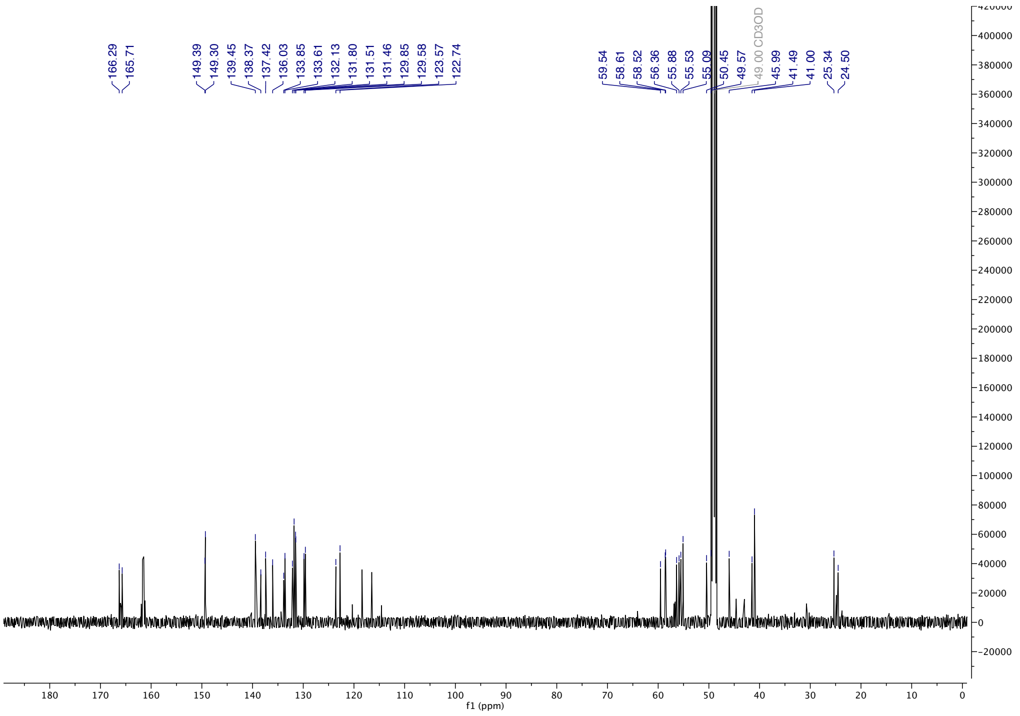

**18** (HPLC, λ = 254 nm, 0-60% MeOH in $H_2O$, +0.1% TFA, 60 min; 60-100% MeOH in $H_2O$, +0.1% TFA, 5 min):

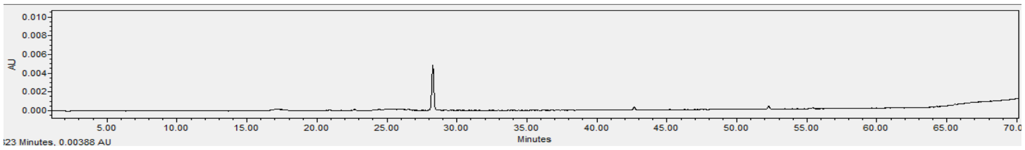


**18** (ESI-HRMS):

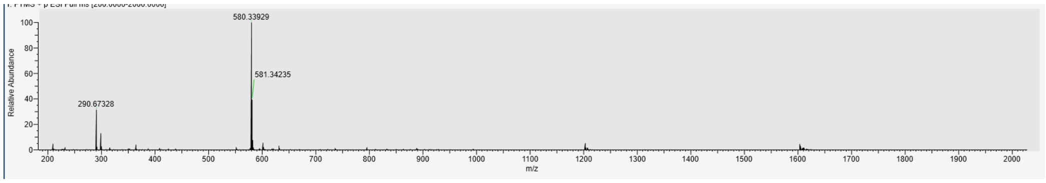

**19** ($^1$H-NMR, 600 MHz, $CD_3OD$):

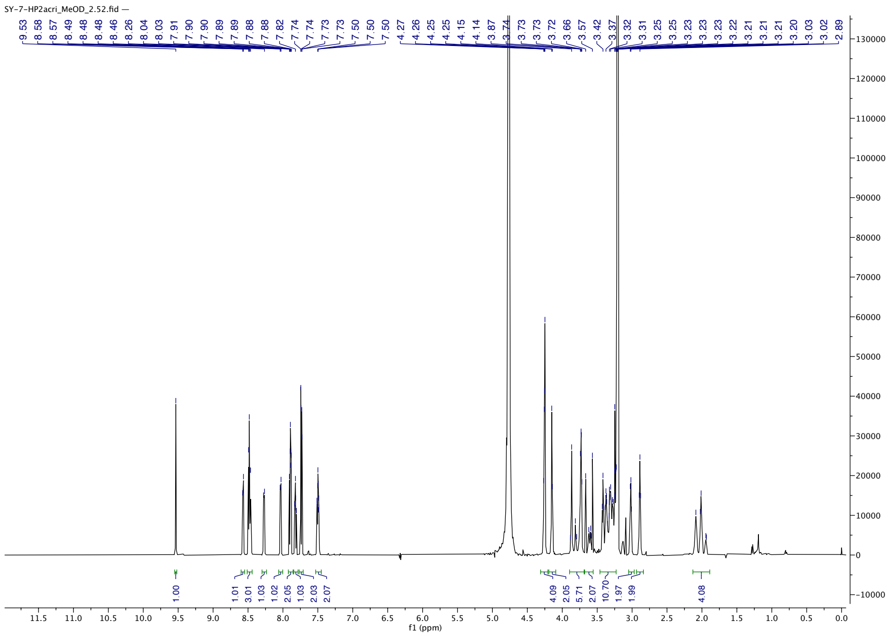

**19** (HPLC, λ = 254 nm, 0-60% MeOH in $H_2O$, +0.1% TFA, 60 min; 60-100% MeOH in $H_2O$, +0.1% TFA, 5 min):

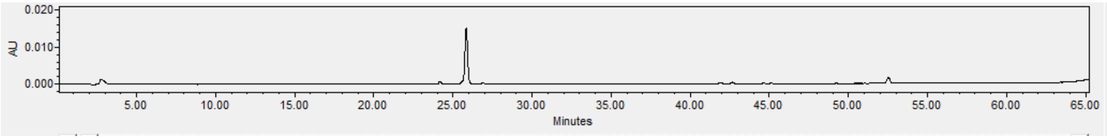


**19** (ESI-HRMS):

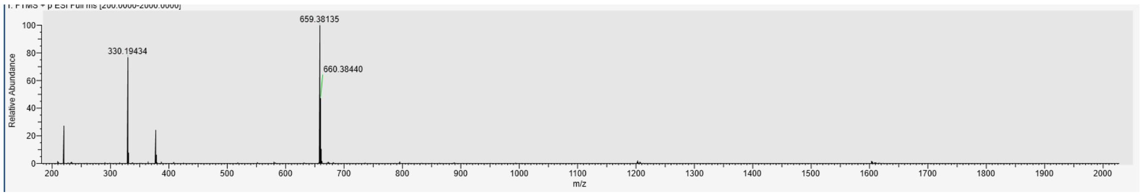
